# *Wwc2* is a novel cell division regulator during preimplantation mouse embryo lineage formation and oogenesis

**DOI:** 10.1101/2019.12.12.872366

**Authors:** Giorgio Virnicchi, Pablo Bora, Lenka Gahurová, Andrej Šušor, Alexander W. Bruce

**Affiliations:** Laboratory of Early Mammalian Developmental Biology (LEMDB), Department of Molecular Biology & Genetics, Faculty of Science, University of South Bohemia, Branišovská 31, 370 05 České Budějovice (Budweis), CZECH REPUBLIC; Laboratory of Biochemistry and Molecular Biology of Germ Cells, Institute of Animal Physiology and Genetics, Czech Academy of Sciences, Rumburská 89, 277 21 Liběchov, CZECH REPUBLIC

## Abstract

Formation of a mature and hatching mouse blastocyst marks the end of the preimplantation development, whereby regulated cell cleavages culminate in the formation of three distinct lineages. We report dysregulated expression of *Wwc2*, an ill-characterised paralog of the Hippo-signalling activator *Kibra/Wwc1*, is specifically associated with cell autonomous deficits in embryo cell number and cell division abnormalities, typified by imbalanced daughter cell chromatin segregation. Division phenotypes are also observed during mouse oocyte meiotic maturation, as *Wwc2* dysregulation blocks progression to the fertilisation competent stage of meiosis II metaphase arrest, characterised by spindle defects and failed Aurora-A kinase (AURKA) activation. Such cell division defects, each occurring in the absence of centrosomes, are fully reversible by expression of recombinant HA-epitope tagged WWC2, restoring activated oocyte AURKA levels. Additionally, clonal dysregulation implicates *Wwc2* in maintaining the pluripotent late blastocyst stage epiblast lineage. Thus, *Wwc2* is a novel regulator of meiotic and early mitotic cell divisions, and mouse blastocyst cell-fate.

## INTRODUCTION

Following the fertilisation of mouse metaphase II arrested secondary (MII) oocytes, a series of asynchronous cleavage divisions generates blastocysts comprising three distinct cell lineages: an outer-residing, differentiating and apical-basolaterally polarised epithelium called the trophectoderm (TE, a progenitor of placental tissue), a further polarised mono-layer of differentiating inner-cell mass (ICM) cells in contact with the fluid filled cavity comprising the primitive endoderm (PrE, progenitors of yolk sac membranes) and pluripotent cells completely encapsulated in the ICM known as the epiblast (EPI, foetal cell progenitors) (see reviews - Frum and Ralston, 2015;Chazaud and Yamanaka, 2016;Rossant, 2016;Mihajlovic and Bruce, 2017;Rossant, 2018;White et al., 2018). Differential regulation of Hippo-signalling (originally identified in *Drosophila* as a cell proliferation and tissue growth regulating pathway, now implicated in varied developmental/pathological paradigms (Davis and Tapon, 2019)) has been identified as an important mechanism of blastocyst lineage specification. Without listing all involved molecular players (see reviews (Hirate et al., 2015;Chazaud and Yamanaka, 2016;Sasaki, 2017)), polarity dependent Hippo-pathway suppression in outer cells enables formation of activating TEAD4 transcriptional complexes (involving nuclear localisation of specific co-factors, YAP and WWTR1/TAZ, collectively referred to here as YAP) to potentiate TE specific gene expression, whereas activated Hippo-signalling in apolar inner cells inhibits this process (via activating LATS1/2 kinases to prevent YAP nuclear localisation)(Nishioka et al., 2009). TEAD4-YAP complexes also simultaneously suppress pluripotent gene expression (*e.g. Sox2*) in derived outer-cells (Wicklow et al., 2014;Frum et al., 2018) and prevent precocious *Sox2* expression prior to the 16-cell stage (Frum et al., 2019). However, eventual EPI specification by the late blastocyst stage, actually requires ICM cell YAP redistribution to the nucleus (implying suppression of Hippo-signalling); an inherently heterogeneous process causing competitive apoptotic elimination of EPI progenitors of reduced naïve pluripotency (Hashimoto and Sasaki, 2019). Collectively, these data illustrate the important and integral nature of Hippo-signalling in regulating key cell-fate events in early mouse embryogenesis and suggests roles for other, particularly upstream and potentially novel, factors capable of functionally interacting with core Hippo-pathway machinery.

The *Drosophila* WW-domain and C2-domain containing (WWC-domain) gene *Kibra* is a positive regulator of Hippo-signalling, causing phosphorylation of the fly orthologue of mammalian LATS1/2 (warts/Wts) in tumour suppressor screens (Baumgartner et al., 2010;Genevet et al., 2010;Yu et al., 2010); a role subsequently confirmed in mammalian cell lines (Xiao et al., 2011a). Unlike *Drosophila*, tetrapod genomes typically encode three WWC-domain paralogs, called *KIBRA/WWC1*, *WWC2* and *WWC3*, although the *Mus musculus* genome does not contain a *Wwc3* gene due to an evolutionarily recent chromosomal deletion. The paralogous human WWC-domain proteins are highly conserved, cable of homo- and hetero-dimerisation, can all activate Hippo-signalling (causing LATS1/2 and YAP phosphorylation) and cause the Hippo-related *Drosophila ‘*rough-eye’ phenotype, caused by reduced cell proliferation, when over-expressed in the developing fly eye (Wennmann et al., 2014). Despite a comparatively large and pan-model KIBRA-related literature, the roles of WWC2/3 are considerably understudied and restricted to limited prognostic reports consistent of tumour suppressor function in specific cancers (*e.g.* hepatocellular carcinoma (Zhang et al., 2017) and epithelial-mesenchymal lung cancers (Han et al., 2018)). There are no reports of any functional roles for WWC-domain containing genes during mammalian preimplantation development.

Ovulated mouse MII oocytes arise from the maturation of subpopulations of meiosis I (MI) prophase arrested primary oocytes, stimulated to re-enter meiosis by maternal reproductive hormones (reviewed (Sanders and Jones, 2018)). Meiotic resumption ensures necessary growth required to support later embryonic development and expression of gene products required to execute meiosis; faithfully segregate homologous chromosome bivalents, form the first polar body and a developmentally competent MII oocyte (itself capable of correctly segregating sister chromatids post-fertilisation). Failed chromosome segregation, resulting in egg and/or zygotic aneuploidy, has usually terminal consequences for embryonic development and aneuploidy attributable to the human female germline is recorded as the leading single cause of spontaneously aborted pregnancy (Hassold and Hunt, 2001;Nagaoka et al., 2012). An extensive literature covering many aspects of the germane segregation of homologous chromosomes during MI exists (see comprehensive reviews (Bennabi et al., 2016;Mihajlovic and FitzHarris, 2018;Mogessie et al., 2018;Namgoong and Kim, 2018;Sanders and Jones, 2018)). As in all mammals, and unlike most mitotic somatic cells, mouse meiotic spindle formation occurs in the absence of centrioles/centrosomes and is initiated around condensed chromosomes from coalescing microtubule organising centres (MTOCs) that are further stabilised by chromosome derived RAN-GTP gradients (Bennabi et al., 2016;Severson et al., 2016;Gruss, 2018;Mogessie et al., 2018;Namgoong and Kim, 2018). Transition from MTOC initiated spindle formation to centrosomal control in mice only occurs by the mid-blastocysts (E4.0) stage, when centrosomes appear *de novo* ((Courtois et al., 2012)). This contrasts with other mammals were fertilising sperm provide a founder centriole that duplicates and ensures the first mitotic spindle is assembled centrosomally (Sathananthan et al., 1991;Schatten and Sun, 2009). Amongst known key regulators of meiotic/mitotic spindle dynamics are the conserved Aurora-kinase family (AURKA, AURKB & AURKC, collectively referred to here as AURKs) that all exhibit germ cell and early embryonic expression (AURKC is not expressed in other somatic cells (Li et al., 2017)). During meiosis, AURKs have important regulatory roles during spindle formation/organisation, MTOC clustering, chromosome condensation and alignment. Moreover, in mitosis, AURKs regulate correct microtubule-kinetochore attachment, chromosomal cohesion and cytokinesis (reviewed in (Nguyen and Schindler, 2017)). Specifically, AURKA protein is essential for MI progression (Saskova et al., 2008) and continually localises with MTOCs and MI/MII spindle poles, regulating MTOC clustering and initiating microtubule nucleation/dynamics, throughout oocyte meiosis (Swain et al., 2008;Solc et al., 2012;Nguyen and Schindler, 2017;Nguyen et al., 2018); with similar roles in post-fertilisation zygotes (Kovarikova et al., 2016).

We report abrogated *Wwc2* expression in mouse embryo blastomeres causes cell autonomous cleavage division phenotypes, yielding blastocysts with fewer overall cells, mis-segregated chromosomes and defective cytokinesis (not replicated by targeted *Kibra* expression). Cell division phenotypes in meiotically maturating mouse oocytes are also caused by RNAi targeted *Wwc2* expression and are coincident with spindle defects and blocked activation of AURKA. Such phenotypes are rescuable by expression of epitope-tagged recombinant WWC2 protein, that in embryos localises with mid-body structures and restores appropriate cellular contribution to the pluripotent EPI, normally lacking after *Wwc2* dysregulation.

## MATERIALS & METHODS

Mouse embryo related experimental procedures were approved and licensed by local ethics committees at the Biology Centre (in České Budějovice) of the Czech Academy of Sciences in accordance with Act No 246/1992 for the protection of animals against cruelty; issued by Ministry of Agriculture of the Czech Republic, 25.09.2014, number CZ02389.

### Embryo and oocyte culture and microinjection

2-cell stage (E1.5) embryo collection, from super-ovulated and mated 10 week old F1 hybrid (C57Bl6 x CBA/W) female mice, and *in vitro* culture in mineral oil covered drops (~20 μL/ ~15 embryos) of commercial KSOM (EmbryoMax® KSOM without Phenol Red; Merck, MR-020P-5F), conducted as previously described (Mihajlovic et al., 2015). Germinal vesicle (GV) stage primary oocytes were mechanically recovered into M2 media (containing 100 μM 3-isobutyl-1-methylxanthine/IBMX; cAMP/cGMP phosphodiesterase inhibitor that maintains GV meiosis I arrest)) from mature Graafian follicles of dissected ovaries of dams after PMSG (pregnant mare serum gonadotrophin) hormone stimulation (7.5U intraperitoneal injection of F1 hybrid strain, 48 hours prior). Recovered oocytes placed into pre-warmed (37°C) drops (~20 μl/ ~15 oocytes) of commercial CZB media (EmbryoMax® CZB media with Phenol Red; Merck, MR-019-D; +IBMX), overlaid with mineral oil and incubated (37°C/ 5% CO_2_) for 2 hours prior to microinjection.

Individual blastomere dsRNA, siRNA or recombinant mRNA microinjections (or combinations thereof), plus post-microinjection culture protocols, performed on 2-cell stage embryos (in either one or both blastomeres) according to defined protocols (Zernicka-Goetz et al., 1996), using apparatus formerly described (Mihajlovic et al., 2015). Recovered GV staged oocytes microinjected using a minimally adapted protocol; namely oocytes were microinjected on 37°C heated stage in concaved glass microscope slides filled with CZB media (+100 μM IBMX), overlaid with mineral oil. Post-microinjection, GV oocytes returned to the incubator for another 18 hours (to permit microinjected siRNA mediated gene knockdown or mRNA expression; CZB +IBMX media) then transferred to pre-equilibrated CZB media drops lacking IBMX, to induce resumption of meiosis I and *in vitro* maturation (IVM). Cultured (and microinjected) embryos/oocytes assayed at various developmental points, as dictated by individual experiments. As indicated, rhodamine-conjugated dextran beads (RDBs; 1 μg/ μl – ThermoFisher Scientific, D1818) were co-microinjected to confirm successful blastomere/oocyte microinjection; a summary of the origin and concentrations of all microinjected RNA species is given in supplementary methods tables SM1. Non-microinjected embryos, or oocytes, (2-3 per microinjection experiment, per plate) served as culture sentinels to confirm successful *in vitro* embryo development/oocyte maturation. In cases of AURKA chemical inhibition (to confirm specificity of anti-p-AURKA antisera), recovered GV oocytes were transferred from IMBX containing CZB media to pre-equilibrated CZB drops lacking IBMX but containing specific AURKA inhibitor, MLN8237 (1μM; Selleckchem, S1133).

### dsRNA and recombinant mRNA synthesis

Specific long dsRNA molecules targeting coding regions of mouse *Kibra* and *Wwc2* derived transcripts (plus negative control GFP) were designed (and *in silico* validated using online E-RNAi resource (Horn and Boutros, 2010)) and synthesised. Briefly, gene specific PCR primer pairs, incorporating 5’-T7-derived RNA polymerase promoters and spanning designed dsRNA complementary sequence, were used to derive *in vitro* transcription (IVT) template, using mouse blastocyst (E3.5) cDNA as a template (or plasmid DNA for GFP). After agarose gel verification, double stranded DNA templates were used in preparatory IVM reactions, incorporating DNaseI and single-stranded RNase treatment (MEGAscript T7; ThermoFisher Scientific, AMB13345), generating dsRNA. The integrity of derived *Kibra*-, *Wwc2*-and *GFP*-dsRNAs was confirmed by non-denaturing gel electrophoresis and quantified (Nanodrop). PCR primer sequences used are provided in supplementary methods table SM2.

Microinjected mRNA constructs derived using commercially available IVT reaction kit (mMACHINE T3; ThermoFisher Scientific, AM1348), using restriction enzyme linearized (using *SfiI*) plasmid DNA as template (2 μg) as follows; i. C-terminal GFP-fusion *Gap43* mRNA plasma membrane marker, from pRN3P-C-term-GFP-Gap43 (Morris et al., 2010)), ii. C-terminal RFP-fusion histone H2B mRNA (from pRN3-C-term-RFP-Hist1h2bb, derived in this study) and iii. siRNA-resistant N-terminal HA-tagged *Wwc2* mRNA (from pRN3P-N-term-HA-siRNAres-Wwc2, derived here). All synthesised mRNAs subject to post-synthesis 3’ poly-adenylation (Poly(A) Tailing Kit - ThermoFisher Scientific, AM1350), confirmed by denaturing gel electrophoresis and quantified (Nanodrop).

### Recombinant plasmid construct generation

Four recombinant plasmids (required for IVT) generated using standard molecular biological protocols. RFP-histone H2B fusion protein (pRN3-C-term-RFP-Hist1h2bb) derived by in frame cloning of a high-fidelity PCR amplified cDNA (Phusion® DNA polymerase, New England BioLabs, M05305) encoding histone H2B (*Hist1h2bb*, lacking endogenous start and stop codons), flanked by oligonucleotide introduced restriction sites (*NheI*), into RNA transcription cassette plasmid pRN3-insert-RFP variant (vector encodes required start and stop codons). Thus, generating required H2B-RFP fusion reporter, downstream of vector encoded T3 RNA polymerase promoter (for IVT) and flanked by UTRs from the frog β-globin gene (Lemaire et al., 1995). The IVT plasmid encoding the siRNA-resistant N-terminal HA-tagged *Wwc2* mRNA (pRN3P-N-term-HA-siRNAres-Wwc2) was created from an annotated full-length Riken mouse *Wwc2* cDNA clone (clone ID: M5C1098O04, Source Bioscience) by deriving high fidelity PCR product, with oligo introduced 5’ (*SpeI*) and 3’ (*NotI*) restriction sites and an in frame N-terminal HA-epitope tag, and ‘TA cloned’ (after addition of 3’ adenine nucleotide overhangs) it into pGEM®-T-Easy plasmid vector (Promega, A1360). The derived plasmid was mutagenised (commercial service; EuroFins) to alter nucleotide (but not redundant amino acid codon) sequence of the siRNA (utilised in this study) recognition motif. The recombinant gene sequence was sub-cloned, using introduced *SpeI* and *NotI* restriction sites, into the multiple cloning site of the RNA transcription cassette plasmid pRN3P (Zernicka-Goetz et al., 1996) to derive IVT competent pRN3P-N-term-HA-siRNAres-Wwc2. All four derived plasmids were sequence verified and the PCR primers (or relevant details) used to generate the required inserts are described in supplementary methods table SM3.

### Q-RT-PCR

Per experimental condition, total RNA was extracted from ~30 (microinjected) mouse embryos (oocytes) *in vitro* cultured to the desired developmental stage, using PicoPure RNA isolation kit (Arcturus Biosciences/ ThermoFisher Scientific, KIT0204), as instructed. Eluted RNA (10 µL) was DNaseI treated (DNA-*free* kit; ThermoFisher Scientific, AM1906) and used to derive cDNA (30 µL), either via oligo-dT (embryos and oocyte) or random hexamer (oocytes only) priming (Superscript-III Reverse Transcriptase; ThermoFisher Scientific, 18080085). 0.5 µl of diluted template cDNA (1:3, nuclease-free water) per real-time PCR reaction (10 µl – SYBR Green PCR kit, Qiagen, 204143) was used to assay specific transcript abundance (CFX96 Real-Time System, BioRad). *Kibra* and *Wwc2* transcript levels were internally normalised against those for *Tbp* (TATA-binding protein) housekeeping gene and fold changes (±s.e.m.) derived using the ΔΔCt method (Livak and Schmittgen, 2001). A minimum of two biological replicates of at least three technical replicates were employed; specific gene oligo primer sequences (final reaction conc. 400 nM) are in supplementary methods table SM4.

### Immuno-fluorescent staining and confocal microscopy imaging

*In vitro* cultured (microinjected) embryos/oocytes were fixed (at required developmental stages) with 4% para-formaldehyde, immuno-fluorescently (IF) stained and imaged in complete z-series by confocal microscopy (using FV10i confocal microscope, Olympus) as described (Mihajlovic et al., 2015); supplementary methods table SM5 summarises the identity and combinations (plus employed concentrations) of primary and fluorescently conjugated secondary antibodies used. The majority of IF stained embryos/oocytes were counterstained for DNA, using DAPI containing mounting solution (Vectashield plus DAPI, Vector Labs) and as indicated some embryo samples were also counterstained against filamentous-actin, using fluorescently-labelled rhodamine-conjugated phalloidin (ThermoFisher, R415 - described in (Mihajlovic and Bruce, 2016)).

### Embryo/oocyte image analysis/ cell counting

Contributions of each individual cell, per fixed embryo sample (in control or *Kibra/Wwc2* gene RNAi KD conditions), to; inner (entirely encapsulated) outer (whereby cells retain cell contactless domains) embryo compartments, emerging blastocyst cell lineages (defined by presence and/or absence of specific and stated lineage marker protein expression), cell populations defined by presence of a defining intra-cellular feature (*e.g.* cell/nuclei morphological abnormalities, apoptosis, lineage marker protein or mid-bodies) or presence within or outwith a microinjected clone (distinguishable by a fluorescent injection marker signal, *i.e.* RDBs or recombinant fluorescent fusion proteins) was determined by serial inspection of individual confocal micrograph z-sections of each embryo, using commercial Fluoview ver.1.7.a (Olympus), Imaris (Bitplane) and ImageJ software. These calculated contributions were individually tabulated (see supplementary tables) and the mean of cells within defined sub-populations, plus standard error of means (mean ± s.e.m.) determined. The statistical significance between relevant experimental and control groups was determined using 2-tailed Student’s t-tests (experiment specific supplementary tables provide comprehensive summaries, plus all individual embryo data). Relating to oocytes, a similar approach was used to assay mean frequencies by which maturing oocytes exhibit intra-cellular morphologies (revealed by combined IF staining -*i.e.* α-TUBULIN and p-AURKA and microinjected fluorescent reporter protein expression - *i.e.* RFP-histone H2B) indicative of appropriate or aberrant oocyte maturation (as categorised), at the indicated developmental time-point; such data are summarised in relevant supplementary tables, including individual oocyte derived data.

### Western blotting

Precise numbers of oocytes per comparable experimental condition (~15-30) were washed in phosphate buffer saline (PBS; Merck, P5493) containing polyvinyl alcohol (PVA; Merck, 341584), frozen in a residual volume at −80°C, before lysis, in 10 μl of 10x SDS reducing agent/loading buffer (NuPAGE buffer, ThermoFisher Scientific, NP 0004, ThermoFisher Scientific), by boiling at 100°C for 5 minutes. Loaded proteins were electrophoretically separated on gradient precast 4–12% SDS–PAGE gels (ThermoFisher Scientific, NP0323) and transferred to Immobilon P membranes (Merck group, IVPD00010) using a semi-dry blotting system (Biometra/ Analytik Jena) for 25 minutes at 5 mA/cm^2^. Blotted membranes were blocked in 5% skimmed milk powder dissolved in 0.05% Tween-Tris pH 7.4 buffered saline (TTBS) for 1 hour, briefly rinsed in TTBS and then incubated overnight at 4°C in 1% milk/TTBS containing primary antibody. Membranes were washed in three changes of TTBS buffer (20 minutes each at room temperature) and horse-radish peroxidase conjugated secondary antibody added to the blot in 1% milk/TTBS, for 1 hour (room temperature). Immuno-detected proteins were visualized by chemiluminescent photographic film exposure (ECL kit; GE Life Sciences, RPN2232) and digitally scanned using a GS-800 calibrated densitometer (Bio-Rad Laboratories). Antibody stripped membrane blots were re-probed, for loading controls, in an identical manner. Supplementary methods tables SM6 details the utilised concentrations of the primary and peroxidase-conjugated secondary antibodies used.

### Supplementary materials

Three components: a) Supplementary Figures S1-S10, plus legends, b) Supplementary Methods Tables SM1 — SM6 and c) Supplementary Tables ST1 – ST26 (detailing all analysed embryo/oocyte cell counts and statistics).

## RESULTS

### WWC-domain gene expression in preimplantation mouse embryos

In the central context of Hippo-signalling during early mouse development (Sasaki, 2017), we investigated the potential role of WWC-domain containing genes (*Kibra* and *Wwc2* (Wennmann et al., 2014)) during blastocyst formation. We assayed *Kibra* and *Wwc2* mRNA expression levels after microinjection of control dsRNA or constructs specific for each gene (Fig. 1a). Stable levels of *Kibra* and *Wwc2* transcripts were readily detectable at both 8-(E2.5) and 32-cell (E3.5) stages, with normalised *Wwc2* expression twice as abundant as *Kibra*. Each transcript could be robustly depleted using dsRNA mediated global knockdown (KD) at both assayed stages, although we could not assay protein expression due to a lack of reliable anti-sera. We next assayed for overt developmental defects associated with individual or combined *Kibra*/*Wwc2* KD by microinjecting dsRNA(s) in one blastomere of 2-cell (E1.5) stage embryos, culturing to the late blastocyst (E4.5) stage and counting total cell number (Fig. 1b). *Kibra* gene KD had no significant effect on total cell number (versus control dsRNA) but a severe attenuation when targeting *Kibra* and *Wwc2* transcripts together was observed. This defect was statistically indistinguishable from embryo groups microinjected with *Wwc2* dsRNA alone, indicating sole *Wwc2* KD was sufficient to induce the observed phenotype (note, *Kibra* mRNA levels were unaffected by *Wwc2* dsRNA at the 8-cell stage, whereas *Wwc2* transcripts were robustly reduced, confirming *Wwc2* dsRNA specificity – Fig. 1a and supplementary Fig. S1). Repetition of *Wwc2* KD, in a fluorescently marked clone (rhodamine conjugated dextran beads/ RDBs were co-microinjected with *Wwc2* dsRNA) confirmed reduced cell numbers, as early as the 16-cell (E3.0) stage, specifically within the marked clone that itself had reduced contribution to the initial inner-cell/ICM founding population (Fig. 1c). Accordingly, we focussed further investigations on the overt cell number deficit caused by *Wwc2* KD.

**Figure 1:**
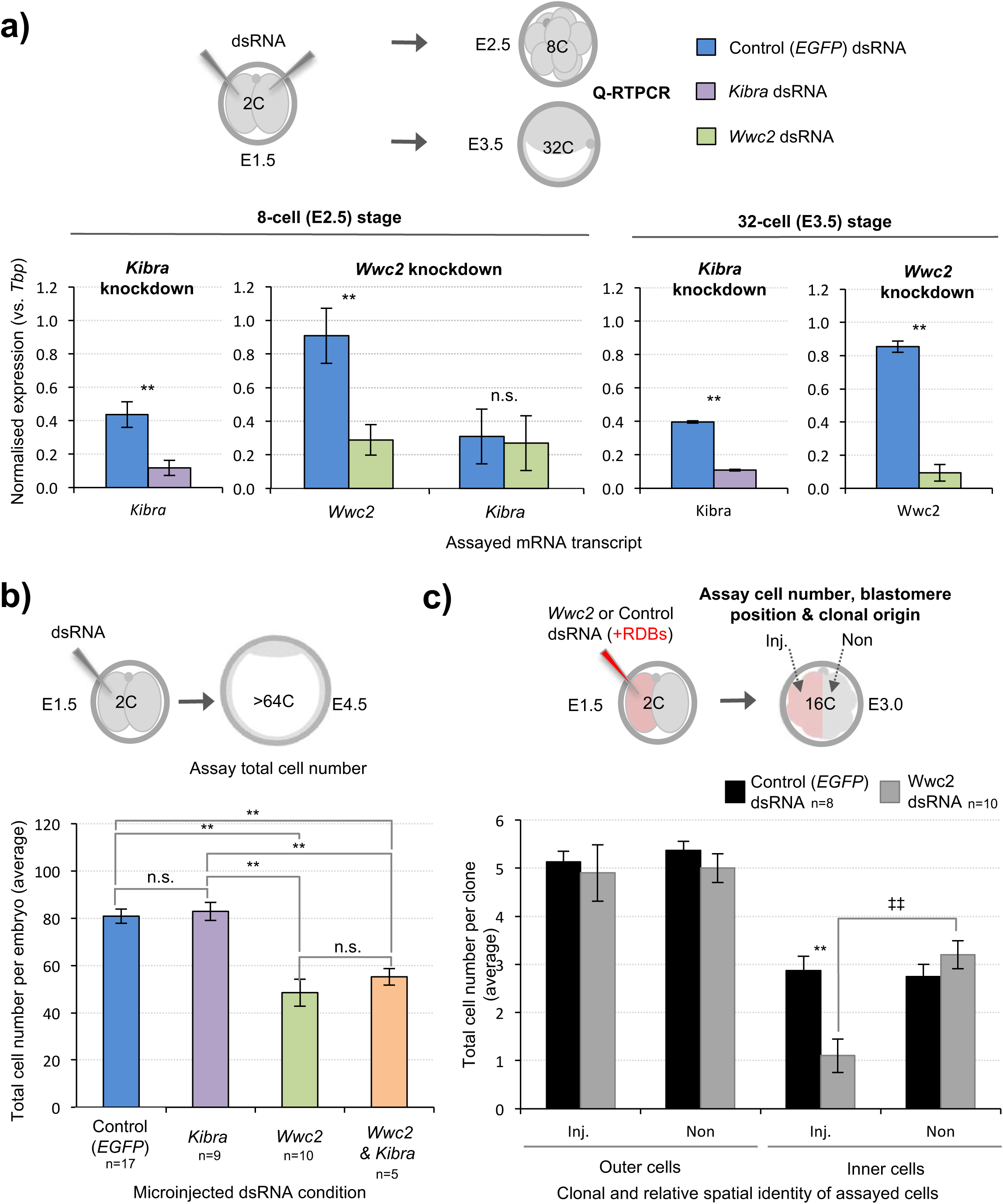
dsRNA mediated WWC-domain containing gene (*Kibra* & *Wwc2*) KD in preimplantation mouse embryos is associated (*Wwc2*) reduced cell number. **a)** Experimental schema of *Kibra* and *Wwc2* specific dsRNA (plus *EGFP n*egative control) microinjection mediated global embryo KD, followed by culture to either 8-cell or 32-cell stages and Q-RT-PCR analysis (upper). Normalised expression of either *Kibra* or *Wwc2* derived mRNAs after stated dsRNA microinjection, at stated developmental stages (lower; error bars represent s.e.m., n=3 and ** denotes p<0.005 in 2-tailed student t-test, n.s. denotes lack of significance). **b)** Experimental schema describing clonal *Kibra*, *Wwc2* or *Kibra*+*Wwc2* KD (microinjection of one cell at the 2-cell stage) and total fixed embryo cell number assay at the late blastocyst (>64-cell) stage (upper). Average total cell number per embryo in each stated dsRNA microinjection group (lower chart). **c)** Experimental strategy to KD *Wwc2* expression (as in b) in marked clones, (*Wwc2* dsRNA and rhodamine conjugated dextran microbeads/RDBs co-microinjection) and assay clonal contribution at the 16-cell stage (upper). Average total cell number per clone (Inj. vs. Non) in *Wwc2* or control dsRNA microinjection groups, as determined by relative position of individual cells within the embryo (encapsulated inside or with a cell contactless domain on outside; lower chart). In panels b) & c) chart error bars represent s.e.m. with indicated ‘n’ numbers. Highlighted statistically significant differences (2-tailed students t-test) between experimental microinjection groups (asterisks), or clones (Inj. vs. Non) within a group (double crosses), with statistical confidence intervals of p<0.05 and p<0.005, denoted by one or two significance markers, respectively; in panel b) a lack of significant difference between compared groups is marked ‘n.s.’. Supplementary tables ST1 and ST2 summarise statistical analysis and individual embryo data.

### Reduced embryo cell numbers caused by *Wwc2* KD are associated with defective cell division

Although the utilised *Wwc2* dsRNA was carefully designed to avoid off target effects (specifically relating to *Kibra* expression, Figs. 1a & S1), we sought to make a phenocopy using an siRNA mediated approach (targeting an alternative *Wwc2* mRNA region – Fig. S1). In addition to confirming the phenotype, the switch was adopted to permit phenotypic rescue experiments, by expressing recombinant siRNA-resistant *Wwc2* mRNA (see below). The *Wwc2* siRNA near completely eliminated detectable *Wwc2* transcripts in global KD (Fig. 2b). A subsequent assay of total cell number at all cleavage stages, revealed an accumulative and statistically robust deficit beginning after the 8-cell stage (Fig. 2c). Indeed, from the equivalent 8-to 32-cell stages (relative to control embryo development), total cell number did not significantly increase in *Wwc2* KD embryos but did start to increase during the blastocyst maturation period (32-to >64-cell stages). Next, we assayed if *Wwc2* siRNA phenotypes were cell autonomous by creating RDB marked *Wwc2* specific siRNA KD clones (comprising 50% of the embryo – Fig. S2). We observed a statistically significant *Wwc2* siRNA mediated deficit in cell number within the microinjected clone (versus both the non-microinjected sister clones and the equivalent microinjected clones in control siRNA embryos), at all post-8-cell stages. We did not observe significant differences between control and *Wwc2* KD groups in the number of cells within the non-microinjected clone (with the exception of the 32-cell stage where there was an average of 2.3 fewer cells in the *Wwc2* KD group). These data support a novel and cell autonomous role for *Wwc2* in ensuring appropriate cell number during preimplantation mouse embryo development.

**Figure 2:**
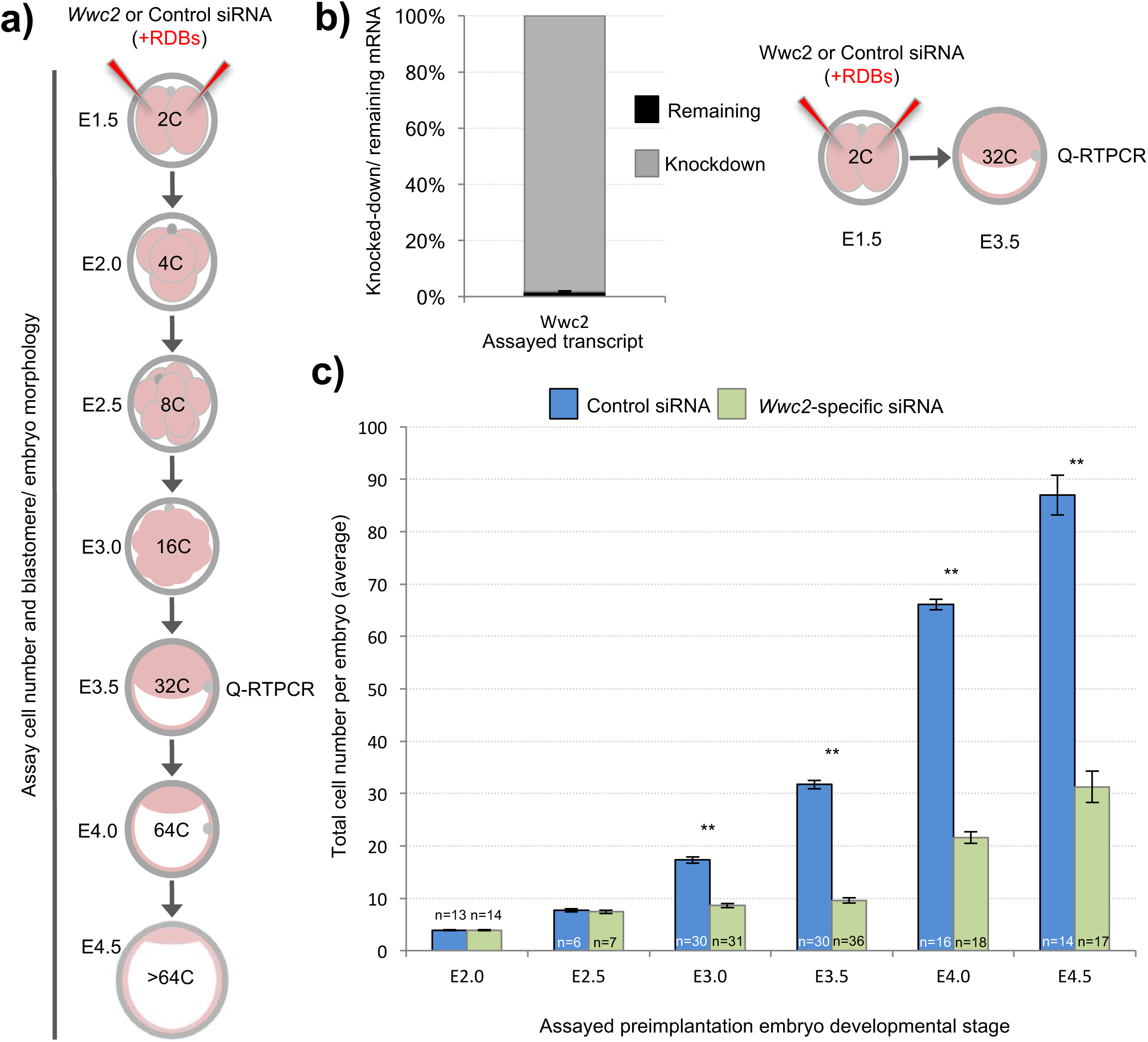
siRNA mediated global *Wwc2* KD causes persistent preimplantation stage cell number deficits from the 8-cell stage. **a)** Experimental design; non-specific control or *Wwc2*-specific siRNA co-microinjected with RDBs, into both blastomeres at the 2-cell stage, and cultured, fixed and subject to total cell assay count (including DAPI and phalloidin staining) at indicated developmental stages. A separate group of microinjected embryos were subject to normalised *Wwc2*-specific Q-RTPCR to assay KD efficiency at the 32-cell stage. **b)** Q-RTPCR data of siRNA mediated endogenous *Wwc2* expression KD at the 32-cellstage (error bars denote s.e.m. of triplicate measurements, n=3). **c)** Average total cell number in control (blue bars) or *Wwc2*-specific (green bars) siRNA microinjected embryos indicated developmental stages (number of embryos in each group is indicated). Errors represent s.e.m. and 2-tailed student t-test derived statistical significance between control and *Wwc2* KD groups indicated (** p<0.005). Supplementary tables ST3-ST8 summarise statistical analysis and individual embryo data. Note, additional data describing frequencies of observable cell morphological/division defects, obtained from this same dataset are presented in Fig. 3.

We also observed a number of morphological nuclear abnormalities within *Wwc2* KD embryos and cell clones, not evident in the controls. Illustrative examples, at the 32-cell equivalent stage are provided (Fig. 3b) and were categorised as; ‘abnormal nuclear morphology’ (including that typical of ‘association with the mid-body’), associated with ‘cytokinesis defects’ (typified by bi-nucleated cells) or coincident with ‘multiple or micronuclei’. We calculated the frequencies by which each or a composite of such abnormalities were observed at each assayed cleavage stage; defined per embryo (in at least one blastomere) or per individual assayed cell (Fig. 3c). Apart from a single incidence of an early blastocyst stage cell exhibiting a single micronucleus, no abnormalities were observed in any control siRNA microinjected embryo, at any developmental stage. Conversely, at the 8-cell stage abnormally shaped nuclei were present in over a quarter of *Wwc2* global KD embryos and in all assayed embryos by the equivalent late blastocyst stage. A similar trend assaying multiple/micronuclei was observed from the 16-cell stage. Although less prevalent, cytokinetic defects also affected nearly a quarter of *Wwc2* KD embryos at the late blastocyst stage. Indeed, all *Wwc2* global KD embryos exhibited one or more defect, in at least one cell, by the mid-blastocyst stage, with the collective defects (excluding cytokinesis/bi-nucleation) first arising in just over a quarter of 8-cell stage embryos. An assay of the same phenotypes at the level of each individually assayed blastomere showed that just over a quarter of cells were affected by the late blastocyst stage, although such embryos comprised a much smaller average of overall cells, versus control siRNA groups (*i.e.* 31.3±3.0 against 87.0±3.8 -Fig. 2c). Thus, cleavage stage *Wwc2* KD is associated with cell autonomous division defects that contribute to embryos with progressively fewer constituent blastomeres from the 8-cell stage and invokes a role for *Wwc2* in regulating appropriate cell division in the early mouse embryo.

**Figure 3:**
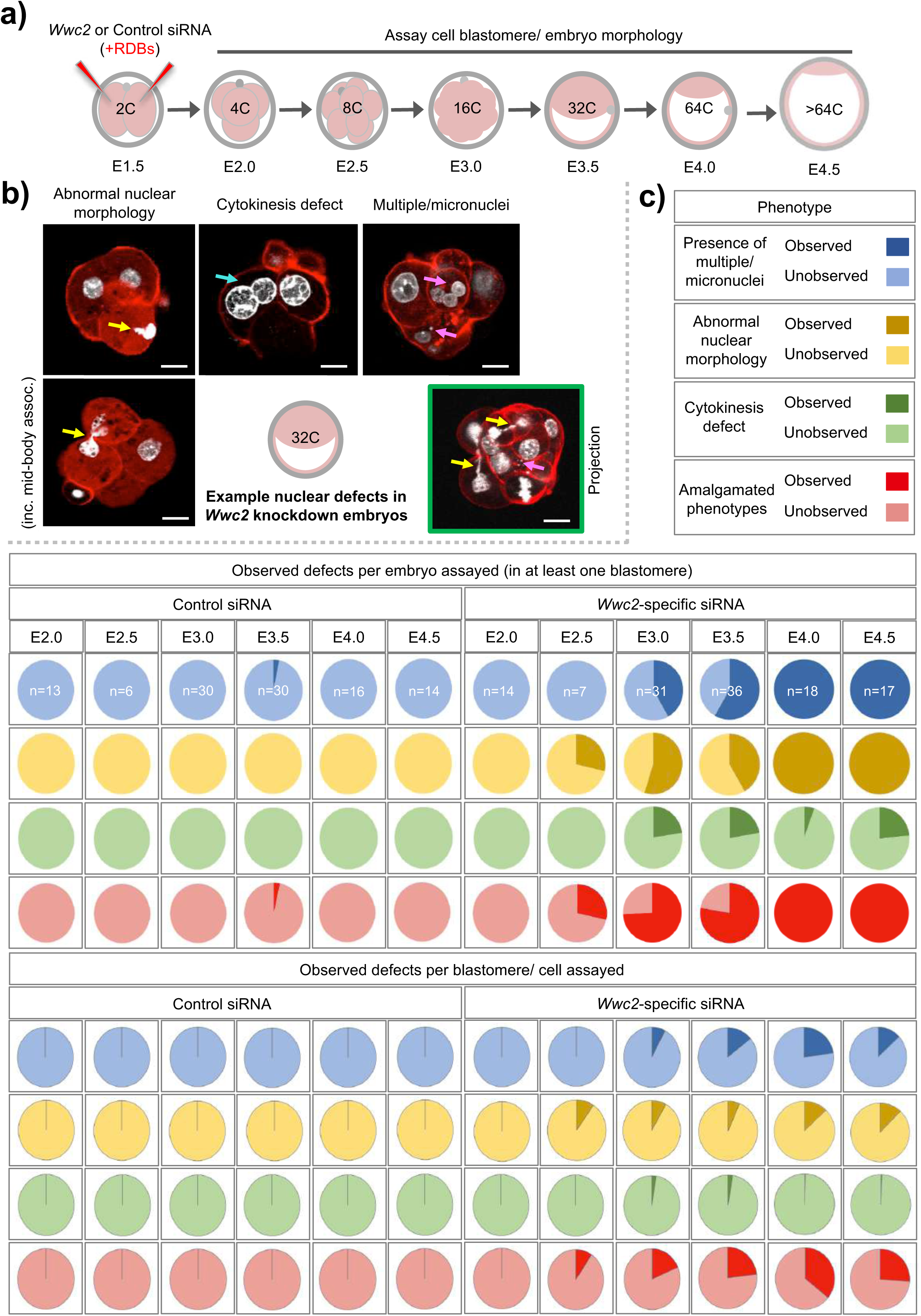
Global *Wwc2* KD mediated cell numbers deficits are associated with defective cell division morphologies. **a)** Schematic of experimental design; 2-cell stage embryos microinjected with control or *Wwc2*-specific siRNA, cultured to indicated developmental stages, fixed and assayed for total cell number or the incidence of abnormal cell morphology indicative of defective cell division (determined by DAPI and rhodamine conjugated phalloidin staining - note, data derived from same experiments as those described in Fig. 2). **b)** Exemplar confocal micrograph z-sections of *Wwc2*-specific siRNA microinjected embryos at the 32-cell stage, illustrating three distinct phenotypic morphological defect categories observed; i. abnormal nuclear morphology (including chromatin mid-body association, highlighted by yellow arrows - left), ii. cytokinesis defects defined by two nuclei per cell (highlighted by blue arrow -centre), iii. presence of multiple (*i.e.* >3, highlighted by pink arrows) nuclei and/or micronuclei per cell (right) and iv. composite of categorised defects highlighted by colour coded arrows in a single embryo confocal z-section projection (green boarder). DAPI (white) and cortical F-actin (red); scale bar = 20 μm. **c)** Frequencies of each observed category of phenotypic/ morphological defect (*i.e.* micronuclei; blue, abnormal nuclei; yellow and failed cytokinesis; green) or a combination of at least two or more (red) in control siRNA (left) and *Wwc2*-specific siRNA (right) microinjected embryos, at indicated developmental stages (numbers of embryos in each group highlighted); data demonstrate defect incidence per embryo (in at least one constituent blastomere – upper row of charts) or on a per assayed cell basis (lower row of charts). Supplementary tables ST3-ST8 summarise statistical analysis and individual embryo data in each experimental group.

### *Wwc2* mRNA depletion in primary (GV) oocytes impairs meiotic maturation

We next addressed whether similar *Wwc2* KD in germinal vesicle (GV) oocytes could cause subsequent meiotic division defects, particularly as both embryo cleavage divisions and meiotic oocyte maturation occur in the absence of centrosomes in mice (Courtois et al., 2012;Coelho et al., 2013;Bennabi et al., 2016;Severson et al., 2016;Gruss, 2018;Mogessie et al., 2018;Namgoong and Kim, 2018). We first confirmed by Q-RT-PCR abundant *Wwc2* transcript expression in GV and MII oocytes (plus zygotes). Exploiting alternative, oligo-dT versus random hexamer, cDNA synthesis priming strategies we uncovered evidence of cytoplasmic *Wwc2* mRNA poly-adenylation, usually associated with increased translational efficiency of functionally important proteins during meiotic maturation (Salles et al., 1992) - Fig. 4a. Consultation of our previously published assay of polysome associated transcripts during mid meiotic maturation supported this interpretation, reporting ~40% polysome association of detectable *Wwc2* transcripts (Koncicka et al., 2018). We then confirmed our siRNA construct could elicit robust *Wwc2* transcript knockdown in microinjected GV oocytes, that had been blocked from re-entering meiosis I (using a cAMP/cGMP phosphodiesterase inhibitor, 3-isobutyl-1-methylxanthine/ IMBX) and then permitted similarly treated oocytes to *in vitro* mature (IVM) to the MII equivalent stage (Fig. 4b & c). Such oocytes were fixed and immuno-fluorescently (IF) stained for α-Tubulin (plus DAPI DNA stain) and assayed by confocal microscopy for potential meiotic maturation/division phenotypes. In the two control conditions (non-microinjected or control siRNA microinjected oocytes) >90% of GV oocytes matured to the MII arrested stage, typified by extruded polar bodies (PB1) and metaphase II arrested spindles. However, in the *Wwc2* KD group, this successful IVM rate was reduced to 8.6%. The remaining oocytes presented with various MI arrested phenotypes (all lacking PB1) categorised by; presence of MI metaphase spindles (-PB1 +MI spindle; 42.9%, note spindles typically failed to migrate to oocyte cortex), spindle-like structures with mis-aligned or dispersed chromosomes (-PB1 +spindle defect; 22.9%) or non-nuclear membrane enveloped chromatin devoid of associated α-Tubulin (-PB1+ultra-condensed chromatin; 25.7%). Such phenotypes collectively indicate profoundly defective oocyte meiotic maturation and impaired homologous chromosome segregation caused by *Wwc2* KD (Fig. 4d & d’). Although assayed oocytes were subject to IVM for 18 hours, we wanted to exclude the possibly the results were indicative of delayed rather than blocked meiotic progression. Consistently, experimental repetition using an extended IVM period of 24 hours did not reveal any increased frequency of MII arrested oocyte generation. Indeed, *Wwc2* KD oocytes either remained blocked in MI or unsuccessfully attempted chromosome segregation without undergoing cytokinesis (Fig. S3).

**Figure 4:**
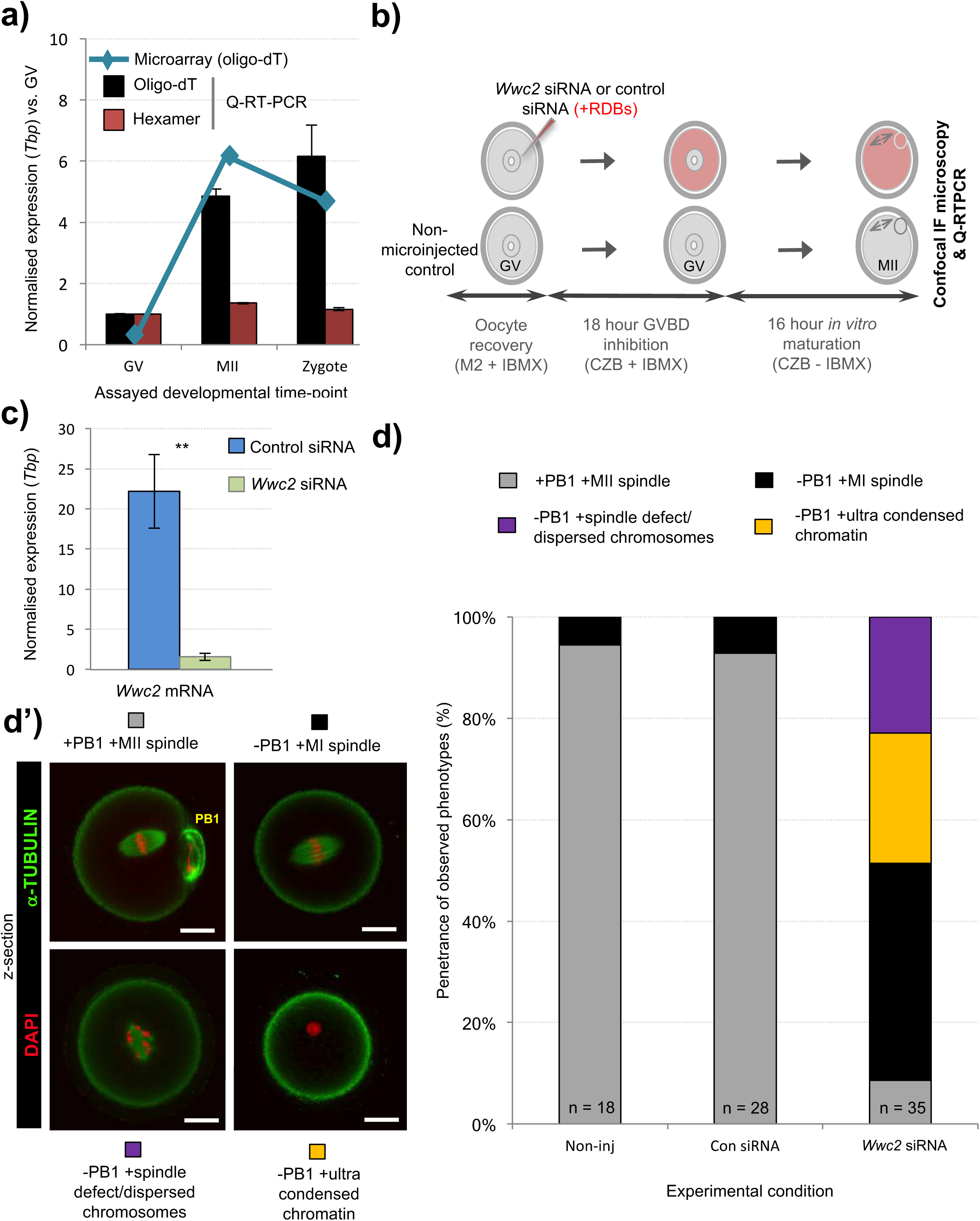
*Wwc2* mRNA is required for mouse oocyte meiotic maturation. **a)** Published microarray (Wang et al., 2004, blue line) and generated Q-RT-PCR (bar charts) normalised *Wwc2* mRNA expression in germinal vesicle and metaphase II arrested oocytes (GV and MII) and fertilised zygotes (Tbp normalised expression, relative to GV stage). Note, microarray data derived from oligo-dT primed reverse transcription and Q-RT-PCR from both oligo-dT (black bars) and random hexamer priming (red bars); error bars represent s.e.m. and n=3. **b)** Experimental schema of *Wwc2* KD in GV oocytes by microinjected *Wwc2* siRNA (plus control siRNA and non-microinjected controls), 18 hour incubation in IMBX containing media (prevents GVBD), 16 hour *in vitro* oocyte maturation (IVM – minus IMBX) and confocal microscopy or Q-RT-PCR analysis; co-microinjected RDBs used as injection control marker. **c)** Q-RT-PCR data of normalised (against *Tbp*) *Wwc2* mRNA levels in control and *Wwc2* siRNA microinjected GV oocytes after IVM to MII stage (error bars represent s.e.m., n=3 and ** denotes p<0.005 in 2-tailed student t-test). **d)** Charts (left) detailing successful IVM frequencies of control non-microinjected (Non-Inj), microinjected control siRNA (Con. siRNA) and microinjected *Wwc2* siRNA GV oocytes, to the MII stage or preceding phenotypic stages. **d’)** examples of successfully matured MII oocytes (note, extruded first polar body/PB1) and quantified categorised phenotypes are shown as confocal z-section micrographs (right – α-Tubulin in green and DAPI DNA pseudo-coloured red; scale bar = 20 μm). Supplementary tables ST14 summarise statistical analysis and individual oocyte data used to generate the figure.

### *Wwc2* KD oocyte phenotypes are associated with failed Aurora kinase-A (AURKA) activation

Phosphorylation dependent activation of Aurora kinase-A (p-AURKA: p-Thr288) is required to initiate acentrosomal and MTOC mediated spindle assembly in mouse oocytes and *Aurka* KD phenotypes closely resemble the *Wwc2* KD defects we describe (Bury et al., 2017). We hypothesised the observed *Wwc2* specific meiotic maturation phenotypes may be associated with impaired p-AURKA activation. Western blot analyses of staged maturing oocytes confirmed characteristically low p-AURKA levels in GV oocytes, that markedly increased in the presence of the MI spindle and were maintained when the MII spindle was formed (Fig. 5b (Saskova et al., 2008;Nguyen and Schindler, 2017)); a trend replicated in control siRNA microinjected GV and resulting MII oocytes (note, anti-p-AURKA antibody specificity was independently confirmed; Fig. S4). However, p-AURKA was not detectable at either the equivalent GV or MII stages in *Wwc2* siRNA microinjected oocytes. Moreover, p-AURKA immuno-reactivity (assayed by confocal microscopy based IF) that would ordinarily localise to spindle poles or spindle-like structures, was also lacking in *Wwc2* KD oocytes. This a stark contrast to that observed on MII arrested spindles in control oocytes (Fig. 5c – note, the p-AURKA channel of the example *Wwc2* KD oocyte is over-exposed to illustrate the lack of spindle associated immuno-reactivity but accentuates non-specific cortical auto-fluorescence also seen in control oocytes). These results confirm failed p-AURKA activation during oocyte maturation, under conditions of experimentally induced *Wwc2* KD, associated with profoundly defective meiotic cell division.

**Figure 5:**
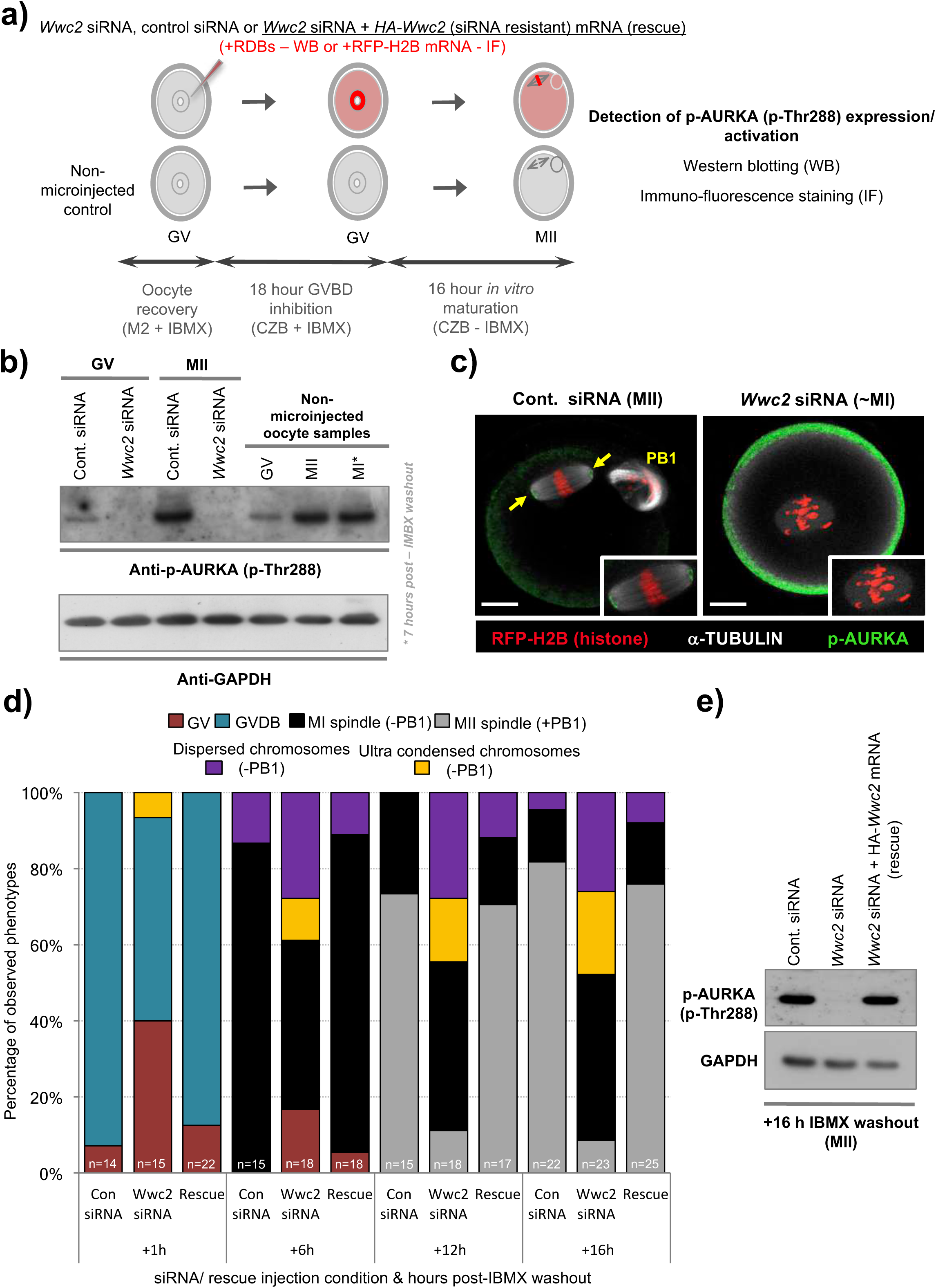
*Wwc2* KD induced oocyte IVM phenotypes are associated with failed Aurora-A (AURKA) phosphorylation/activation _(both rescuable by co-microinjection of siRNA resistant *HA-Wwc2* mRNA). **a)** Experimental schema of GV oocyte microinjection conditions; *i.e. Wwc2* siRNA or control siRNA alone or *Wwc2* siRNA + *HA-Wwc2* (siRNA resistant) mRNA (*i.e. ‘*rescue’), co-microinjected with RDBs (for western blot analysis) or RFP-H2B mRNA (IF – fluorescent marker). Microinjected (plus non-microinjected control) oocytes incubated in IMBX containing media (18 hours - to prevent GVBD), subject to IVM (max. 16 hours – media minus IMBX) and processed for western blotting and IF at designated oocyte maturation time-points (assaying phospho-Aurora/ p-AURKA levels) or assayed for developmental progression/*Wwc2* KD induced phenotypes. **b)** Western blots of activated p-AURKA levels (plus GAPDH housekeeping control), at indicated stages of oocyte maturation (note, GV; +0 hours relative to IMBX washout, metaphase of meiosis I/ MI; +7 hours and MII; +16 hours), in either control or *Wwc2* siRNA microinjected, or non-microinjected conditions. **c)** Exemplar single confocal z-section IF micrographs of control (left) or *Wwc2* siRNA (right) IVM cultured microinjected oocytes at the MII equivalent stage (16 hours post-IBMX) stained for p-AURKA (green) and α-TUBULIN (white) and labelled with RFP-H2B chromatin reporter (red); insets show zoomed region of meiotic spindle. Note, *Wwc2* siRNA microinjection group example image is over-saturated compared to control (to illustrate lack of spindle associated pAURKA – cortical signal is auto-fluorescence); PB1 (control siRNA) denotes extruded first polar body and the scale bars represents 20 μm. **d)** Chart detailing IVM rates of control siRNA, *Wwc2* siRNA and *Wwc2* siRNA + *HA-Wwc2* mRNA (rescue) microinjected oocytes at staged time-points post IBMX removal; as described by percentage of oocytes at any of stated IVM stages or categorised phenotypes (typically associated with *Wwc2* KD – see Fig. 4); number of oocytes in each experimental condition at each assayed IVM stage is indicated. **e)** Western blot of activated p-AURKA levels (plus GAPDH housekeeping control) at MII equivalent stage (16 hours post-IVM), in control siRNA, *Wwc2* siRNA or *Wwc2* siRNA + *HA-Wwc2* mRNA microinjected (rescue) oocytes. Supplementary tables S16-19 summarise statistical analysis and individual oocyte data used to generate the figure.

Despite consistent *Wwc2*-specifc RNAi cell division phenotypes uncovered with distinct dsRNA/siRNA constructs in preimplantation mouse embryos/oocytes (plus optimised design to avoid potential off-target effects or cross reactivity with paralogous *Kibra* mRNA - Figs. 1 and S1), we sought further verification of the identified *Wwc2* specific oocyte role. Accordingly, we derived a recombinant and N-terminally HA-epitope tagged *Wwc2* mRNA construct (*HA-Wwc2*), specifically mutated within the siRNA complementary sequence (yet preserving, via redundancy, the sequence of amino acid codons; Fig. S5), that could remain available for translation in *Wwc2* siRNA microinjected oocytes. Repeating the described IVM experiments (but assaying at multiple time-points), we found that co-microinjection of *HA-Wwc2* mRNA and *Wwc2* siRNA caused a robust rescue of the maturation phenotypes previously observed by *Wwc2* siRNA microinjection alone (MII arrested oocytes +PB1, 16 hours post-IBMX; control siRNA – 81.8%, *Wwc2* siRNA - 8.7% & *HA-Wwc2* mRNA plus *Wwc2* siRNA – 76.0%). Moreover, the progression of the observed rescue through recognisable oocyte maturation stages was in-step with that of control siRNA microinjected oocytes (as measured at 1, 6, 12 and 16 hours post-IBMX; Fig. 5d). We also found *HA-Wwc2* mRNA co-microinjection was associated with reappearance of detectable p-AURKA protein expression (Fig. 5e), and sub-cellular localisation with forming or matured MI/MII stage meiotic spindles (Fig. S6). Collectively, these data confirm a novel *Wwc2* role in regulating appropriate chromosome segregation and cell division during mouse oocyte maturation (associated with activation of the key spindle-associated regulatory kinase AURKA); displaying functional conservation with that observed in similarly acentrosomal mitotic cleavage divisions of preimplantation stage embryo blastomeres.

### siRNA mediated *Wwc2* KD affects blastocyst cell-fate

Embryo cell number deficits and defective division phenotypes were first evident after global (siRNA) *Wwc2* KD around the 8-to 16-cell stage (Figs. 2 & 3), with clonal *Wwc2* KD (dsRNA) being associated with reduced inner-cell clone numbers (Fig. 1c). Transition from the 8- to 16-cell stage represents the first developmental point embryo blastomeres are overtly distinct in terms of intra-cellular apical-basolateral polarity and relative spatial positioning, with consequences for ultimate blastocyst cell-fate (*i.e.* polarised outer-cells that can give rise to both TE and further inner-cells, plus apolar inner ICM progenitors (Johnson and Ziomek, 1981)).Therefore, we assayed if clonal *Wwc2* KD (siRNA) altered TE versus ICM fate in 32-cell stage equivalent blastocysts, by creating stably marked *Wwc2* KD and control cell clones (expressing recombinant GAP43-GFP as an injection membrane marker; microinjecting one blastomere at the 2-cell stage) and assaying expression of the Hippo-sensitive TE marker CDX2 (Strumpf et al., 2005) (Fig. 6). Control siRNA embryos developed appropriately to form blastocysts containing an average of 59% outer (CDX2 positive) and 41% inner (CDX2 negative) cell populations, with statistically equal contributions from each clone (either control siRNA microinjected or non-microinjected derived). As expected, *Wwc2* siRNA microinjected embryos had significantly fewer cells and the microinjected clone exhibited abnormal nuclear morphologies and a robustly significant impaired ICM contribution; on average 1.6±0.2 cells versus 6.8±0.3 in the non-microinjected clone (or 7.3±0.4 or 7.4±0.3 in the equivalent control siRNA embryo clones, respectively). Clonal TE contribution was also significantly impaired but to a lesser degree (overall TE numbers being compensated by increased non-microinjected clone contribution). Small numbers of outer-cell *Wwc2* KD clones failed to express CDX2 (~20%; not observed in control embryos) with the remainder exhibiting reduced immunoreactivity as compared to the non-microinjected clone (or either clone in control siRNA groups; concurrently IF stained and imaged; Fig. 6b). However, no ectopic inner-cell CDX2 expression within *Wwc2* KD clones was observed and non-microinjected sister clones appropriately expressed CDX2 in outer-cells only. Consequently, in *Wwc2* siRNA microinjected embryos the overall composition of outer versus inner-cells was skewed in favour of outer TE (largely CDX2 positive – 72%) over ICM (exclusively CDX2 negative - 28%). These results suggest that outer-cell Hippo-pathway suppression (specifying TE) and inner-cell activation (preventing TE differentiation and promoting pluripotency) are predominately intact within *Wwc2* KD clones, although outer-cell maintenance of TE differentiation is not optimal. Indeed, a lack of overt outer-cell apical polarity defects (assaying PARD6B) or ectopic nuclear exclusion of YAP (a correlate of activated Hippo-signalling (Nishioka et al., 2009)) within *Wwc2* KD outer blastomeres (Fig. S7) support this conclusion. Although interestingly, YAP nuclear exclusion in inner-*Wwc2* KD cells was consistently less robust than in control siRNA microinjected inner-cells; indicating potential impairment of inner-cell Hippo-signalling activation (Fig. S7 – although note, observed nuclear YAP was far from equivalent to outer residing blastomeres).

**Figure 6:**
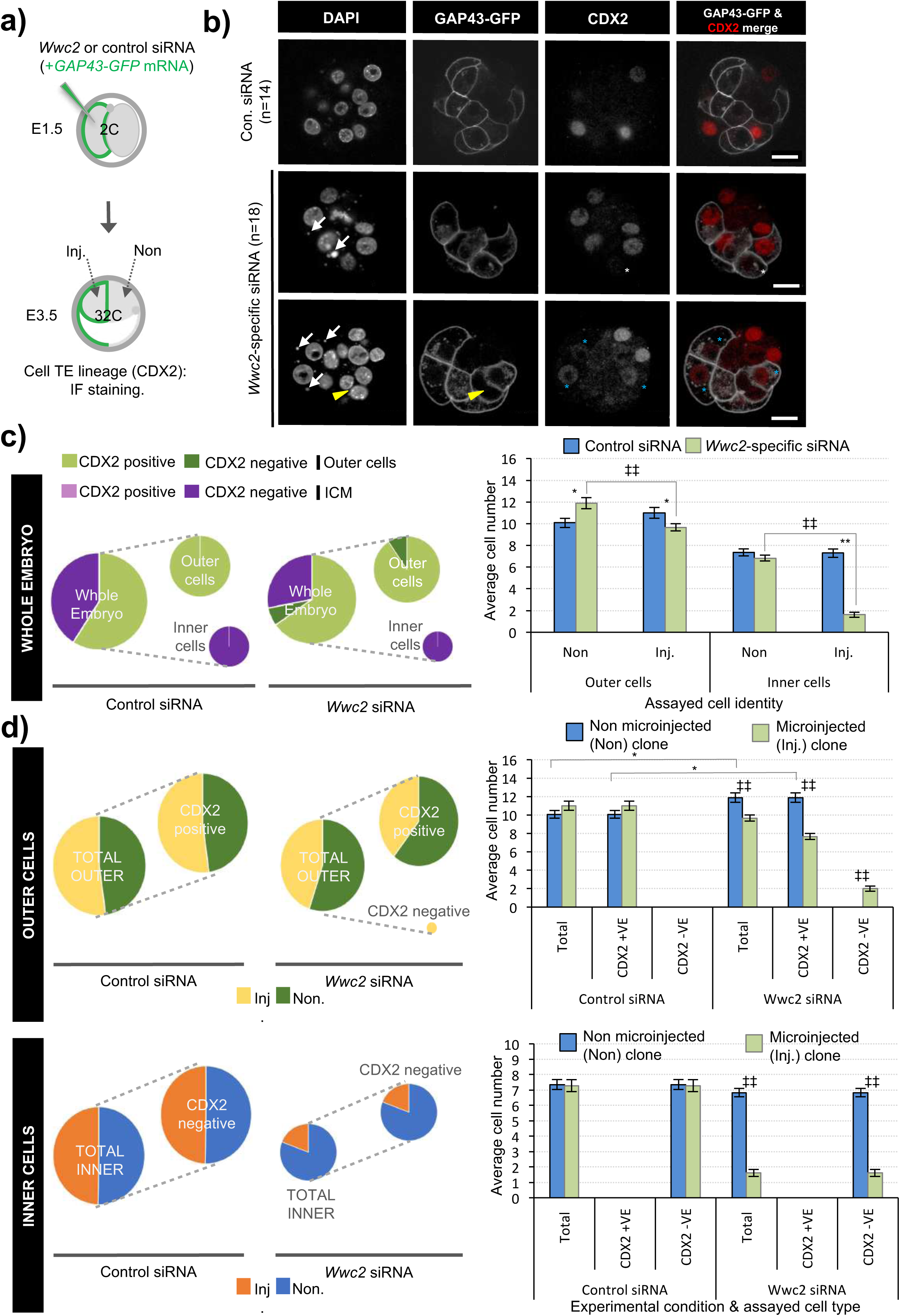
Outer trophectoderm (TE) versus ICM formation in clonal *Wwc2* gene KD at early blastocyst stage. **a)** Experimental strategy for *Wwc2* expression KD in marked clones (representing 50% of embryo) by co-microinjection (in one blastomere of 2-cell stage embryos) of *Wwc2* siRNA and GAP43-GFP mRNA (stable plasma membrane fluorescent injection clone marker – Inj.) and IF based assay of clonal contribution (Inj vs. Non) to CDX2 (TE marker) positive or negative cells, plus outer and inner-cell embryo populations at 32-cell stage (versus control siRNA microinjections). **b)** Exemplar confocal z-section micrographs of IF stained blastocysts (individual greyscale channels plus merged image; CDX2 is red) from control and *Wwc2* siRNA microinjected groups. Note, white arrows and yellow arrow head denote DAPI stained micronuclei and a bi-nucleated cell, respectively, uniquely found in *Wwc2* siRNA microinjected clones. Blue and white asterisks denote respective low level (compared to non-injected clone) or undetectable outer-cell CDX2 protein expression in *Wwc2* KD clones. The number of embryos in each group is provided; scale bar equals 20μm. **c)** Left - pie charts detailing average distribution of CDX2 protein expression (positive or negative) between control and *Wwc2* siRNA groups, in both outer or ICM cells (note, pie chart areas for each spatial population are scaled to that summarising the whole embryo; similarly, pie chart area describing the whole embryo in the *Wwc2* siRNA condition is scaled to that in the control siRNA group). Right - average clonal contribution (Inj vs. Non) of outer and ICM cells in control (blue bars) and *Wwc2* siRNA (green bars) microinjected embryos, irrespective of CDX2 expression. **d)** As in c) only describing clonal contribution of CDX2 expressing (or not) cells to outer and ICM populations, as a fraction of the total sub-population size (left - pie charts) or in raw average cell number (right – bar charts) In panel c) and d) charts, errors represent s.e.m. and 2-tailed student t-test derived statistical significance between control and *Wwc2* KD groups (asterisks), or clones (Inj. vs Non) within a group (double crosses) are highlighted with statistical confidence intervals of p<0.05 and p<0.005, as denoted by one or two significance markers, respectively. Supplementary tables ST20 summarise statistical analysis and individual embryo data.

We extended the IF staining analyses to late blastocysts (E4.5), assaying TE and EPI (CDX2 & NANOG) or PrE and EPI markers (GATA4 & NANOG or GATA4 & SOX2) as pairwise combinations (Figs. 7 & S8). *Wwc2* KD clone specific cell number (plus total blastocyst cell number) deficits, nuclear morphology and cell division phenotypes were again observed. However, in contrast to early blastocysts, the ratio of total outer (TE) to inner (ICM) cells was not significantly different between *Wwc2* and control siRNA microinjection groups (assayed across all lineage combinations; Fig. 7d-f; note, data shown as pie charts to emphasise similar TE:ICM ratios in embryos with significantly different numbers of cells – see Fig. S8 for bar charts of absolute numbers); indicative of regulative development during blastocyst maturation. However, the incidence of fragmented nuclei, consistent with apoptosis, was significantly higher within the *Wwc2* siRNA microinjected clones of the ICM, versus equivalent control siRNA clones (Fig. 7b). Consistently, the overall and significantly reduced contributions of *Wwc2* siRNA clones were more pronounced for ICM versus outer-cell populations (Fig. S8). Focussing, on CDX2 and NANOG stained groups, a population of CDX2 negative outer-cells and generally reduced CDX2 expression within the *Wwc2* KD outer-residing clone was again observed (Fig. 7c). Interestingly, the significantly reduced *Wwc2* KD clone derived ICM cell numbers did not segregate, as per marked control siRNA clones, between NANOG positive (indicative of EPI) and NANOG negative (potentially PrE) populations. Rather, they were biased towards NANOG negative cells, representative of potential PrE (Fig. 7d; again note data represented as pie charts to emphasise differences in the clonal contribution to EPI and PrE ICM lineages in embryos with significantly different numbers of cells – see Fig. S8 for bar charts of absolute numbers in each assayed ICM lineage). Considering all assayed embryos together, the overall reductions in potential PrE (NANOG negative) population size (associated with clonal *Wwc2* KD), were substantially smaller than that observed for the EPI (Fig. S8); further suggesting inner *Wwc2* KD clones are impaired in their sustained contribution to the EPI but are more able to differentiate into PrE. Data from directly assaying the two ICM lineages together (GATA4 & NANOG or SOX2; Figs. 7e & f, plus Fig. S8) also demonstrated characteristically low numbers of inner *Wwc2* KD clones, similarly segregated in favour of the PrE over EPI (although compensatory increases in EPI contribution from non-microinjected clones were noted - Fig. S8). Numerous examples of SOX2 positive fragmented inner-cell nuclei within the *Wwc2* KD clone were also noted (Fig.7c); indicating the observed increased incidence of apoptosis (Fig. 7b) was centred on specified EPI that was unable to be maintained (note, reliable quantification of SOX2 positive apoptotic nuclei was not possible). Interestingly, we noted a small population of marked ICM *Wwc2* KD clones expressing neither PrE or EPI markers (4.6% of all ICM cells and 18.7% of the clone - not present in controls), potentially indicative of further ICM cell-fate defects.

**Figure 7:**
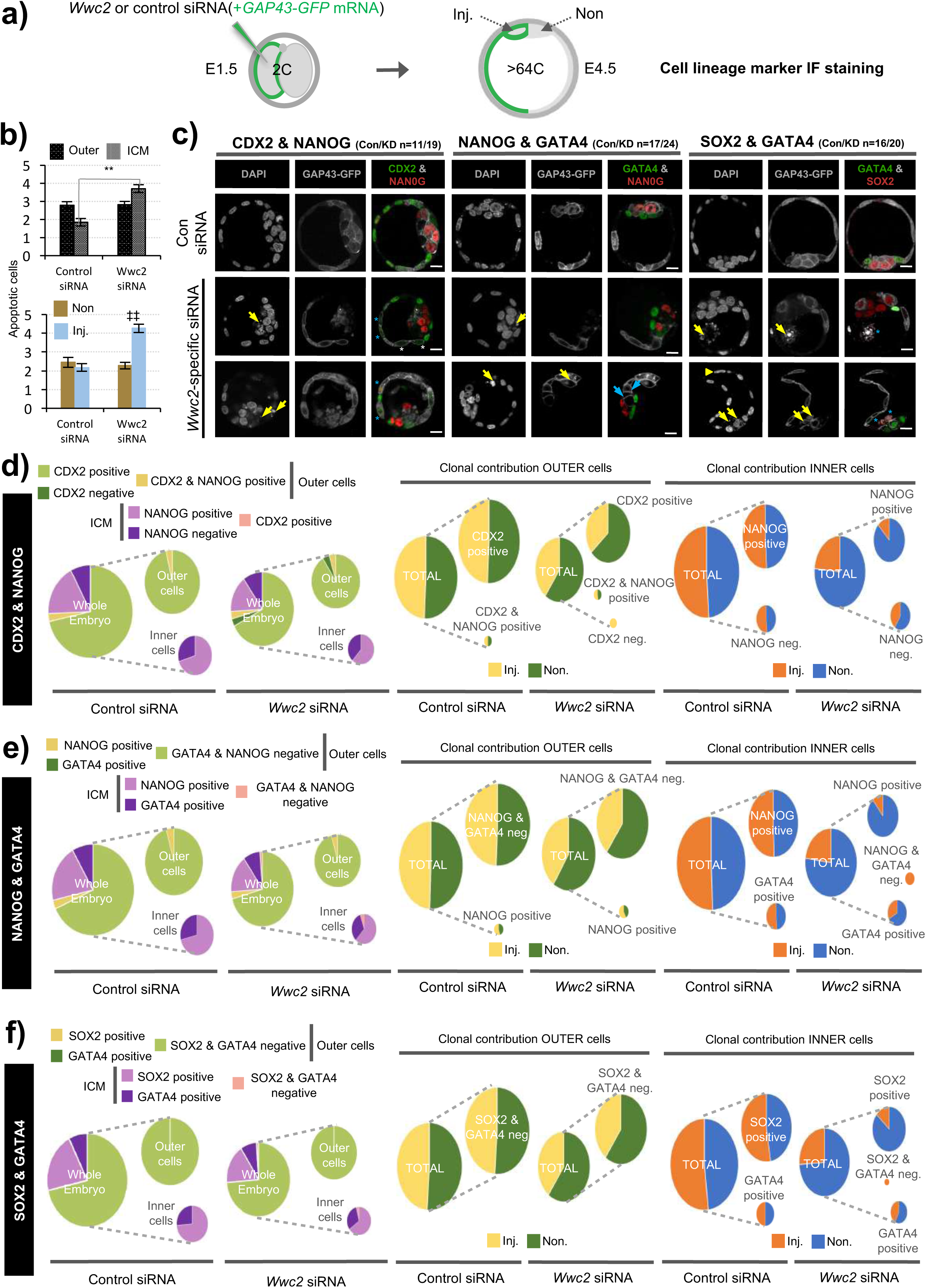
Overall cell lineage formation after clonal *Wwc2* gene KD at the late blastocyst stage. **a)** Experimental strategy for *Wwc2* expression KD in marked clones (representing 50% of the embryo) by co-microinjection (in one blastomere of 2-cell stage embryos) of *Wwc2* siRNA and GAP43-GFP mRNA (stable plasma membrane fluorescent injection clone marker lineage marker – Inj.) and an IF based assay of clonal contribution (Inj vs. Non) to blastocyst protein lineage marker expressing cells (TE; CDX2, PrE; GATA4, EPI; NANOG or SOX2; assayed in combination see below), plus outer and inner-cell embryo populations, at the late blastocyst stage (versus control siRNA microinjections). **b)** Average number of apoptotic cells in control and *Wwc2* siRNA microinjected embryo groups (averaged across all IF regimes – see below) in outer and inner cell populations and clones (Inj. vs. Non); errors represent s.e.m. and 2-tailed student t-test derived statistical significance between control and *Wwc2* KD groups (asterisks) or clones (Inj. vs Non) within a group (double crosses) highlighted with statistical confidence intervals of p<0.05 and p<0.005 (denoted by one or two significance markers, respectively). **c)** Exemplar confocal z-section micrographs of IF stained blastocysts (individual greyscale channels plus merged images, in which assayed combinations of cell lineage markers are respectively coloured green and red; CDX2 & NANOG, GATA4 & NANOG and GATA4 and SOX2) from control and *Wwc2* siRNA microinjected groups. Within the *Wwc2* siRNA microinjected clone, yellow arrows denote apoptotic cell remnants, blue arrows interphase ICM cells that neither express GATA4 or NANOG, the yellow arrow head a bi-nucleated cell whereas blue or white asterisks highlight outer-cells with basal or undetectable CDX2 expression, respectively. The number of assayed embryos in each control (Con) and *Wwc2* siRNA (KD) microinjection group is provided; scale bar equals 20μm. **d)** Pie charts detailing, (left of panel), average distribution of detectable CDX2 and NANOG protein expression (positive or negative) between control and *Wwc2* siRNA groups, in both outer or ICM cells (note, pie chart areas for each spatial population are scaled to that summarising the whole embryo; similarly the pie chart area describing the whole embryo in the *Wwc2* siRNA condition is scaled to that in the control siRNA group). Central and right panels, contain pie charts describing clonal contribution of CDX2 and NANOG expressing (or not) cells to outer and ICM populations, as a fraction of the total sub-population size. **e)** As in d) only assaying NANOG (EPI) and GATA4 (PrE) expression. **f)** As in d) only assaying SOX2 (EPI) and GATA4 (PrE) expression. Note, supplementary figure S8 details the same data in raw average cell number format (*i.e.* as presented in Fig. 6d - bar charts on right) and supplementary tables ST21-ST23 summarise statistical analysis and individual embryo data.

In summary, these data confirm *Wwc2* KD clone specific and autonomous reductions in overall embryo cell number that more robustly affect formation of ICM versus TE (compensated by regulation within the non-injected clone to preserve the overall TE:ICM ratio at the late blastocyst stage; albeit with fewer overall cells). Moreover, ICM lineage contribution of *Wwc2* KD clones is also biased against the pluripotent EPI and favours PrE differentiation, potentially via a mechanism involving selective apoptosis of specified EPI. Thus, uncovering a blastocyst cell-fate role of *Wwc2* that is additional to that related to cell division.

### Embryo *Wwc2* siRNA KD phenotypes are reversible by recombinant HA-WWC2 protein expression

To confirm the uncovered embryo related roles of *Wwc2*, we attempted phenotypic rescue experiments using the same recombinant siRNA-resistant *HA-Wwc2* mRNA construct described above (oocyte IVM – Fig. 5). We therefore repeated the marked clonal microinjection experiments but included a third condition in which *Wwc2* siRNA was co-microinjected with *HA-Wwc2* mRNA. Total and clonal cell contributions to inner and outer-cell populations, plus the incidence of abnormal nuclear morphology, were calculated in early and late blastocysts (Fig. 8). Embryos microinjected with *Wwc2* siRNA alone exhibited the typical clone autonomous phenotypes. However, under *Wwc2* siRNA conditions supplemented with co-microinjected *HA-Wwc2* mRNA (expression of which was confirmed, see below and Fig. 8d) the average cell number, clonal and spatial composition of constituent early and late blastocysts was statistically indistinguishable from control siRNA microinjection groups (except for a small reduction in ICM contribution of the non-microinjected clone at the late blastocyst stage - from 9.5±0.4 to 8.2±0.3). Additionally, a near complete rescue of abnormal morphologies was observed (only one exception across all blastomeres assayed – shown in Fig. 8c & d). These data demonstrate the specificity of the employed siRNA and confirm *Wwc2* as a novel regulatory gene of embryo cell division and subsequent cell-fate.

**Figure 8:**
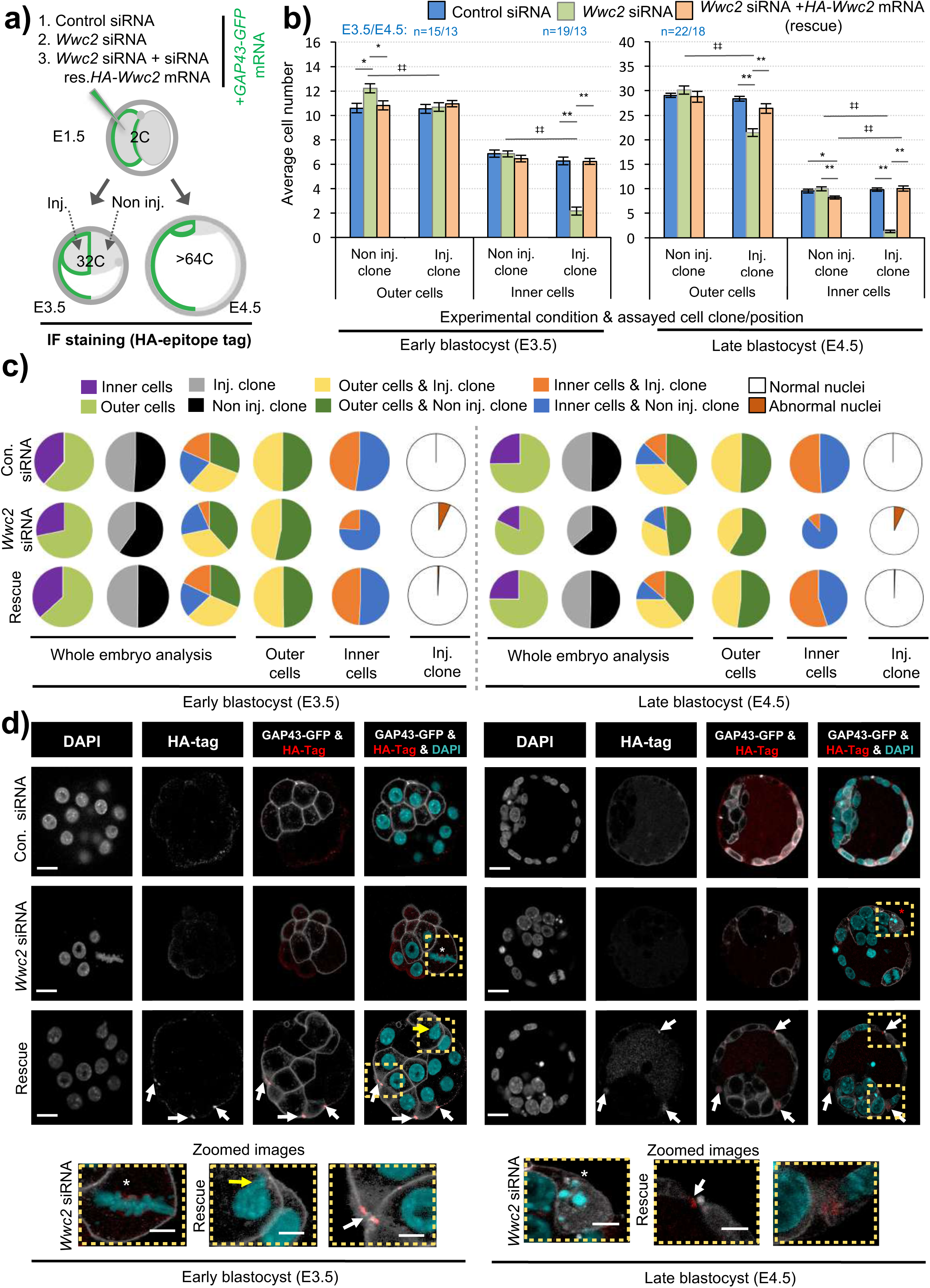
Expression of siRNA resistant *HA-Wwc2* mRNA rescues *Wwc2* siRNA cell number/division phenotypes. **a)** Experimental strategy for potential phenotypic rescue of clonal *Wwc2* KD by comparing co-microinjection (in one blastomere of 2-cell stage embryos) of GAP43-GFP mRNA (stable plasma membrane fluorescent injection marker – Inj.; to distinguish from the non-microinjected clone – Non inj.) with control siRNA, *Wwc2* siRNA or *Wwc2* siRNA + recombinant *HA-Wwc2* mRNA (containing N-terminal HA-epitope tag and point mutants conferring siRNA resistance – see supplementary figure S5) and assaying total, outer and inner-cell number at the 32-cell and late blastocyst (>64-cell) stages. **b)** Average clonal contribution (Inj vs. Non) of outer and ICM cells in control siRNA (blue bars), *Wwc2* siRNA (green bars) and *Wwc2* siRNA + *HA-Wwc2* mRNA (rescue – orange bars) microinjected embryos; errors represent s.e.m. and 2-tailed student t-test derived statistical significance between the experimental groups (asterisks), or clones (Inj. vs Non inj.) within a group (double crosses) highlighted with statistical confidence intervals of p<0.05 and p<0.005 (denoted by one or two significance markers, respectively). The number of embryos in each experimental group is provided. **c)** Pie charts detailing overall and clonal distribution of constituent embryo cells between outer and inner-cell populations, plus the incidence of abnormal nuclear morphology (as defined in Fig. 3), in each of the three indicated experimental groups, at the 32-cell (left) and late blastocyst (right) stages. Note, areas of all individual pie chart categories are normalised against those describing the appropriate criteria in the control siRNA condition. **d)** Exemplar confocal z-section micrographs of 32-cell and late blastocyst stage embryos, derived from the indicated microinjection groups, IF stained to detect HA-epitope tag expression (individual greyscale channels plus merged images - anti-HA channel is red and the DAPI cyan). Within microinjected clones of *Wwc2* siRNA treated groups, white or red asterisks respectively denote characteristic abnormal mitotic or apoptotic cell remnants, whilst in the microinjected clone of *Wwc2* siRNA + *HA-Wwc2* rescue group, white arrows indicate anti-HA epitope tag signal associated with mid-bodies, particular characteristic, in 32-cell stage embryos (see also indicated zoomed images); scale bar equals 20μm. Supplementary tables ST24 & ST25 summarise statistical analysis and individual embryo data used to generate the figure.

Incorporation of a HA-epitope tag within the confirmed *HA-Wwc2* phenotypic rescue mRNA enabled IF mediated verification of its translation and provided information on protein sub-cellular localisation within the microinjected clone. We consistently detected specific anti-HA immuno-reactivity (not observed either control or *Wwc2* siRNA microinjected embryos) at structures resembling mitotic spindle mid-bodies, particularly evident within microinjected clones at the 32-cell stage (Fig. 8d & Fig. S9a). A recent study identified a critical role for uncharacteristically persistent interphase mid-bodies (referred to as ‘interphase microtubule bridges’) as important MTOCs within cleavage stage blastomeres (Zenker et al., 2017). Therefore, we assayed their number, IF-staining for α-Tubulin and activated phospho-Aurora-B (pAURKB) as recognised midbody markers (Terada et al., 1998), in control and *Wwc2* KD embryos at the same 32-cell stage, but could not detect any statistically significant variation in their incidence (when corrected for reduced cell number caused by *Wwc2* KD - Fig. S9c-e). Hence, whilst *HA-Wwc2* derived recombinant protein associates with cell division mid-bodies, the removal of endogenous *Wwc2* mRNA does not appear to impair mid-body formation or persistence following compromised cell divisions.

## DISCUSSION

We have identified *Wwc2*, the sole murine WWC-domain containing paralog of the recognised Hippo-pathway activator *Kibra* (Baumgartner et al., 2010;Genevet et al., 2010;Yu et al., 2010;Xiao et al., 2011a;Wennmann et al., 2014), as a novel regulator of preimplantation mouse embryo mitotic cell cleavages (Figs. 1-3) and oocyte meiotic maturation (Fig. 4); the latter predicated on appropriate activation of AURKA (Figs. 5 and S6). Additionally, in the embryo we show *Wwc2* KD cell clones are compromised in EPI lineage contribution and are prone to apoptosis or disproportionate PrE contribution (Figs. 6 & 7 and S8). These data implicate at least two functional roles for *Wwc2* during early mouse development/reproduction; i) spatial and temporal regulation of germane meiotic and mitotic cell divisions (paradigms of typical and atypical acentrosomal cell division, relative to other mammalian species, respectively (Courtois et al., 2012;Bennabi et al., 2016;Severson et al., 2016;Gruss, 2018;Mogessie et al., 2018;Namgoong and Kim, 2018)), and ii) influencing blastomere pluripotency and differentiation, particularly relating to EPI and PrE formation (Figs. 6 & 7 and S8). However, the full extent of the cell-fate related role is difficult to experimentally dissect, given the described cell division defects, evident as early as the 8-cell stage.

Regarding cell-fate regulation, active Hippo-signalling is known to support initial pluripotency and resist TE differentiation in ICM founders, by blocking YAP nuclear localisation (Sasaki, 2017). However, recently it has been shown nuclear that YAP translocation is later required for blastocyst EPI progenitors to compete with each other to achieve optimal naïve pluripotency (Hashimoto and Sasaki, 2019). Hence, depending on developmental context, active Hippo-signalling appears to both initially promote and latterly suppress relative pluripotency status. Our data, although compounded by the additional cell division phenotypes, suggests *Wwc2* (as a paralog of the known Hippo-activator *Kibra* (Baumgartner et al., 2010;Genevet et al., 2010;Yu et al., 2010;Xiao et al., 2011a)) may contribute to germane Hippo-signalling mediated regulation of pluripotency. For example, we find ICM-residing late blastocyst *Wwc2* KD clones, although fewer than their non-microinjected clone counterparts, statistically favour contribution to PrE over EPI (unlike in control groups); indicative of impaired pluripotent potential (Figs. 6 & 7 and S8). Whether this is caused by compromised Hippo-signalling activation directly after initial ICM founder internalisation or is related to an inability to successfully compete for naïve pluripotency in the EPI, is not clear. However, increased apoptosis within ICM residing *Wwc2* KD clones, often marked by detectable SOX2 protein expression (but never GATA4 - Fig. 7b & c) is consistent with the competitive elimination of initially pluripotent EPI progenitors unable to achieve/compete for, naïve pluripotency. Indeed, analysis of early blastocyst stage *Wwc2* KD embryos/clones, suggests the establishment of required differential Hippo-signalling between TE (*i.e.* suppression) and ICM (*i.e.* activation) was largely intact (Figs. 6 and S7; see CDX2, PARD6B, CDH1 and YAP IF staining). Although, comparatively reduced (or sometimes absent) outer-cell CDX2 expression, plus less efficient inner-cell YAP protein nuclear exclusion, in *Wwc2* KD clones (Figs. 6 & 7 and S7) implies a degree of early onset mild Hippo-signalling dysregulation; potentially becoming more manifest during blastocyst maturation. EPI specific exclusion of *Wwc2* KD clones may also be related to increased aneuploidy caused by defective cell divisions, as derived experimental clonal aneuploidy in mouse chimeras is known to be selectively eliminated from the EPI by apoptosis but tolerated within extra-embryonic blastocyst tissues (Bolton et al., 2016). The mild early cell-fate effects may also reflect functional redundancy between WWC2 and KIBRA; indeed, robust *Wwc2* KD does not affect *Kibra* mRNA levels (Fig. 1 and S1). Functional redundancy between WWC-domain proteins paralogs has been demonstrated in human cells (relating to KIBRA, WWC2 and WWC3 (Wennmann et al., 2014)) and genetic ablation of the mouse *Kibra* gene is developmentally viable (adult mice only exhibit minor learning and memory defects (Makuch et al., 2011)). However, if KIBRA and WWC2 proteins share functional redundancy during mouse preimplantation development, it is probably only in relation to potential Hippo-signalling regulation and not applicable to the described cell division functions of *Wwc2* (as sole *Kibra* KD did not affect cell number – Fig. 1).

Given *Wwc2* specific siRNA embryo microinjections were performed at the 2-cell stage it is unclear why cell autonomous division phenotypes (particularly affecting inner-cell populations) are only manifest by the 16-cell stage (Figs. 1 - 3 and S2). One possibility is that all post-microinjection cell-cycles are elongated in a manner only revealed at the whole embryo level by the 16-cell stage, implicating *Wwc2* as a regulator of the atypically long cell-cycles of preimplantation mammalian embryo blastomeres (reviewed (O’Farrell et al., 2004)). Alternatively, it may reflect functional exhaustion of maternally provided WWC2 protein, unaffected by siRNA (although, it was impossible to directly assay WWC2 protein expression, due to unavailability of antibodies). The emergence of phenotypes at the 16-cell stage also coincides with the first spatial segregation of blastomeres (outer- and inner-positions), polarity/position dependant differential Hippo-signalling establishment and TE versus ICM cell-fate formation (Sasaki, 2017). It is tempting to speculate such developmental timing is particularly relevant in this context. However, the onset of individual nuclear morphological phenotypes, indicative of defective cell division/cytokinesis (mid-body association, bi-/multi-/micro-nucleation - Fig. 3), were already evident by the 8-cell stage. Additionally, although weak, outer-cell restricted CDX2 protein expression confirmed intact differential Hippo-signalling (Fig. 6 & 7). Interestingly, whilst average cell number was impaired by *Wwc2* KD after the 8-cell stage, we noted it did not significantly increase again until the mid-blastocyst stage (remaining around 8-10 cells - Fig. 2). Significantly, the mid-blastocyst stage marks the developmental point that atypical (compared to most other mammalian species) mouse embryo acentrosomal cell divisions revert back to centrosomal control, after *de novo* centrosome synthesis (Courtois et al., 2012). We propose our data invoke a role for WWC2 in regulating mouse preimplantation stage acentrosomal blastomere cleavage, supported by the uncovered nuclear morphology phenotypes associated with *Wwc2* KD (Fig. 3) and readily detectable recombinant HA-WWC2 protein in mitotic mid-bodies up until the early blastocyst stage (not easily detectable in late blastocysts - Figs. 8 and S9). We suspect HA-WWC2 association with mid-bodies reflects a role for WWC2 during preceeding cell divisions and that the localisation represents a remant of this role rather than affecting the previously described MTOC function of such atypically persistent mouse cleavage blastomere mid-bodies (Zenker et al., 2017); indeed, when corrected for lower overall cell numbers, the total number of detectable mid-bodies is not significantly different in *Wwc2* KD embryos from controls (Fig. S9). Moreover, the impaired oocyte maturation data, by which GV oocytes (again in the absence of centrosomes (Bennabi et al., 2016;Severson et al., 2016;Gruss, 2018;Mogessie et al., 2018;Namgoong and Kim, 2018)) depleted of *Wwc2* transcripts present with multiple spindle defects associated with failed AURKA activation (Figs. 4 & 5 and S3 & S6), further substantiates this hypothesis. Interestingly, clonal *Aurkb* and *Aurkc* downregulation in preimplantation mouse embryos, respectively increases or decreases the rate of mitotic cell division (over-expression exhibiting the opposing effects). Indeed, the reported cell number deficits associated with *Aurkc* downregulation, strongly resemble those associated with *Wwc2* KD (Li et al., 2017), suggesting a potential link between WWC2 and AURKB/AURKC function in the early embryo.

Regulation of mammalian Hippo-signalling by WWC-domain proteins was first demonstrated in human HEK293T cells, whereby direct binding of KIBRA was shown to activate LATS1/2, causing YAP phosphorylation (Xiao et al., 2011a). A subsequent report detailed AURKA/B directed and mitosis specific KIBRA phosphorylation, centred on a conserved Ser539 residue, and showed the KIBRA mutant (S539A) promotes precocious M-phase exit in models of spindle assembly check-point (SAC) mediated arrest (Xiao et al., 2011b). The same group later demonstrated full activation/phosphorylation of AURKA (hence, LATS2 activation and appropriate centrosomal localisation) requires KIBRA and that *KIBRA* KD causes spindle defects, lagging chromosomes and micronuclei formation (reminiscent of our uncovered *Wwc2* KD phenotypes), plus centrosomal fragmentation in human cell line models (Zhang et al., 2012). Interestingly, we have identified conserved AURK consensus motifs in both murine KIBRA and WWC2 proteins; including those paralogous to Ser539 (Fig. S10). We therefore hypothesise the *Wwc2* KD phenotypes we observe here are mechanistically related to those described for human KIBRA, with the caveat they function on the level of MTOC regulation due to the absence of centrosomes (Courtois et al., 2012). For example, genetic ablations of either murine *Lats1* and *Lats2*, do not cause embryo cell number deficits typical of *Wwc2* KD, nor those centrosome directed mitotic phenotypes described above, rather presenting in early blastocysts as ectopic ICM cell nuclear YAP localisations (Nishioka et al., 2009). Therefore, not all insights will be directly transferable. Notwithstanding and despite the lack of early mouse embryo centrosomes, key centrosome regulators are expressed and reported to regulate embryo cleavage divisions. For example, genetic knockout of Polo-like kinase 1 (*Plk1*), known to cooperate with AURKA within centrosomes to mediated bi-polar spindle formation (Asteriti et al., 2015), causes arrest at the same 8-cell stage at which global *Wwc2* KD embryos first exhibit cell division defects (Figs. 2 and 3) (Lu et al., 2008). Moreover, an important role in blastomere division for the related *Plk4* gene, itself recognised as a key regulator of centriole formation/duplication (Bettencourt-Dias et al., 2005;Habedanck et al., 2005), has been demonstrated. Accordingly, active PLK4 protein localises to chromatin proximal and coalescing MTOCs to promote microtubule nucleation and bi-polar spindle formation (Coelho et al., 2013). It will be interesting to investigate the identified *Wwc2* KD phenotypes in the context of such pre-existing and characterised MTOC-related factors and to test their applicability within the conceptually similar paradigm of acentrosomal meiotic oocyte maturation. Indeed, upon resumption of meiosis, PLK4 was recently described to cooperate with AURKA (within MTOCs) to initiate microtubule nucleation around condensing chromosomes. Functional inhibition of PLK4 did not however block spindle assembly but increased assembly time and the individual MTOC regulatory roles of PLK4 and AURKA were shown to be only partially overlapping (Bury et al., 2017) and distinct from the co-existing chromosome derived RAN-GTP gradient driven mechanisms (as reviewed (Bennabi et al., 2016;Severson et al., 2016;Gruss, 2018;Mogessie et al., 2018;Namgoong and Kim, 2018)). Importantly, the described functional inhibition of PLK4 was associated with eventual polar body formation (Bury et al., 2017), unlike after *Wwc2* KD (even after an extended IVM incubation time – Fig. S3). This indicates that whilst meiotic spindle assembly is impaired to varying degrees by *Wwc2* KD (Fig. 4d & d’), other impediments to successful meiotic maturation must exist (potentially relating, although not necessarily limited, to spindle migration, SAC, cytokinesis *etc.*). The comparative distribution of observed meiotic maturation phenotypes caused by *Wwc2* KD (*i.e.* failed spindle formation, defective spindles or persistent/ non-dividing MI spindles – Fig. 4d & d’) suggests some functional redundancy in response to loss of WWC2. This may be potentially related to the compensatory abilities of the three expressed AURK paralogs (*Aurka*, *Aurkb* and *Aurkc*), as demonstrated in combinatorial genetic ablations in mouse oocytes (Nguyen et al., 2018).

In summary, we have identified the ill-characterised WWC-domain containing gene *Wwc2*, as an important regulator of cell division in both preimplantation mouse embryo blastomeres and oocytes; both paradigms of acentrosomal cell division. We have also uncovered evidence for *Wwc2*, as a paralog of the described Hippo-signalling activator *Kibra*, in participating in blastocyst cell-fate regulation and pluripotency. Despite the compounding nature of the two identified *Wwc2* KD phenotypes, it will be of great future interest to further our molecular understanding of these newly identified roles of the *Wwc2* gene during early mouse development.

## Supporting information

Supplementary Info (Suppl. Figs + Suppl. Methods Tables + Suppl. Tables (individual embryo data & statistics)

## ACKNOWLEDGEMENTS

Authors acknowledge Marta Gajewska (Institute of Oncology, Warsaw, Poland) and Anna Piliszek (Institute of Genetics & Animal Breeding, Polish Academy of Sciences, Jastrzębiec, Poland) for providing founder CBA/W mice used in this study, Martin Anger (Central European Institute of Technology (CEITEC)/ Veterinary Research Institute, Brno, Czech Republic) for valuable technical advice and discussions and Alena Krejčí (Faculty of Science, University of South Bohemia, České Budějovice, Czech Republic) for advice and pooling resources. The Animal Facility (Institute of Parasitology, Biology Centre of the Czech Academy of Sciences, České Budějovice, Czech Republic) is acknowledged for housing experimental mice. The research was supported by a Czech Science Foundation/ GA ČR grant (18-02891S) awarded to A.W.B. and a Grant Agency of the University of South Bohemia Ph.D. student award to G.V (GA JU: 015/2017/P).

## CONFLICT OF INTEREST

The authors declare that they have no conflict of interests.

## Notes

### Competing Interest Statement

The authors have declared no competing interest.

### Summary of Updates

Reformatted and edited text

## REFERENCES

Asteriti, I.A., De Mattia, F., and Guarguaglini, G. (2015). Cross-Talk between AURKA and Plk1 in Mitotic Entry and Spindle Assembly. Front Oncol 5, 283.

Baumgartner, R., Poernbacher, I., Buser, N., Hafen, E., and Stocker, H. (2010). The WW domain protein Kibra acts upstream of Hippo in Drosophila. Dev Cell 18, 309–316.

Bennabi, I., Terret, M.E., and Verlhac, M.H. (2016). Meiotic spindle assembly and chromosome segregation in oocytes. J Cell Biol 215, 611–619.

Bettencourt-Dias, M., Rodrigues-Martins, A., Carpenter, L., Riparbelli, M., Lehmann, L., Gatt, M.K., Carmo, N., Balloux, F., Callaini, G., and Glover, D.M. (2005). SAK/PLK4 is required for centriole duplication and flagella development. Curr Biol 15, 2199–2207.

Bolton, H., Graham, S.J.L., Van Der Aa, N., Kumar, P., Theunis, K., Fernandez Gallardo, E., Voet, T., and Zernicka-Goetz, M. (2016). Mouse model of chromosome mosaicism reveals lineage-specific depletion of aneuploid cells and normal developmental potential. Nat Commun 7, 11165.

Bury, L., Coelho, P.A., Simeone, A., Ferries, S., Eyers, C.E., Eyers, P.A., Zernicka-Goetz, M., and Glover, D.M. (2017). Plk4 and Aurora A cooperate in the initiation of acentriolar spindle assembly in mammalian oocytes. J Cell Biol 216, 3571–3590.

Chazaud, C., and Yamanaka, Y. (2016). Lineage specification in the mouse preimplantation embryo. Development 143, 1063–1074.

Coelho, P.A., Bury, L., Sharif, B., Riparbelli, M.G., Fu, J., Callaini, G., Glover, D.M., and Zernicka-Goetz, M. (2013). Spindle formation in the mouse embryo requires Plk4 in the absence of centrioles. Dev Cell 27, 586–597.

Courtois, A., Schuh, M., Ellenberg, J., and Hiiragi, T. (2012). The transition from meiotic to mitotic spindle assembly is gradual during early mammalian development. J Cell Biol 198, 357–370.

Davis, J.R., and Tapon, N. (2019). Hippo signalling during development. Development 146.

Frum, T., Murphy, T.M., and Ralston, A. (2018). HIPPO signaling resolves embryonic cell fate conflicts during establishment of pluripotency in vivo. Elife 7.

Frum, T., and Ralston, A. (2015). Cell signaling and transcription factors regulating cell fate during formation of the mouse blastocyst. Trends Genet 31, 402–410.

Frum, T., Watts, J.L., and Ralston, A. (2019). TEAD4, YAP1 and WWTR1 prevent the premature onset of pluripotency prior to the 16-cell stage. Development 146.

Genevet, A., Wehr, M.C., Brain, R., Thompson, B.J., and Tapon, N. (2010). Kibra is a regulator of the Salvador/Warts/Hippo signaling network. Dev Cell 18, 300–308.

Gruss, O.J. (2018). Animal Female Meiosis: The Challenges of Eliminating Centrosomes. Cells 7.

Habedanck, R., Stierhof, Y.D., Wilkinson, C.J., and Nigg, E.A. (2005). The Polo kinase Plk4 functions in centriole duplication. Nat Cell Biol 7, 1140–1146.

Han, Q., Kremerskothen, J., Lin, X., Zhang, X., Rong, X., Zhang, D., and Wang, E. (2018). WWC3 inhibits epithelial-mesenchymal transition of lung cancer by activating Hippo-YAP signaling. Onco Targets Ther 11, 2581–2591.

Hashimoto, M., and Sasaki, H. (2019). Epiblast Formation by TEAD-YAP-Dependent Expression of Pluripotency Factors and Competitive Elimination of Unspecified Cells. Dev Cell 50, 139–154 e135.

Hassold, T., and Hunt, P. (2001). To err (meiotically) is human: the genesis of human aneuploidy. Nat Rev Genet 2, 280–291.

Hirate, Y., Hirahara, S., Inoue, K., Kiyonari, H., Niwa, H., and Sasaki, H. (2015). Par-aPKC-dependent and -independent mechanisms cooperatively control cell polarity, Hippo signaling, and cell positioning in 16-cell stage mouse embryos. Dev Growth Differ 57, 544–556.

Horn, T., and Boutros, M. (2010). E-RNAi: a web application for the multi-species design of RNAi reagents--2010 update. Nucleic Acids Res 38, W332–339.

Johnson, M.H., and Ziomek, C.A. (1981). The foundation of two distinct cell lineages within the mouse morula. Cell 24, 71–80.

Koncicka, M., Tetkova, A., Jansova, D., Del Llano, E., Gahurova, L., Kracmarova, J., Prokesova, S., Masek, T., Pospisek, M., Bruce, A.W., Kubelka, M., and Susor, A. (2018). Increased Expression of Maturation Promoting Factor Components Speeds Up Meiosis in Oocytes from Aged Females. Int J Mol Sci 19.

Kovarikova, V., Burkus, J., Rehak, P., Brzakova, A., Solc, P., and Baran, V. (2016). Aurora kinase A is essential for correct chromosome segregation in mouse zygote. Zygote 24, 326–337.

Lemaire, P., Garrett, N., and Gurdon, J.B. (1995). Expression cloning of Siamois, a Xenopus homeobox gene expressed in dorsal-vegetal cells of blastulae and able to induce a complete secondary axis. Cell 81, 85–94.

Li, W., Wang, P., Zhang, B., Zhang, J., Ming, J., Xie, W., and Na, J. (2017). Differential regulation of H3S10 phosphorylation, mitosis progression and cell fate by Aurora Kinase B and C in mouse preimplantation embryos. Protein Cell 8, 662–674.

Livak, K.J., and Schmittgen, T.D. (2001). Analysis of relative gene expression data using real-time quantitative PCR and the 2(-Delta Delta C(T)) Method. Methods 25, 402–408.

Lu, L.Y., Wood, J.L., Minter-Dykhouse, K., Ye, L., Saunders, T.L., Yu, X., and Chen, J. (2008). Polo-like kinase 1 is essential for early embryonic development and tumor suppression. Mol Cell Biol 28, 6870–6876.

Makuch, L., Volk, L., Anggono, V., Johnson, R.C., Yu, Y., Duning, K., Kremerskothen, J., Xia, J., Takamiya, K., and Huganir, R.L. (2011). Regulation of AMPA receptor function by the human memory-associated gene KIBRA. Neuron 71, 1022–1029.

Mihajlovic, A.I., and Bruce, A.W. (2016). Rho-associated protein kinase regulates subcellular localisation of Angiomotin and Hippo-signalling during preimplantation mouse embryo development. Reprod Biomed Online 33, 381–390.

Mihajlovic, A.I., and Bruce, A.W. (2017). The first cell-fate decision of mouse preimplantation embryo development: integrating cell position and polarity. Open Biol 7.

Mihajlovic, A.I., and Fitzharris, G. (2018). Segregating Chromosomes in the Mammalian Oocyte. Curr Biol 28, R895–R907.

Mihajlovic, A.I., Thamodaran, V., and Bruce, A.W. (2015). The first two cell-fate decisions of preimplantation mouse embryo development are not functionally independent. Sci Rep 5, 15034.

Mogessie, B., Scheffler, K., and Schuh, M. (2018). Assembly and Positioning of the Oocyte Meiotic Spindle. Annu Rev Cell Dev Biol 34, 381–403.

Morris, S.A., Teo, R.T., Li, H., Robson, P., Glover, D.M., and Zernicka-Goetz, M. (2010). Origin and formation of the first two distinct cell types of the inner cell mass in the mouse embryo. Proc Natl Acad Sci U S A 107, 6364–6369.

Nagaoka, S.I., Hassold, T.J., and Hunt, P.A. (2012). Human aneuploidy: mechanisms and new insights into an age-old problem. Nat Rev Genet 13, 493–504.

Namgoong, S., and Kim, N.H. (2018). Meiotic spindle formation in mammalian oocytes: implications for human infertility. Biol Reprod 98, 153–161.

Nguyen, A.L., Drutovic, D., Vazquez, B.N., El Yakoubi, W., Gentilello, A.S., Malumbres, M., Solc, P., and Schindler, K. (2018). Genetic Interactions between the Aurora Kinases Reveal New Requirements for AURKB and AURKC during Oocyte Meiosis. Curr Biol 28, 3458–3468 e3455.

Nguyen, A.L., and Schindler, K. (2017). Specialize and Divide (Twice): Functions of Three Aurora Kinase Homologs in Mammalian Oocyte Meiotic Maturation. Trends Genet 33, 349–363.

Nishioka, N., Inoue, K., Adachi, K., Kiyonari, H., Ota, M., Ralston, A., Yabuta, N., Hirahara, S., Stephenson, R.O., Ogonuki, N., Makita, R., Kurihara, H., Morin-Kensicki, E.M.,, Nojima, H., Rossant, J., Nakao, K., Niwa, H., and Sasaki, H. (2009). The Hippo signaling pathway components Lats and Yap pattern Tead4 activity to distinguish mouse trophectoderm from inner cell mass. Dev Cell 16, 398–410.

O’farrell, P.H., Stumpff, J., and Su, T.T. (2004). Embryonic cleavage cycles: how is a mouse like a fly? Curr Biol 14, R35–45.

Rossant, J. (2016). Making the Mouse Blastocyst: Past, Present, and Future. Curr Top Dev Biol 117, 275–288.

Rossant, J. (2018). Genetic Control of Early Cell Lineages in the Mammalian Embryo. Annu Rev Genet 52, 185–201.

Salles, F.J., Darrow, A.L., O’connell, M.L., and Strickland, S. (1992). Isolation of novel murine maternal mRNAs regulated by cytoplasmic polyadenylation. Genes Dev 6, 1202–1212.

Sanders, J.R., and Jones, K.T. (2018). Regulation of the meiotic divisions of mammalian oocytes and eggs. Biochem Soc Trans 46, 797–806.

Sasaki, H. (2017). Roles and regulations of Hippo signaling during preimplantation mouse development. Dev Growth Differ 59, 12–20.

Saskova, A., Solc, P., Baran, V., Kubelka, M., Schultz, R.M., and Motlik, J. (2008). Aurora kinase A controls meiosis I progression in mouse oocytes. Cell Cycle 7, 2368–2376.

Sathananthan, A.H., Kola, I., Osborne, J., Trounson, A., Ng, S.C., Bongso, A., and Ratnam, S.S. (1991). Centrioles in the beginning of human development. Proc Natl Acad Sci U S A 88, 4806–4810.

Schatten, H., and Sun, Q.Y. (2009). The role of centrosomes in mammalian fertilization and its significance for ICSI. Mol Hum Reprod 15, 531–538.

Severson, A.F., Von Dassow, G., and Bowerman, B. (2016). Oocyte Meiotic Spindle Assembly and Function. Curr Top Dev Biol 116, 65–98.

Solc, P., Baran, V., Mayer, A., Bohmova, T., Panenkova-Havlova, G., Saskova, A., Schultz, R.M., and Motlik, J. (2012). Aurora kinase A drives MTOC biogenesis but does not trigger resumption of meiosis in mouse oocytes matured in vivo. Biol Reprod 87, 85.

Strumpf, D., Mao, C.A., Yamanaka, Y., Ralston, A., Chawengsaksophak, K., Beck, F., and Rossant, J. (2005). Cdx2 is required for correct cell fate specification and differentiation of trophectoderm in the mouse blastocyst. Development 132, 2093–2102.

Swain, J.E., Ding, J., Wu, J., and Smith, G.D. (2008). Regulation of spindle and chromatin dynamics during early and late stages of oocyte maturation by aurora kinases. Mol Hum Reprod 14, 291–299.

Terada, Y., Tatsuka, M., Suzuki, F., Yasuda, Y., Fujita, S., and Otsu, M. (1998). AIM-1: a mammalian midbody-associated protein required for cytokinesis. EMBO J 17, 667–676.

Wang, Q.T., Piotrowska, K., Ciemerych, M.A., Milenkovic, L., Scott, M.P. Davis, R.W., and Zernicka-Goetz, M. (2004). A genome-wide study of gene activity reveals developmental signaling pathways in the preimplantation mouse embryo. Dev Cell 6, 133–144.

Wennmann, D.O., Schmitz, J., Wehr, M.C., Krahn, M.P., Koschmal, N., Gromnitza, S., Schulze, U., Weide, T., Chekuri, A., Skryabin, B.V., Gerke, V., Pavenstadt, H., Duning, K., and Kremerskothen, J. (2014). Evolutionary and molecular facts link the WWC protein family to Hippo signaling. Mol Biol Evol 31, 1710–1723.

White, M.D., Zenker, J., Bissiere, S., and Plachta, N. (2018). Instructions for Assembling the Early Mammalian Embryo. Dev Cell 45, 667–679.

Wicklow, E., Blij, S., Frum, T., Hirate, Y., Lang, R.A., Sasaki, H., and Ralston, A. (2014). HIPPO pathway members restrict SOX2 to the inner cell mass where it promotes ICM fates in the mouse blastocyst. PLoS Genet 10, e1004618.

Xiao, L., Chen, Y., Ji, M., and Dong, J. (2011a). KIBRA regulates Hippo signaling activity via interactions with large tumor suppressor kinases. J Biol Chem 286, 7788–7796.

Xiao, L., Chen, Y., Ji, M., Volle, D.J., Lewis, R.E., Tsai, M.Y., and Dong, J. (2011b). KIBRA protein phosphorylation is regulated by mitotic kinase aurora and protein phosphatase 1. J Biol Chem 286, 36304–36315.

Yu, J., Zheng, Y., Dong, J., Klusza, S., Deng, W.M., and Pan, D. (2010). Kibra functions as a tumor suppressor protein that regulates Hippo signaling in conjunction with Merlin and Expanded. Dev Cell 18, 288–299.

Zenker, J., White, M.D., Templin, R.M., Parton, R.G., Thorn-Seshold, O., Bissiere, S., and Plachta, N. (2017). A microtubule-organizing center directing intracellular transport in the early mouse embryo. Science 357, 925–928.

Zernicka-Goetz, M., Pines, J., Ryan, K., Siemering, K.R., Haseloff, J., Evans, M.J., and Gurdon, J.B. (1996). An indelible lineage marker for Xenopus using a mutated green fluorescent protein. Development 122, 3719–3724.

Zhang, L., Iyer, J., Chowdhury, A., Ji, M., Xiao, L., Yang, S., Chen, Y., Tsai, M.Y., and Dong, J. (2012). KIBRA regulates aurora kinase activity and is required for precise chromosome alignment during mitosis. J Biol Chem 287, 34069–34077.

Zhang, Y., Yan, S., Chen, J., Gan, C., Chen, D., Li, Y., Wen, J., Kremerskothen, J., Chen, S., Zhang, J., and Cao, Y. (2017). WWC2 is an independent prognostic factor and prevents invasion via Hippo signalling in hepatocellular carcinoma. J Cell Mol Med 21, 3718–3729.

