## Supplementary Info (Suppl. Figs + Suppl. Methods Tables + Suppl. Tables (individual embryo data & statistics) for "*Wwc2* is a novel cell division regulator during preimplantation mouse embryo lineage formation and oogenesis"

Virnicchi *et al.,* 2020

INC.

A) SUPPLMENTARY FIGURES & LEGENDS: **FIG. S1 – FIG. S10**

B) SUPPLMENTARY METHODS TABLES: **SM1 – SM6**

**C)** SUPPLMENTARY TABLES (individual embryo cell counts & oocyte phenotypes + statistics): **ST1 – ST26**

A) SUPPLMENTARY FIGURES & LEGENDS: FIG. S1 – FIG. S10

**_Wwc2_ is a novel cell division regulator during preimplantation mouse embryo lineage formation and oogenesis.**

Virnicchi *et al.,* 2020

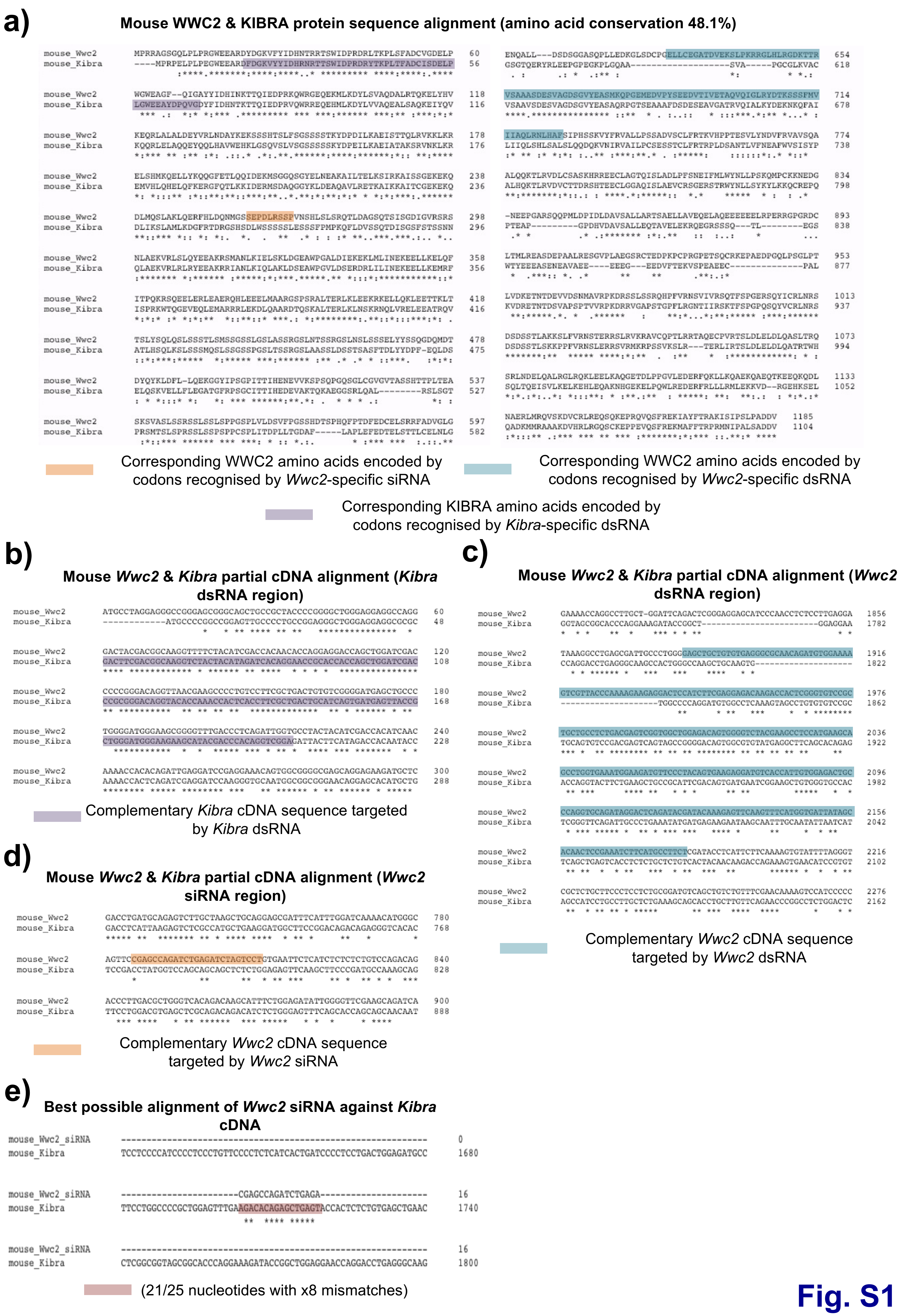
a)
Mouse WWC2 & KIBRA protein sequence alignment (amino acid conservation 48.1%)
Corresponding WWC2 amino acids encoded by codons recognised by Wwc2-specific siRNA
Corresponding WWC2 amino acids encoded by codons recognised by Wwc2-specific dsRNA
Corresponding KIBRA amino acids encoded by codons recognised by Kibra-specific dsRNA
b)
Mouse Wwc2 & Kibra partial cDNA alignment (Kibra dsRNA region)
Complementary Kibra cDNA sequence targeted by Kibra dsRNA
c)
Mouse Wwc2 & Kibra partial cDNA alignment (Wwc2 dsRNA region)
Complementary Wwc2 cDNA sequence targeted by Wwc2 dsRNA
d)
Mouse Wwc2 & Kibra partial cDNA alignment (Wwc2 siRNA region)
Complementary Wwc2 cDNA sequence targeted by Wwc2 siRNA
e)
Best possible alignment of Wwc2 siRNA against Kibra cDNA
(21/25 nucleotides with x8 mismatches)

**Fig. S1**

**Supplementary figure S1: Annotated amino acid and cDNA sequence alignments for mouse WWC2/*Wwc2* & KIBRA/*Kibra* detailing regions targeted by dsRNA and siRNA constructs. a)** Full-length amino acid sequence of mouse WWC2 and KIBRA proteins; purple denotes the region encoded by codons targeted by the *Kibra*-specific dsRNA, orange that targeted by the *Wwc2*-specific siRNA, turquoise targeted by the *Wwc2*-specifc dsRNA. **b)** Partial cDNA sequence alignment of the *Kibra* gene region coding sequence targeted by the *Kibra*-specific dsRNA (highlighted in purple), against the equivalent/aligned region in *Wwc2* cDNA. **c)** Partial cDNA sequence alignment of the region of the *Wwc2* gene coding sequence targeted by the *Wwc2*-specific dsRNA (highlighted in turquoise), against the equivalent/aligned region in *Kibra* cDNA. **d)** Partial cDNA sequence alignment of the *Wwc2* gene region sequence targeted by the *Wwc2*-specific siRNA (highlighted in orange), against the equivalent/aligned region in K*ibra* cDNA. **e)** The best result of aligning the used *Wwc2*-specific siRNA sequence against the mouse *Kibra* cDNA sequence (highlighted in red); note the best possible match of the *Wwc2-specific siRNA* sequence to the *Kibra* cDNA sequence is a 21 nucleotide stretch that contains eight evenly spread base-pair mis-matches, thus, minimising the chance of off-target recognition of *Kibra* transcripts by the *Wwc2*-designed siRNA. All alignments were conducted using default setting of the online Clustal-Ω multiple sequence alignment tool; asterisks (*) denote perfectly aligned and identical amino acids/DNA base-pairs, whereas colons (:) detail aligned conservative amino acid substitutions.

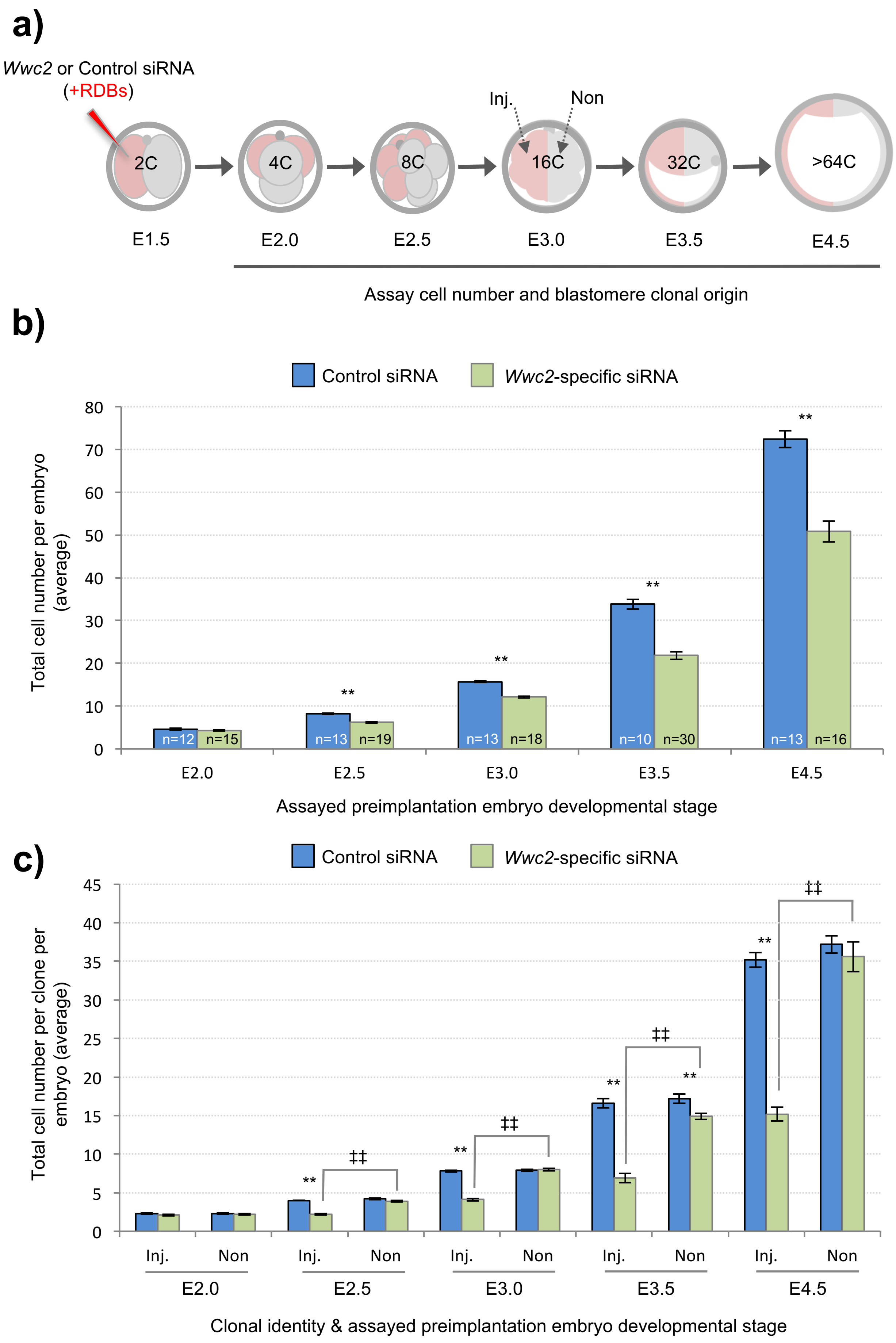
a)
Wwc2 or Control siRNA
(+RDBs)
Inj.
Non
2C
4C
8C
16C
32C
>64C
E1.5
E2.0
E2.5
E3.0
E3.5
E4.5
Assay cell number and blastomere clonal origin
b)
Control siRNA
Wwc2-specific siRNA
Total cell number per embryo (average)
80
70
60
50
40
30
20
10
0
**
n=12
n=15
n=13
n=19
n=13
n=18
n=10
n=30
n=13
n=16
E2.0
E2.5
E3.0
E3.5
E4.5
Assayed preimplantation embryo developmental stage
c)
Control siRNA
Wwc2-specific siRNA
Total cell number per clone per embryo (average)
45
40
35
30
25
20
15
10
5
0
**
‡‡
Inj.
Non
E2.0
E2.5
E3.0
E3.5
E4.5
Clonal identity & assayed preimplantation embryo developmental stage

**Fig. S2**

**Supplementary figure S2: Preimplantation embryo *Wwc2* KD associated cell division phenotypes are cell autonomous. a)** Experimental strategy for *Wwc2* KD in a marked clone (representing 50% of the embryo) by co-microinjection (in one blastomere of 2-cell stage embryos) of *Wwc2* siRNA and RDBs (fluorescent lineage marker – Inj.) and an assay of clonal contribution (Inj vs. Non) at stated developmental stages (compared to similar control siRNA microinjections). **b)** Average total number of cells per embryo (irrespective of clonal origin), at each developmental stage, from control siRNA (blue bars) and *Wwc2*-specifc siRNA microinjected (green bars) embryos. The number of embryos in each experimental group is shown. **c)** As in b) but detailing the clonal contribution of cells (Inj vs. Non) to total embryo cell number in control siRNA (blue bars) or *Wwc2* siRNA (green bars) microinjected embryos. In panels b) & c) chart error bars represent s.e.m. and statistically significant differences (2-tailed students t-test) between the experimental microinjection groups (asterisks) or clones within a group (double crosses) are highlighted with statistical confidence intervals of $p<0.05$ and $p<0.005$ (denoted by one or two significance markers, respectively). Supplementary tables ST9-ST13 summarise statistical analysis and individual embryo data.

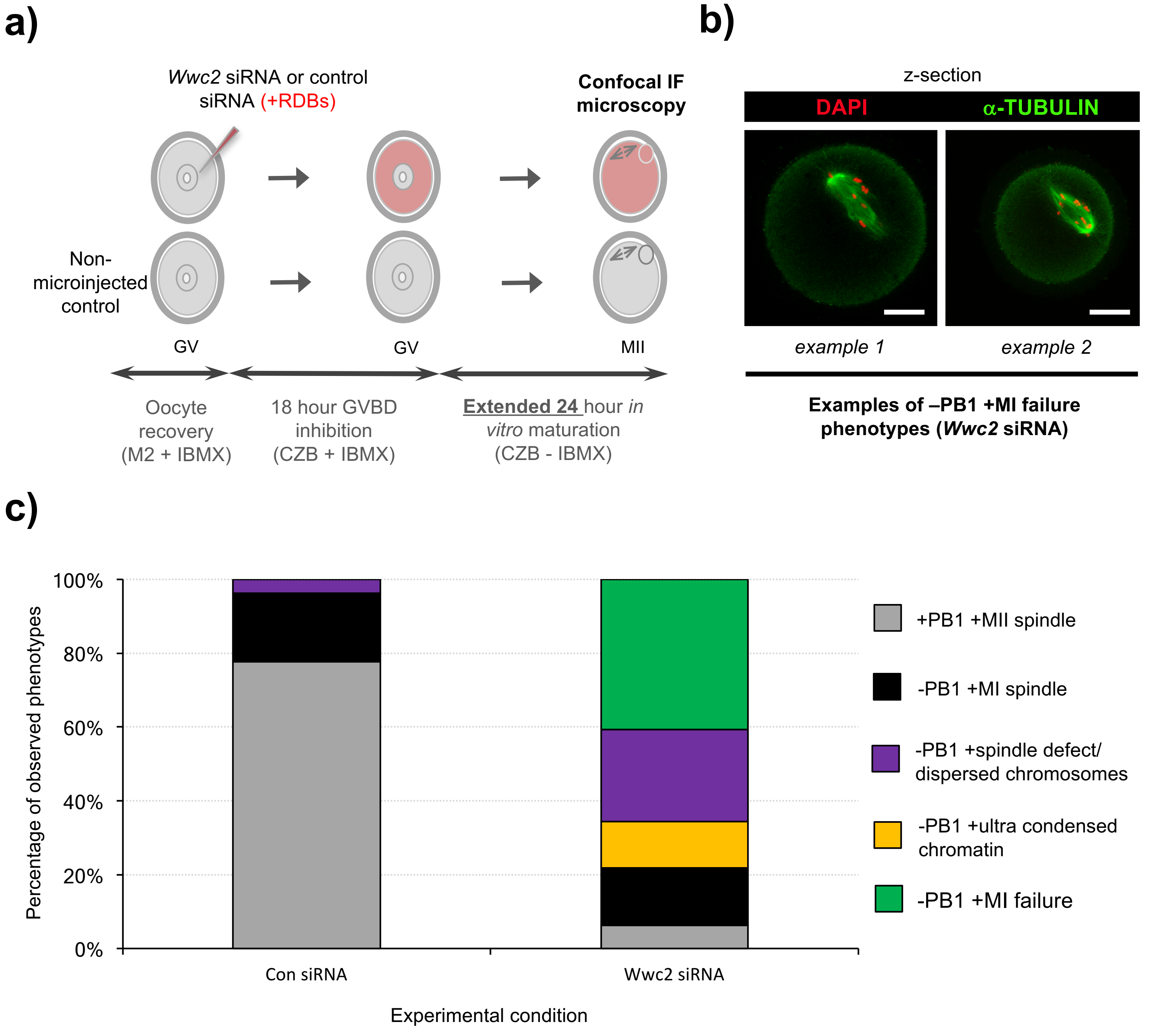
a)
Wwc2 siRNA or control siRNA (+RDBs)
Confocal IF microscopy
Non-microinjected control
GV
GV
MII
Oocyte recovery (M2 + IBMX)
18 hour GVBD inhibition (CZB + IBMX)
Extended 24 hour in vitro maturation (CZB - IBMX)
b)
z-section
DAPI
α-TUBULIN
example 1
example 2
Examples of –PB1 +MI failure phenotypes (Wwc2 siRNA)
c)
100%
80%
60%
40%
20%
0%
Percentage of observed phenotypes
Con siRNA
Wwc2 siRNA
Experimental condition
+PB1 +MII spindle
-PB1 +MI spindle
-PB1 +spindle defect/ dispersed chromosomes
-PB1 +ultra condensed chromatin
-PB1 +MI failure

**Fig. S3**

**Supplementary figure S3: Meiotic *in vitro* maturation phenotypes of *Wwc2* KD GV oocytes are persistent (up to 24 hours post-IBMX) a)** Experimental schema of *Wwc2* transcript KD in GV oocytes by microinjected *Wwc2* siRNA (plus control siRNA groups), 18 hour incubation in IMBX containing media (preventing GVBD) and extended 24 hour IVM (minus IMBX treatment), followed by IF and confocal microscopic analysis; co-microinjected RDBs were used as injection control marker. **b)** Exemplar single z-section confocal micrographs (two) of *Wwc2* siRNA induced and extended (24 hours) IVM phenotypes, classified as "-PB1 +MI failure". DNA is counter-stained with DAPI (pseudo-coloured red) and microtubules IF stained with anti-α-Tubulin antibodies (green); scale bar = 20μm. **c)** Chart detailing successful IVM maturation frequencies of control siRNA (Con. siRNA) and microinjected *Wwc2* siRNA oocytes, to the MII stage or preceding phenotypic stages, as defined/illustrated in Fig. 4 or here (MI failure - panel b). Supplementary tables ST15 summarise statistical analysis and individual oocyte data in each experimental group.

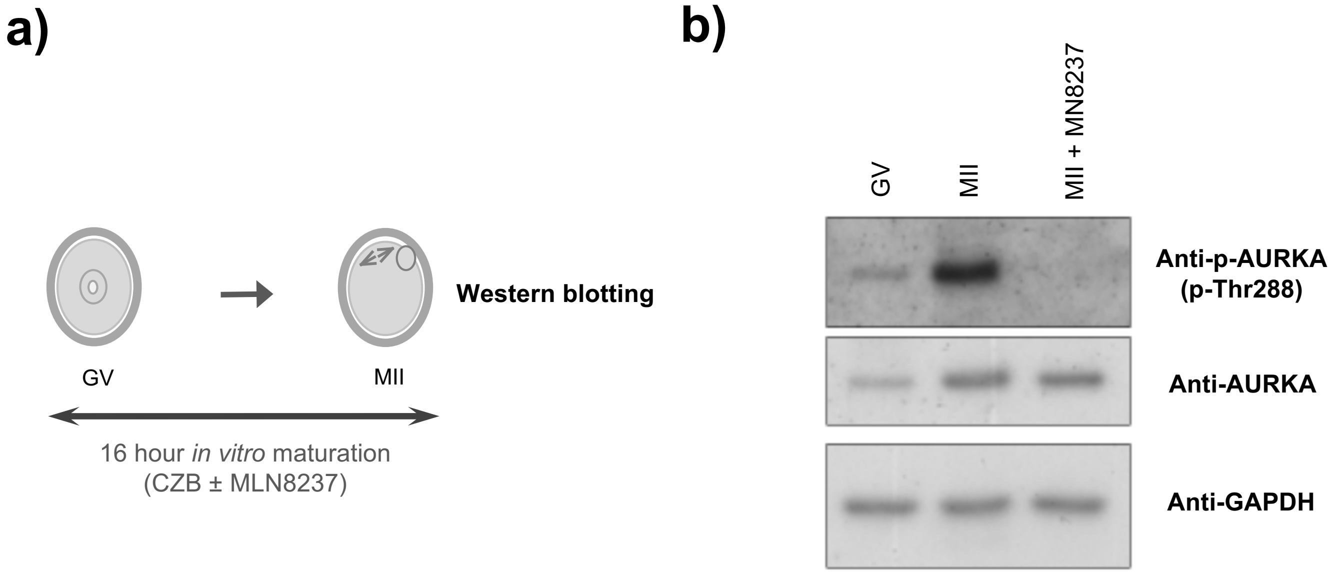
a)
GV
MII
Western blotting
16 hour in vitro maturation
(CZB ± MLN8237)
b)
GV
MII
MII + MN8237
Anti-p-AURKA
(p-Thr288)
Anti-AURKA
Anti-GAPDH

**Fig. S4**

**Supplementary figure S4: anti-p-AURKA (p-Thr288) antibody specificity verification. a)** Experimental strategy to verify specificity of the anti-p-AURKA (p-Thr288) used in western blotting experiments. Recovered GV stage primary oocytes were directly processed for western blotting (as described in the materials and methods) or immediately following *in vitro* maturation (IVM) to the MII arrested stage; in the presence or absence of the confirmed AURKA inhibitor, MLN8237 (1μM: Pollard and Mortimore, 52: 2629-2651, 2009) - MLN8237 blocks auto-phosphorylation/activation of AURKA (Katsha *et al.,* 8:1419-1428, 2014). **b)** Western blots showing activated/phosphorylated p-AURKA (upper panel; probed with anti-p-AURKA/ p-Thr288 primary antibody – Cell Signalling Technology, cs2914), total AURKA (middle panel – BD Biosciences, 610938) or house-keeping control GAPDH (lower panel – Merck, G9545) protein expression levels in freshly recovered GV stage primary oocytes (first lane), IVM MII-arrested secondary oocytes (middle lane) or oocytes subject to IVM in the presence of MLN8237.

**Partial mouse *Wwc2* cDNA sequence (*Wwc2* siRNA target region)**

GACCTGATGCAGAGTCTTGCTAAGCTGCAGGAGCGATTTCATTTGGATCAAAACATGGGC 780
AGTTCCGAGCCAGATCTGAGATCTAGTCCTGTGAATTCTCATCTCTCTCTGTCCAGACAG 840

Complementary *Wwc2* cDNA sequence targeted by *Wwc2* siRNA

| | | | | | | | | | | |
|---|---|---|---|---|---|---|---|---|---|---|
| AGT | TCC | GAG | CCA | GAT | CTG | AGA | TCT | AGT | CCT | GTG |
| S | S | E | P | D | L | R | S | S | P | V |
| 259 | | | | | | | | | | 269 |

Amino acid codon information

| | | | | | | | | | | |
|---|---|---|---|---|---|---|---|---|---|---|
| AGT | TCC | GAG | CCt | GAT | tta | AGg | TCT | tcT | CCT | GTG |
| S | S | E | P | D | L | R | S | S | P | V |
| 259 | | | | | | | | | | 269 |

**Introduced silent mutations in siRNA resistant recombinant *HA-Wwc2* cDNA construct (lowercase)**

**Fig. S5**

**Supplementary figure S5: Information on site directed mutagenesis design to generate siRNA-resistant *HA-Wwc2* 'rescue' construct (microinjected as a mRNA).** Partial cDNA sequence of mouse genome derived *Wwc2* transcript (upper panel), onto which the recognition sequence of the *Wwc2*-specific siRNA used in this study is highlighted (orange: *i.e.* positions 786-813 bp); a plasmid clone of which was generated during this study (*i.e.* pGEM-T-Easy:HA-Wwc2). Expanded representations of this region, detailing the corresponding amino acid codons (delineated by spaces) and the coded residues in the WWC2 protein (as specified using the single letter code), are provided below. The upper expanded depiction describes the endogenous wild-type *Wwc2* sequence and the lower representation details how this was altered by site directed mutagenesis to yield a mRNA construct that, when *in vitro* transcribed, would be insensitive to RNAi mediated degradation when employing our *Wwc2*-specific siRNA (as there would be insufficient base-pair complementarity). The mutated bases are denoted by lower-case font and occur within the would be recognised *Wwc2*-specific siRNA motif but due to the redundancies in the genetic code, do not alter any amino acid coding information. Hence the only distinguishing feature of protein derived from the siRNA resistant mutated recombinant construct/transcript, in relation to its endogenous counterpart, would be the incorporation of the N-terminal HA-epitope tag.

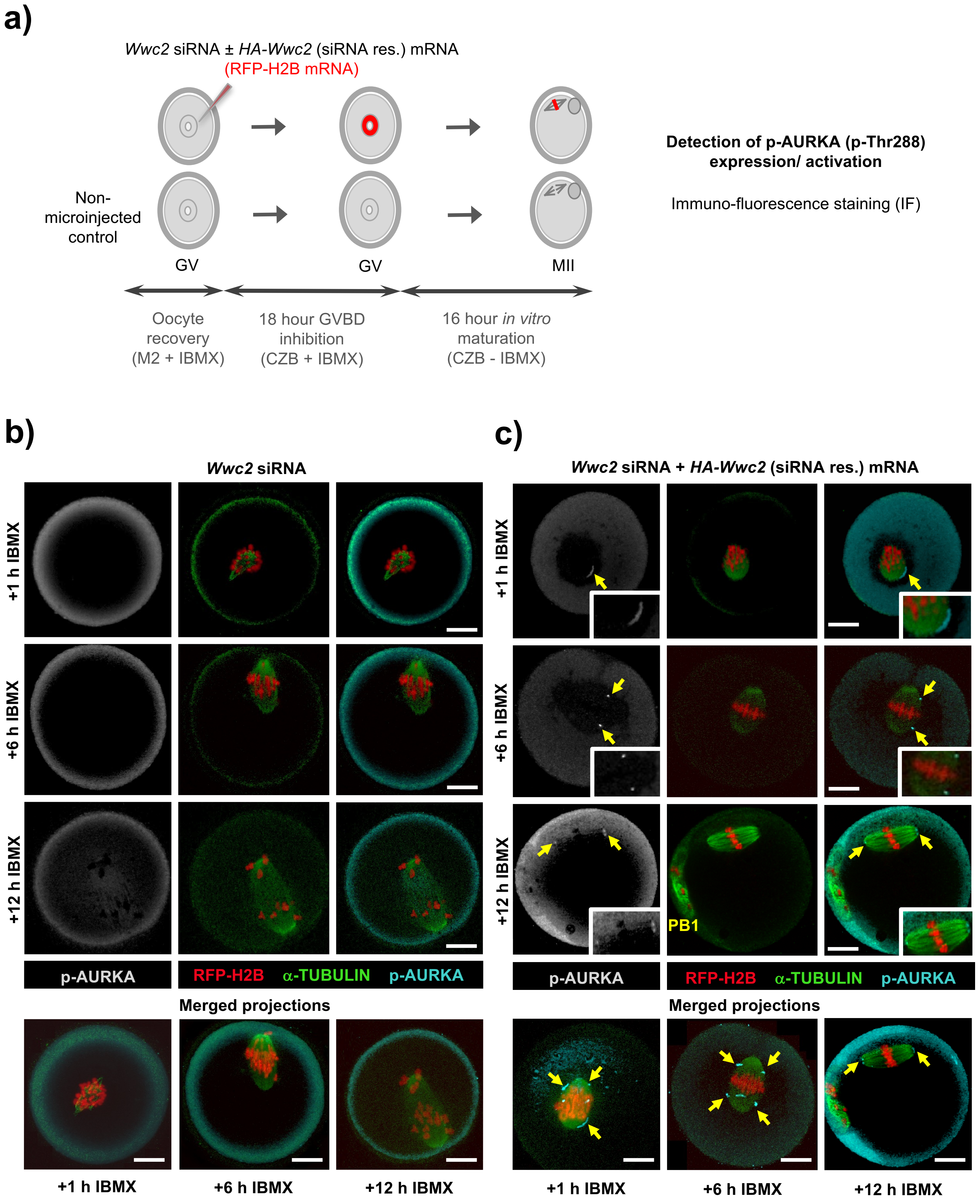
a)
Wwc2 siRNA ± HA-Wwc2 (siRNA res.) mRNA
(RFP-H2B mRNA)
Non-microinjected control
GV
GV
MII
Oocyte recovery (M2 + IBMX)
18 hour GVBD inhibition (CZB + IBMX)
16 hour in vitro maturation (CZB - IBMX)
Detection of p-AURKA (p-Thr288) expression/ activation
Immuno-fluorescence staining (IF)
b)
Wwc2 siRNA
+1 h IBMX
+6 h IBMX
+12 h IBMX
p-AURKA
RFP-H2B α-TUBULIN p-AURKA
Merged projections
+1 h IBMX
+6 h IBMX
+12 h IBMX
c)
Wwc2 siRNA + HA-Wwc2 (siRNA res.) mRNA
+1 h IBMX
+6 h IBMX
+12 h IBMX
PB1
p-AURKA
RFP-H2B α-TUBULIN p-AURKA
Merged projections
+1 h IBMX
+6 h IBMX
+12 h IBMX

**Fig. S6**

**Supplementary figure S6: Phenotypic rescue of *Wwc2* KD induced GV oocyte IVM spindle defects by co-microinjection of siRNA resistant *HA-Wwc2* mRNA. a)** Experimental schema of GV oocyte microinjection conditions; *i.e.* *Wwc2* siRNA alone or *Wwc2* siRNA + *HA-Wwc2* (siRNA resistant) mRNA, each co-microinjected with RFP-H2B mRNA (injection marker). Microinjected oocytes were incubated in IMBX containing media (18 hours - preventing GVBD), subject to IVM (max. 16 hours – media minus IMBX) and processed (at the indicated time-points) for IF assaying of phospho-Aurora (p-AURKA) and α-TUBULIN expression/localisation. **b)** Exemplar z-section confocal micrographs of *Wwc2* siRNA alone injected oocytes after IVM to indicated time-points (post IBMX removal). Detected p-AURKA (greyscale; cyan in merge) and α-TUBULIN (green) expression/localisation, plus the RFP-H2B microinjection reporter (red) are shown. Projected confocal micrographs of colour merged images are depicted at each time-point (lower panels). **c)** As in b) but pertaining to the *Wwc2* siRNA + *HA-Wwc2* (siRNA resistant) mRNA microinjection embryo group. Yellow arrows denote detectable and spindle proximal p-AURKA (p-Thr288) immuno-reactivity (enlarged in the greyscale and merged channel insets); PB1 denotes the extruded first meiotic polar body. In panels b) and c) the scale bar = 20 μm.

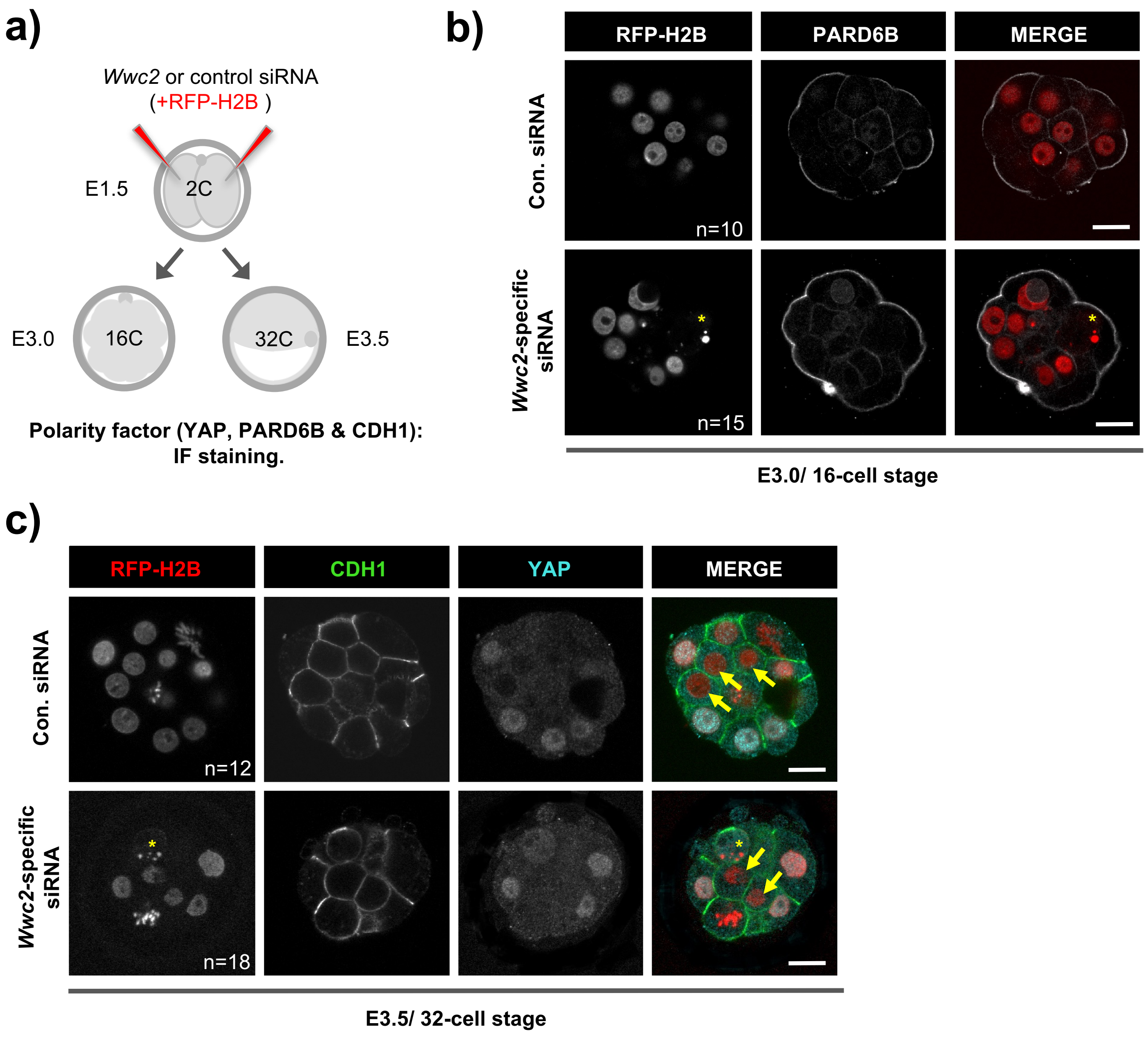
a)
Wwc2 or control siRNA
(+RFP-H2B )
E1.5
2C
E3.0
16C
32C
E3.5
Polarity factor (YAP, PARD6B & CDH1):
IF staining.
b)
RFP-H2B
PARD6B
MERGE
Con. siRNA
n=10
Wwc2-specific siRNA
n=15
E3.0/ 16-cell stage
c)
RFP-H2B
CDH1
YAP
MERGE
Con. siRNA
n=12
Wwc2-specific siRNA
n=18
E3.5/ 32-cell stage

**Fig. S7**

**Supplementary figure S7: *Wwc2* KD does not affect outer-cell apical-basolateral polarisation nor differential inner versus outer-cell Hippo-signalling pathway activity. a)** Experimental strategy to assay apical-basolateral polarity factor (PARD6B and CDH1) and Hippo-pathway effector (YAP) protein expression in 16- or 32-cell stage embryos derived from 2-cell stage embryos co-microinjected in both blastomeres with *Wwc2* (or control) siRNA and mRNA for fluorescent histone H2B fusion protein (RFP-H2B). **b)** Representative confocal z-section micrographs of control and *Wwc2* siRNA microinjected embryos IF stained at the 16-cell stage for PARD6B expression. Individual fluorescent channels are shown in greyscale and merged (in which the RFP-H2B signal is red). **c)** As in b) although embryos are assayed at the 32-cell stage and double-IF stained for CDH1 and YAP (respectively, green and cyan in merged images). Yellow arrows denote inner-cells (from each microinjection group) that do not exhibit nuclear accumulated YAP, as opposed to outer-cell populations. In panels b) and c) yellow asterisk denotes the existence of phenotypically characteristic micronuclei in the *Wwc2* siRNA microinjected embryo groups. Additionally, the number of assayed embryos per group is indicated and the scale bar equals 20μm.

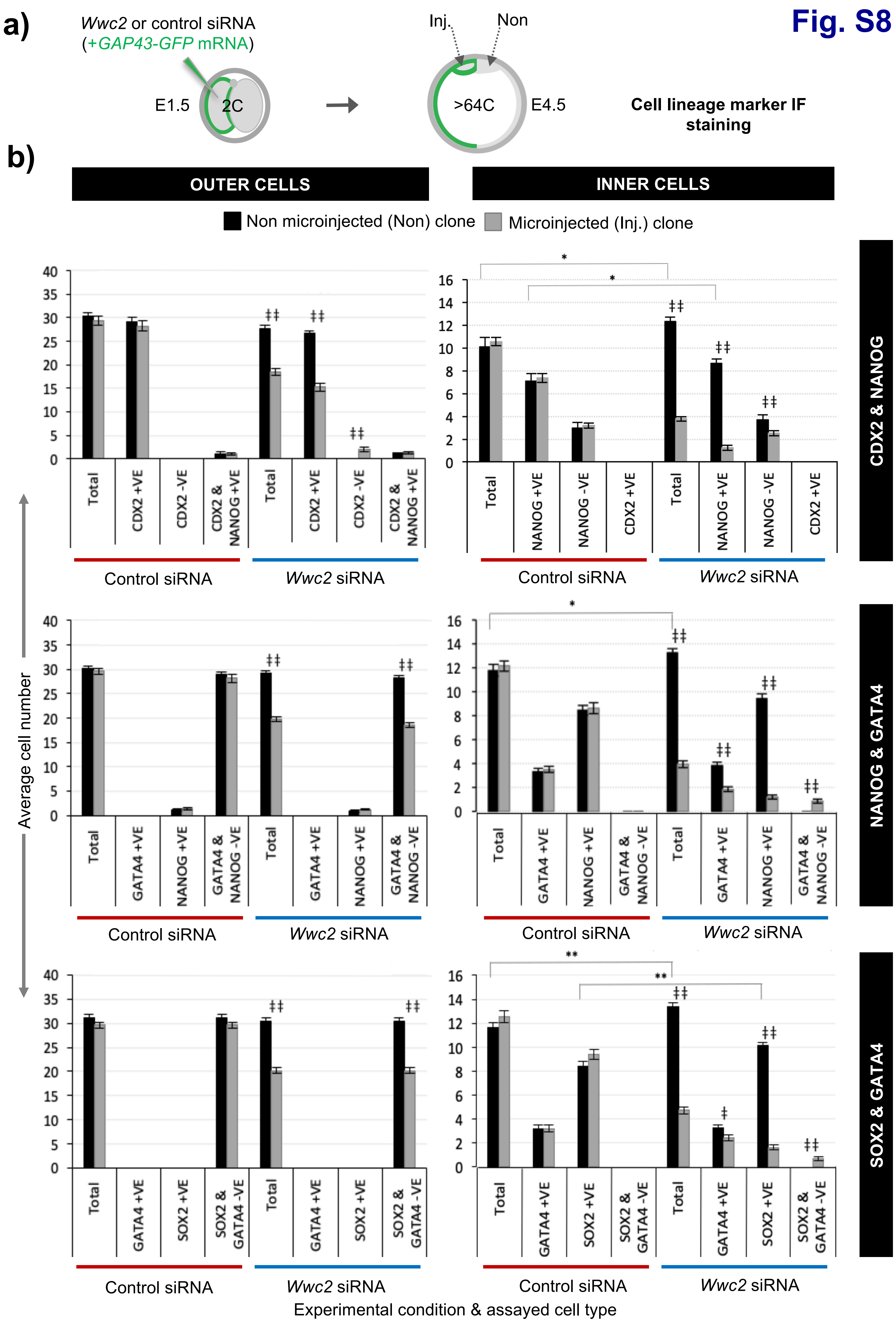
Fig. S8
a)
Wwc2 or control siRNA
(+GAP43-GFP mRNA)
E1.5
2C
Inj.
Non
>64C
E4.5
Cell lineage marker IF staining
b)
OUTER CELLS
INNER CELLS
Non microinjected (Non) clone
Microinjected (Inj.) clone
CDX2 & NANOG
NANOG & GATA4
SOX2 & GATA4
Average cell number
Total
CDX2 +VE
CDX2 -VE
CDX2 & NANOG +VE
NANOG +VE
NANOG -VE
GATA4 +VE
GATA4 & NANOG -VE
SOX2 +VE
SOX2 & GATA4 -VE
Control siRNA
Wwc2 siRNA
Experimental condition & assayed cell type

**Supplementary figure S8: Overall cell lineage formation in clonal *Wwc2* gene KD at the late blastocyst stage (raw average number data format). a)** Experimental strategy for *Wwc2* expression KD in marked clones (representing 50% of the embryo) by co-microinjection (in one blastomere of 2-cell stage embryos) of *Wwc2* siRNA and GAP43-GFP mRNA (stable plasma membrane fluorescent injection marker – Inj.) and an IF based assay of clonal contribution (Inj vs. Non) to blastocyst protein lineage marker expressing cells (TE; CDX2, PrE; GATA4, EPI; NANOG or SOX2; assayed in combination – see below), plus outer and inner-cell embryo populations, at the late blastocyst (>64-cell) stage (versus similar control siRNA microinjections). **b)** The average number of cells contributing to populations expressing, co-expressing or not expressing the stated combinations of late blastocyst lineage marker proteins, their designated clonal origin (Inj vs. Non) and relative distribution between outer and ICM compartments in both control and *Wwc2* siRNA microinjected embryos; errors represent s.e.m. and 2-tailed student t-test derived statistical significance between the control and *Wwc2* knockdown groups (asterisks – note for simplicity only potential regulative differences between the non-microinjected clones are shown; see supplementary tables for full statistical summary) or clones (Inj. vs Non) within a group (double crosses), are highlighted with statistical confidence intervals of $p<0.05$ and $p<0.005$ (denoted by one or two significance markers, respectively). Supplementary tables ST15-ST17 summarise the statistical analysis and individual embryo data in each experimental group.

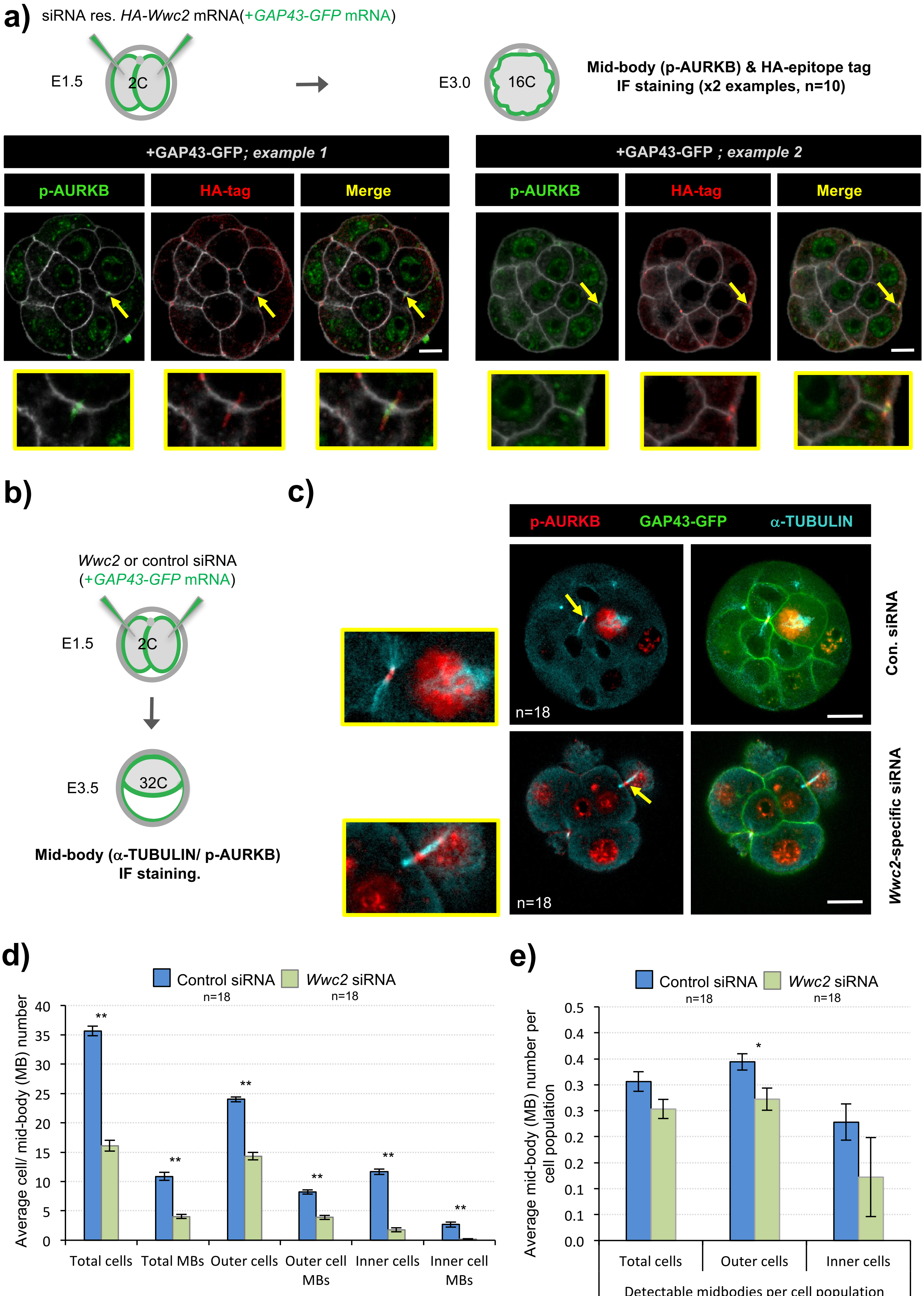
a)
siRNA res. HA-Wwc2 mRNA(+GAP43-GFP mRNA)
E1.5
2C
E3.0
16C
Mid-body (p-AURKB) & HA-epitope tag IF staining (x2 examples, n=10)
+GAP43-GFP; example 1
+GAP43-GFP ; example 2
p-AURKB
HA-tag
Merge
b)
Wwc2 or control siRNA (+GAP43-GFP mRNA)
E1.5
2C
E3.5
32C
Mid-body (α-TUBULIN/ p-AURKB) IF staining.
c)
p-AURKB
GAP43-GFP
α-TUBULIN
n=18
Con. siRNA
Wwc2-specific siRNA
d)
Control siRNA n=18
Wwc2 siRNA n=18
Average cell/ mid-body (MB) number
Total cells
Total MBs
Outer cells
Outer cell MBs
Inner cells
Inner cell MBs
Assayed cell/ mid-body (MB) population
e)
Average mid-body (MB) number per cell population
Total cells
Outer cells
Inner cells
Detectable midbodies per cell population

Fig. S9

**Supplementary figure S9: Global *Wwc2* KD does not significantly alter the number of detectable mid-bodies by the 32-cell stage. a)** Experimental approach to assay co-localisation of recombinant siRNA resistant HA-epitope tagged WWC2 protein with the mitotic mid-body marker phospho-Aurora kinase B (p-AURKB) at the 16-cell stage (upper). Briefly, recovered 2-cell stage embryos were co-microinjected in both blastomeres with recombinant mRNAs encoding siRNA resistant HA-WWC2 and GAP43-GFP (injection marker), cultured to the 16-cell stage and IF stained with antibodies against p-AURKB and HA-epitope tag. Exemplar micrographs (selected single z-sections) of two assayed embryos (lower panels) detail p-AURKB (green), HA-epitope (red) and merged channels; yellow arrows highlight mid-body structures, with magnified images of the regions of interest provided in the lower insets (note, localisation of HA-WWC2 on mid-body proximal microtubules that does not perfectly co-localise with the *de facto* mid-body marker, pAURKB). Scale bar = 20 μm. **b)** Experimental strategy to assay the number of mid-bodies in *Wwc2* KD embryos by co-microinjecting *Wwc2* siRNA (or control siRNA) and GAP43-GFP mRNA (stable plasma membrane fluorescent injection marker) in both cells at the 2-cell stage, fixing after culture to the 32-cell stage and IF staining for the mid-body marker proteins α-TUBULIN and p-AURKB. **c)** Exemplar confocal z-section micrographs of composite IF stained 32-cell stage embryos; p-AURKB (red), α-TUBULIN (cyan) and GAP43-GFP (injection marker – green). Yellow arrows denote persistent mid-bodies characteristic of cleavage stage mouse embryos (zoomed images provided; yellow boxes – left). Number of embryos in each microinjection group provided; scale bars equal 20 μm). **d)** Average number of cells and detectable mid-bodies (in total or either outer or inner-cell populations; note, inner-cell populations defined by mid-body association with at least one inner-cell) per 32-cell stage embryo after microinjection of control (blue bars) or *Wwc2* (green bars) siRNA. **e)** The average number of mid-bodies observed in each stated microinjection group and cell type category, as normalised to the average cell number for that category (*i.e.* total, outer and inner-cells). In panels d) and e) chart error bars represent s.e.m. and 2-tailed student t-test derived statistical significance between the control and *Wwc2* knockdown groups indicated (** $p<0.005$). Supplementary tables ST20 summarise statistical analysis and individual embryo data in each experimental group.

**Human & Mouse KIBRA & WWC2 protein sequence alignment; consensus AURK phosphorylation motifs**

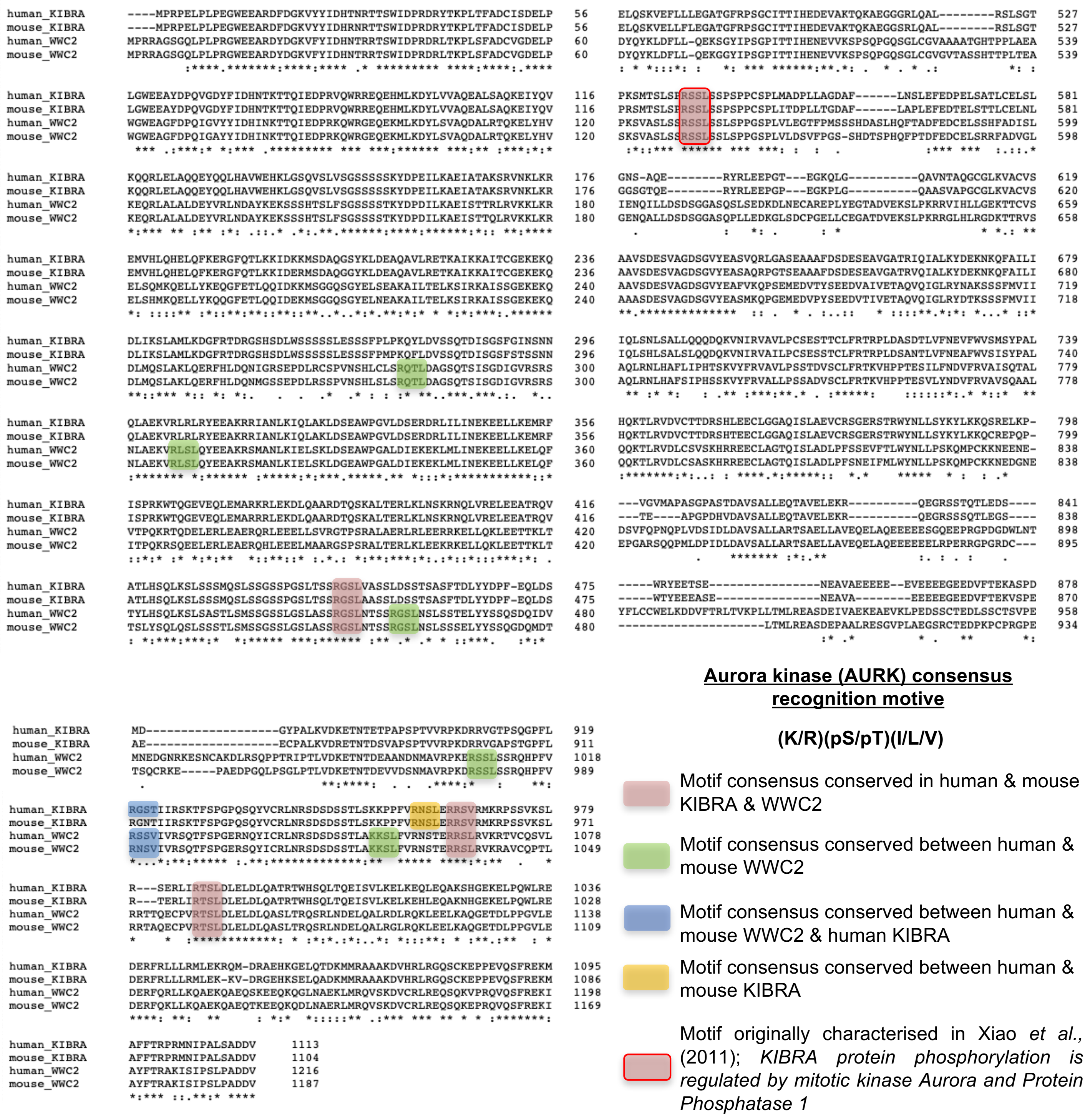


### Fig. S10

**Supplementary figure S10: Primary amino acid sequence alignment of human and mouse KIBRA and WWC2 proteins; with highlighted Aurora kinase consensus phosphorylation motifs.** Amino acid sequence alignments are shown and sequences conforming to the Aurora kinase (AURK) consensus motif [(K/R)(pS/pT)(I/L/V)] – where pS/pT denote the residue targeted for phosphorylation, highlighted according to the legend provided. All alignments were conducted using default setting of the online Clustal-Ω multiple sequence alignment tool; asterisks (*) denote perfectly aligned and identical amino acids, whereas colons (:) detail aligned conservative amino acid substitutions.

**B) SUPPLMENTARY METHODS TABLES: SM1 – SM6**

***Wwc2* is a novel cell division regulator during preimplantation mouse embryo lineage formation and oogenesis.**

Virnicchi *et al.,* 2020

**Supplementary Methods Tables SM1**

| Microinjected RNAi constructs (Virnicchi *et al.,* 2019) | | | | |
|---|---|---|---|---|
| **Type** | **Gene Target** | **Souce (supplier)** | **Cat. No.** | **Microinjection conc.** |
| dsRNA | *Kibra* | in house synthesised | n/a | 200 ng/μl |
| dsRNA | *Wwc2* | in house synthesised | n/a | 200 ng/μl |
| dsRNA | *EGFP* | in house synthesised | n/a | 200 ng/μl |
| siRNA | mouse genome neg. control | Qiagen (Murine Allstars neg. control siRNA) | SI03650318 | 10 μM |
| siRNA | *Wwc2* | ThermoFisher Scientific (Silencer Select) | 4390771: s78812 | 10 μM |

| Microinjected recombinant & IVT derived mRNA constructs (Virnicchi *et al.,* 2019) | | | | |
|---|---|---|---|---|
| **Type** | **Gene Target** | **Souce (supplier)** | **Cat. No.** | **Microinjection conc.** |
| mRNA | *GAP43-GFP* | in house synthesised | n/a | 60 ng/μl |
| mRNA | *H2B-RFP* | in house synthesised | n/a | 40 ng/μl |
| mRNA | *Wwc2-HA (siRNA res.)* | in house synthesised | n/a | 85 ng/μl |

**Supplementary Methods Table SM2**

| Oligonucleotide PCR primer sequence for dsRNA template generation (Virnicchi *et al.,* 2019) | | |
|---|---|---|
| **dsRNA gene target** | **T7 RNA polymerase promoter sequence (lowever case) linked gene specific primer senquence** | |
| | **Sense (5'-3')** | **Anti-sense (5'-3')** |
| Kibra | taatacgactcactatagggGACTTCGACGGCAAGGTCTA | taatacgactcactatagggTCCGACCTGTGGGTCGTAT |
| Wwc2 | taatacgactcactatagggGAGCTGCTGTGTGAGGGC | taatacgactcactatagggAGAAGGCATGAAGATTTCGG |
| Egfp | taatacgactcactatagggAGAGTACAAATTTTCTGTCAGTGGAGAGG | taatacgactcactatagggAGATGTATAGTTCATCCATGCCATGTGTA |

**Supplementary Methods Table SM3**

| Recombinant plasmid derivation information (Virnicchi *et al.,* 2019) | | | | | |
|---|---|---|---|---|---|
| **Plasmid name** | **Backbone vector** | **Insert cDNA(species derived from)** | **ligonucleotide primer pair sequences (used for insert generation; *if applicable* ) + introduced restriction enzyme recognition sites for sub cloning (lower case & under** | | **Notes** |
| | | | **Sense (5'-3') - introduced KOZAK & HA-epitope tag sequences highlighted** | **Anti-sense (5'-3')** | |
| pRN3-C-term-RFP-Hist1h2bb | pRN3-insert-RFP | Hist1h2bb, histonme H2B (mouse) | GACTATgctagcCCAGAGCCTTCTAAGTCTGCAC | GACTATgctagcCTTGGAGCTGGTGTACTTGGTGA | insert generating PCR oligos lack 'start' and 'stop' codons, as these are encoded in the vector (allowing N & C-terminal RFP fusions) |
| pGEM-T-Easy-N-term-HA-wildtype-Wwc2 | pGEM®-T-Easy | Wwc2, wildtype (mouse) | GACTATactagtGCCACCATGggctacccatacgatgttcctgactatgctCCTAGGAGGGCCGGGAGC (*SpeI*) | GACTATgcggccgcTCACACGTCGTCAGCGGGC (*NotI*) | Insert generated by PCR and 'TA' cloned into pGEM®-T-Easy (and sequence verified) |
| pGEM-T-Easy-N-term-HA-siRNAres-Wwc2 | pGEM®-T-Easy | Wwc2, siRNA resistant (mouse) | **n/a (**Wwc2 siRNA recognition sequence mutated *in situ* by SDM; commercial serive - EuorFins) | | siRNA resistant mutant derived by commercial SDM (siRNA recognition motif mutated, codon info retained - see Fig. S5) |
| pRN3P-N-term-HA-siRNAres-Wwc2 | pRN3P | Wwc2, siRNA resistant (mouse) | **n/a** (insert was subcloned from pGEM-T-Easy-N-term-HA-siRNAres-Wwc2, using previously introduced *SpeI* & *NotI* restriction sites) | | Cloned directlyinto pRN3P from a Spei & NotI digest of pGEM-T-Easy-N-term-HA-siRNAres-Wwc2 |

**Supplementary Methods Table SM4**

| Q-RT-PCR Oligonucleotide primer sequences (Virnicchi *et al.,* 2019) | | |
|---|---|---|
| Target gene mRNA/ cDNA | Q-RT-PCR oligonucleotide primer sequences | |
| | Sense (5'-3') | Anti-sense (5'-3') |
| *Tbp* (TATA-binding protein) | GAAGAACAATCCAGACTAGCAGCA | CCTTATAGGGAACTTCACATCACAG |
| *Kibra* | ATGATGAGAGCTGCTGCCAAGG | ATCCGAGGCCGGGTGAAAAATG |
| *Wwc2* | TGCTGGAGGACGAGAGATTC | GAGACCTGCCTCATCAACCT |

**Supplementary Methods Tables SM5**

| Primary antibody (immuno-fluorescence staining) information (Virnicchi *et al.,* 2019) | | | | | | |
|---|---|---|---|---|---|---|
| **#** | **Antigen** | **cat. no.** | **Supplier** | **Species raised in & clonicity** | **Dilution used** | **Secondary antibody combination used (see below)** |
| 1 | CDH1 | 3195 | Cell Signalling Technology | rabbit, polyclonal | 1 in 500 | E |
| 2 | CDX2 | MU392A-UC | BioGenex | mouse, monoclonal | 1 in 200 | D |
| 3 | GATA4 | sc-9053 | Santa Cruz | rabbit, polyclonal | 1 in 100 | E |
| 4 | HA-TAG | ab9134 | Abcam | goat, polyclonal | 1 in 200 | H |
| 5 | NANOG | 14-5761 | Affymetrix/eBioscience | rat, monoclonal | 1 in 100 | B |
| 6 | PARD6B | sc-67393 | Santa Cruz | rabbit, polyclonal | 1 in100 | G |
| 7 | phospho-AURKA (Thr288) | NB100-2371 | Novus Biologicals | rabbit, polyclonal | 1 in 200 | E |
| 8 | phospho-AURKB (Thr232) | 600-401-677S | Rockland Antibodies | rabbit, polyclonal | 1 in 200 | F |
| 9 | SOX2 | sc-365823 | Santa Cruz | mouse, monoclonal | 1 in 200 | D |
| 10 | TUBULIN (alpha) | A11126 | ThermoFisher Scientific | mouse, monoclonal | 1 in 200 | A/C |
| 11 | YAP1 | sc-101199 | Santa Cruz | mouse, monoclonal | 1 in100 | A |

| Secondary antibody (immuno-fluorescence staining) information (Virnicchi *et al.,* 2019) | | | | | | |
|---|---|---|---|---|---|---|
| **#** | **Species of antibody targetted** | **cat. no.** | **Supplier** | **Species raised in & fluorophore** | **Dilution used** | **Primary antibody combination used (see above)** |
| A | mouse | 715-605-150 | Jackson Immuno Research Inc. | donkey, Alexa647 | 1 in 500 | 10/11 |
| B | rat | 715-096-150 | Jackson Immuno Research Inc. | donkey, FITC | 1 in 400 | 5 |
| C | mouse | A-21202 | ThermoFisher Scientific | donkey Alexa488 | 1 in 500 | 10 |
| D | mouse | A-31570 | ThermoFisher Scientific | donkey Alexa555 | 1 in 500 | 2/9 |
| E | rabbit | A-21206 | ThermoFisher Scientific | donkey, Alexa488 | 1 in 500 | 1/3/7 |
| F | rabbit | A-31572 | ThermoFisher Scientific | donkey Alexa555 | 1 in 500 | 8/9 |
| G | rabbit | A-31573 | ThermoFisher Scientific | donkey, Alexa647 | 1 in 500 | 6 |
| H | goat | A-21432 | ThermoFisher Scientific | donkey, Alexa555 | 1 in 500 | 4 |

**Supplementary Methods Tables SM6**

| Primary antibody (western blot) information (Virnicchi *et al.,* 2019) | | | | | | |
|---|---|---|---|---|---|---|
| **#** | **Antigen** | **cat. no.** | **Supplier** | **Species raised in & clonicity** | **Dilution used** | **Secondary antibody combination used (see below)** |
| 1 | phospho-AURKA (Thr288) | 2914 | Cell Signalling Technology | rabbit, monoclonal | 1 in 2,000 | A |
| 2 | GAPDH | G9545 | Merck (Sigma-Aldrich) | rabbit, polyclonal | 1 in 20,000 | A |

| Secondary HRP-CONJUGATED antibody (western blot) information (Virnicchi *et al.,* 2019) | | | | | | |
|---|---|---|---|---|---|---|
| **#** | **Species of antibody targetted** | **cat. no.** | **Supplier** | **Species raised in** | **Dilution used** | **Primary antibody combination used (see above)** |
| A | rabbit | 711-035-152 | Jackson Immuno Research Inc. | donkey, polyclonal | 1 in 10,000 | 1/2 |

**C) SUPPLMENTARY TABLES (individual embryo cell counts & oocyte phenotypes + statistics): ST1 – ST26**

***Wwc2* is a novel cell division regulator during preimplantation mouse embryo lineage formation and oogenesis.**

Virnicchi *et al.,* 2020

Supplementary Tables ST1

| Control (GFP) dsRNA (300ng/μl), 1in2 micro-injection, >64-cell (E4.5) stage | | | |
|---|---|---|---|
| # | Total cell number per embryo | Cells in injected clone | Cells in non-injected clone |
| 1 | 63 | not assayed | not assayed |
| 2 | 99 | not assayed | not assayed |
| 3 | 55 | not assayed | not assayed |
| 4 | 83 | not assayed | not assayed |
| 5 | 102 | not assayed | not assayed |
| 6 | 76 | not assayed | not assayed |
| 7 | 83 | not assayed | not assayed |
| 8 | 83 | 38 | 45 |
| 9 | 75 | 37 | 38 |
| 10 | 91 | 44 | 47 |
| 11 | 82 | 41 | 41 |
| 12 | 71 | 38 | 33 |
| 13 | 76 | 39 | 37 |
| 14 | 89 | 46 | 43 |
| 15 | 93 | 45 | 48 |
| 16 | 68 | 27 | 41 |
| 17 | 86 | 44 | 42 |
| TOTAL | 1375 | 399 | 415 |
| AVERAGE | 80.9 | 39.9 | 41.5 |
| SEM | 3.0 | 1.8 | 1.5 |
| Stat. sig. (inter-clone) ‡p<0.05, ‡‡p<0.005 | | | |
| p-value (2-tailed students t-test) | | 4.94E-01 | |

| Wwc2-specific (CDS) dsRNA (300ng/μl), 1in2 micro-injection, >64-cell (E4.5) stage | | | |
|---|---|---|---|
| # | Total cell number per embryo | Cells in injected clone | Cells in non-injected clone |
| 1 | 77 | not assayed | not assayed |
| 2 | 80 | not assayed | not assayed |
| 3 | 55 | not assayed | not assayed |
| 4 | 41 | not assayed | not assayed |
| 5 | 33 | not assayed | not assayed |
| 6 | 44 | not assayed | not assayed |
| 7 | 43 | not assayed | not assayed |
| 8 | 23 | not assayed | not assayed |
| 9 | 42 | not assayed | not assayed |
| 10 | 45 | not assayed | not assayed |
| TOTAL | 483 | n/a | n/a |
| AVERAGE | 48.3 | n/a | n/a |
| SEM | 5.7 | n/a | n/a |
| Stat. sig. (inter-clone) ‡p<0.05, ‡‡p<0.005 | | | |
| p-value (2-tailed students t-test) | | n/a | |
| Stat. sig. (exp. vs. con embryo) *p<0.05, **p<0.005 | ** | | |
| p-value (2-tailed students t-test) | 8.35E-06 | n/a | n/a |

| Kibra-specific (CDS) dsRNA (300ng/μl), 1in2 micro-injection, >64-cell (E4.5) stage | | | |
|---|---|---|---|
| # | Total cell number per embryo | Cells in injected clone | Cells in non-injected clone |
| 1 | 91 | 45 | 46 |
| 2 | 89 | 45 | 44 |
| 3 | 80 | 48 | 32 |
| 4 | 83 | 35 | 48 |
| 5 | 96 | 57 | 39 |
| 6 | 66 | 19 | 47 |
| 7 | 63 | 42 | 21 |
| 8 | 88 | 46 | 42 |
| 9 | 90 | 30 | 60 |
| TOTAL | 746 | 367 | 379 |
| AVERAGE | 82.9 | 40.8 | 42.1 |
| SEM | 3.8 | 3.7 | 3.6 |
| Stat. sig. (inter-clone) ‡p<0.05, ‡‡p<0.005 | | | |
| p-value (2-tailed students t-test) | | 8.01E-01 | |
| Stat. sig. (exp. vs. con embryo) *p<0.05, **p<0.005 | | | |
| p-value (2-tailed students t-test) | 6.91E-01 | 8.29E-01 | 8.73E-01 |
| Stat. sig. (exp. vs. Wwc2 (CDS) dsRNA embryo) §p<0.05, §§p<0.005 | §§ | | |
| p-value (2-tailed students t-test) | 1.23E-04 | n/a | n/a |

| Kibra & Wwc2-specific (CDS) dsRNA (300ng/μl), 1in2 micro-injection, >64-cell (E4.5) stage | | | |
|---|---|---|---|
| # | Total cell number per embryo | Cells in injected clone | Cells in non-injected clone |
| 1 | 55 | 17 | 38 |
| 2 | 42 | 14 | 28 |
| 3 | 59 | 15 | 44 |
| 4 | 58 | 15 | 43 |
| 5 | 62 | 21 | 41 |
| TOTAL | 276 | 82 | 194 |
| AVERAGE | 55.2 | 16.4 | 38.8 |
| SEM | 3.5 | 1.2 | 2.9 |
| Stat. sig. (inter-clone) ‡p<0.05, ‡‡p<0.005 | | ‡‡ | |
| p-value (2-tailed students t-test) | | 1.00E-04 | |
| Stat. sig. (exp. vs. con embryo) *p<0.05, **p<0.005 | ** | ** | |
| p-value (2-tailed students t-test) | 3.19E-04 | 8.20E-07 | 3.66E-01 |
| Stat. sig. (exp. vs. Wwc2 (CDS) dsRNA embryo) §p<0.05, §§p<0.005 | | | |
| p-value (2-tailed students t-test) | 4.32E-01 | n/a | n/a |
| Stat. sig. (exp. vs. Kibra (CDS) dsRNA embryo) +p<0.05, ++p<0.005 | ++ | ++ | |
| p-value (2-tailed students t-test) | 4.34E-04 | 5.09E-04 | 5.51E-01 |

**Supplementary Tables ST2**

| Control (GFP) dsRNA (300ng/µl), 1in2 micro-injection, 16-cell (E3.0) stage | | | | | | | | | |
|---|---|---|---|---|---|---|---|---|---|
| # | Total cell number per embryo | Cells in injected clone | Cells in non-injected clone | Total OUTER cells per embryo | OUTER CELLS | | Total INNER cells per embryo | INNER CELLS | |
| | | | | | Injected clone | Non-Injected clone | | Injected clone | Non-Injected clone |
| **1** | 17 | 8 | 9 | 9 | 4 | 5 | 8 | 4 | 4 |
| **2** | 16 | 8 | 8 | 10 | 5 | 5 | 6 | 3 | 3 |
| **3** | 16 | 8 | 8 | 12 | 6 | 6 | 4 | 2 | 2 |
| **4** | 17 | 9 | 8 | 10 | 5 | 5 | 7 | 4 | 3 |
| **5** | 16 | 8 | 8 | 11 | 5 | 6 | 5 | 3 | 2 |
| **6** | 16 | 8 | 8 | 11 | 6 | 5 | 5 | 2 | 3 |
| **7** | 15 | 7 | 8 | 10 | 5 | 5 | 5 | 2 | 3 |
| **8** | 16 | 8 | 8 | 11 | 5 | 6 | 5 | 3 | 2 |
| **TOTAL** | **129** | **64** | **65** | **84** | **41** | **43** | **45** | **23** | **22** |
| **AVERAGE** | **16.1** | **8.0** | **8.1** | **10.5** | **5.1** | **5.4** | **5.6** | **2.9** | **2.8** |
| SEM | **0.2** | **0.2** | **0.1** | **0.3** | **0.2** | **0.2** | **0.5** | **0.3** | **0.3** |
| Stat. sig. (inter-clone) ‡p<0.05, ‡‡p<0.005 | | | | | | | | | |
| p-value (2-tailed students t-test) | | 5.90E-01 | | | 4.05E-01 | | | 7.51E-01 | |

| Wwc2-specific (CDS) dsRNA (300ng/µl), 1in2 micro-injection, 16-cell (E3.0) stage | | | | | | | | | |
|---|---|---|---|---|---|---|---|---|---|
| # | Total cell number per embryo | Cells in injected clone | Cells in non-injected clone | Total OUTER cells per embryo | OUTER CELLS | | Total INNER cells per embryo | INNER CELLS | |
| | | | | | Injected clone | Non-Injected clone | | Injected clone | Non-Injected clone |
| **1** | 12 | 4 | 8 | 7 | 2 | 5 | 5 | 2 | 3 |
| **2** | 12 | 4 | 8 | 8 | 3 | 5 | 4 | 1 | 3 |
| **3** | 14 | 6 | 8 | 10 | 6 | 4 | 4 | 0 | 4 |
| **4** | 16 | 8 | 8 | 11 | 6 | 5 | 5 | 2 | 3 |
| **5** | 12 | 4 | 8 | 8 | 4 | 4 | 4 | 0 | 4 |
| **6** | 13 | 4 | 9 | 8 | 4 | 4 | 5 | 0 | 5 |
| **7** | 16 | 8 | 8 | 10 | 5 | 5 | 6 | 3 | 3 |
| **8** | 12 | 4 | 8 | 9 | 4 | 5 | 3 | 0 | 3 |
| **9** | 16 | 8 | 8 | 13 | 7 | 6 | 3 | 1 | 2 |
| **10** | 19 | 10 | 9 | 15 | 8 | 7 | 4 | 2 | 2 |
| **TOTAL** | **142** | **60** | **82** | **99** | **49** | **50** | **43** | **11** | **32** |
| **AVERAGE** | **14.2** | **6.0** | **8.2** | **9.9** | **4.9** | **5.0** | **4.3** | **1.1** | **3.2** |
| SEM | **0.8** | **0.7** | **0.1** | **0.8** | **0.6** | **0.3** | **0.3** | **0.3** | **0.3** |
| Stat. sig. (inter-clone) ‡p<0.05, ‡‡p<0.005 | | ‡ | | | | | | ‡‡ | |
| p-value (2-tailed students t-test) | | 8.32E-03 | | | 3.70E-01 | | | 2.07E-04 | |
| Stat. sig. (exp. vs. con embryo) *p<0.05, **p<0.005 | * | * | | | | | * | ** | |
| p-value (2-tailed students t-test) | 4.63E-02 | 2.98E-02 | 6.93E-01 | 5.33E-01 | 7.48E-01 | 3.29E-01 | 2.37E-02 | 1.68E-03 | 2.71E-01 |

Supplementary Tables ST3

| Control siRNA (10µM), 2in2 micro-injection, 4-cell (E2.0) stage | | | | | |
|---|---|---|---|---|---|
| # | Total cell number per embryo | Total number of dividing cells | Number of cells exhibiting an atypical/ unusual phenotype morphology | | |
| | | | Abnormal nuclear morphology (e.g. non-spheroidal or mid-body associated) | Defective cytokinesis (i.e. >x2 equally sized nuclei per cell) | Presence of micronuclei |
| 1 | 4 | 0 | 0 | 0 | 0 |
| 2 | 4 | 0 | 0 | 0 | 0 |
| 3 | 4 | 0 | 0 | 0 | 0 |
| 4 | 4 | 0 | 0 | 0 | 0 |
| 5 | 4 | 0 | 0 | 0 | 0 |
| 6 | 4 | 0 | 0 | 0 | 0 |
| 7 | 4 | 0 | 0 | 0 | 0 |
| 8 | 4 | 0 | 0 | 0 | 0 |
| 9 | 3 | 1 | 0 | 0 | 0 |
| 10 | 4 | 0 | 0 | 0 | 0 |
| 11 | 4 | 0 | 0 | 0 | 0 |
| 12 | 4 | 0 | 0 | 0 | 0 |
| 13 | 4 | 0 | 0 | 0 | 0 |
| TOTAL | 51 | 1 | 0 | 0 | 0 |
| AVERAGE | 3.9 | 0.1 | 0.0 | 0.0 | 0.0 |
| SEM | 0.1 | 0.08 | 0.0 | 0.0 | 0.0 |

| Wwc2-specific siRNA (10µM), 2in2 micro-injection, 4-cell (E2.0) stage | | | | | |
|---|---|---|---|---|---|
| # | Total cell number per embryo | Total number of dividing cells | Number of cells exhibiting an atypical/ unusual phenotype morphology | | |
| | | | Abnormal nuclear morphology (e.g. non-spheroidal or mid-body associated) | Defective cytokinesis (i.e. >x2 equally sized nuclei per cell) | Presence of micronuclei |
| 1 | 4 | 0 | 0 | 0 | 0 |
| 2 | 4 | 0 | 0 | 0 | 0 |
| 3 | 4 | 0 | 0 | 0 | 0 |
| 4 | 4 | 0 | 0 | 0 | 0 |
| 5 | 4 | 0 | 0 | 0 | 0 |
| 6 | 3 | 1 | 0 | 0 | 0 |
| 7 | 4 | 0 | 0 | 0 | 0 |
| 8 | 4 | 0 | 0 | 0 | 0 |
| 9 | 4 | 0 | 0 | 0 | 0 |
| 10 | 4 | 0 | 0 | 0 | 0 |
| 11 | 4 | 0 | 0 | 0 | 0 |
| 12 | 4 | 0 | 0 | 0 | 0 |
| 13 | 4 | 0 | 0 | 0 | 0 |
| 14 | 4 | 0 | 0 | 0 | 0 |
| TOTAL | 55 | 1 | 0 | 0 | 0 |
| AVERAGE | 3.9 | 0.1 | 0.0 | 0.0 | 0.0 |
| SEM | 0.1 | 0.1 | 0.0 | 0.0 | 0.0 |
| Stat. sig. (exp. vs. con embryo) *p<0.05, **p<0.005 | | | | | |
| p-value (2-tailed students t-test) | 9.59E-01 | 9.59E-01 | n/a | n/a | n/a |

**Supplementary Tables ST4**

| Control siRNA (10μM), 2in2 micro-injection, 8-cell (E2.5) stage | | | | | |
|---|---|---|---|---|---|
| # | Total cell number per embryo | Total number of dividing cells | Number of cells exhibiting an atypical/ unusual phenotype morphology | | |
| | | | Abnormal nuclear morphology (e.g. non-spheroidal or mid-body associated) | Defective cytokinesis (i.e. >x2 equally sized nuclei per cell) | Presence of micronuclei |
| **1** | 6 | 0 | 0 | 0 | 0 |
| **2** | 8 | 0 | 0 | 0 | 0 |
| **3** | 8 | 0 | 0 | 0 | 0 |
| **4** | 8 | 0 | 0 | 0 | 0 |
| **5** | 8 | 0 | 0 | 0 | 0 |
| **6** | 8 | 0 | 0 | 0 | 0 |
| **TOTAL** | **46** | **0** | **0** | **0** | **0** |
| **AVERAGE** | **7.7** | **0.0** | **0.0** | **0.0** | **0.0** |
| SEM | **0.3** | **0.0** | **0.0** | **0.0** | **0.0** |

| Wwc2-specific siRNA (10μM), 2in2 micro-injection, 8-cell (E2.5) stage | | | | | |
|---|---|---|---|---|---|
| # | Total cell number per embryo | Total number of dividing cells | Number of cells exhibiting an atypical/ unusual phenotype morphology | | |
| | | | Abnormal nuclear morphology (e.g. non-spheroidal or mid-body associated) | Defective cytokinesis (i.e. >x2 equally sized nuclei per cell) | Presence of micronuclei |
| **1** | 7 | 1 | 2 | 0 | 0 |
| **2** | 7 | 0 | 0 | 0 | 0 |
| **3** | 6 | 0 | 0 | 0 | 0 |
| **4** | 8 | 0 | 0 | 0 | 0 |
| **5** | 8 | 0 | 3 | 0 | 0 |
| **6** | 8 | 0 | 0 | 0 | 0 |
| **7** | 8 | 0 | 0 | 0 | 0 |
| **TOTAL** | **52** | **1** | **5** | **0** | **0** |
| **AVERAGE** | **7.4** | **0.1** | **0.7** | **0.0** | **0.0** |
| SEM | **0.3** | **0.1** | **0.5** | **0.0** | **0.0** |
| Stat. sig. (exp. vs. con embryo) *p<0.05, **p<0.005 | | | | | |
| p-value (2-tailed students t-test) | 6.04E-01 | 3.77E-01 | 1.93E-01 | n/a | n/a |

Supplementary Tables ST5

| Control siRNA (10μM), 2in2 micro-injection, 16-cell (E3.0) stage | | | | | |
|---|---|---|---|---|---|
| # | Total cell number per embryo | Total number of dividing cells | Number of cells exhibiting an atypical/ unusual phenotype morphology | | |
| | | | Abnormal nuclear morphology (e.g. non-spheroidal or mid-body associated) | Defective cytokinesis (i.e. >x2 equally sized nuclei per cell) | Presence of micronuclei |
| **1** | 15 | | 0 | 0 | 0 |
| **2** | 16 | | 0 | 0 | 0 |
| **3** | 14 | | 0 | 0 | 0 |
| **4** | 16 | | 0 | 0 | 0 |
| **5** | 16 | | 0 | 0 | 0 |
| **6** | 16 | | 0 | 0 | 0 |
| **7** | 19 | | 0 | 0 | 0 |
| **8** | 16 | | 0 | 0 | 0 |
| **9** | 16 | | 0 | 0 | 0 |
| **10** | 17 | | 0 | 0 | 0 |
| **11** | 16 | | 0 | 0 | 0 |
| **12** | 19 | | 0 | 0 | 0 |
| **13** | 19 | | 0 | 0 | 0 |
| **14** | 16 | | 0 | 0 | 0 |
| **15** | 16 | | 0 | 0 | 0 |
| **16** | 19 | | 0 | 0 | 0 |
| **17** | 15 | | 0 | 0 | 0 |
| **18** | 18 | | 0 | 0 | 0 |
| **19** | 17 | | 0 | 0 | 0 |
| **20** | 16 | | 0 | 0 | 0 |
| **21** | 17 | | 0 | 0 | 0 |
| **22** | 27 | | 0 | 0 | 0 |
| **23** | 16 | | 0 | 0 | 0 |
| **24** | 19 | | 0 | 0 | 0 |
| **25** | 31 | | 0 | 0 | 0 |
| **26** | 16 | | 0 | 0 | 0 |
| **27** | 16 | | 0 | 0 | 0 |
| **28** | 14 | | 0 | 0 | 0 |
| **29** | 16 | | 0 | 0 | 0 |
| **30** | 15 | | 0 | 0 | 0 |
| **TOTAL** | **519** | **0** | **0** | **0** | **0** |
| **AVERAGE** | **17.3** | **0.0** | **0.0** | **0.0** | **0.0** |
| SEM | **0.6** | **0.00** | **0.0** | **0.0** | **0.0** |

| Wwc2-specific siRNA (10μM), 2in2 micro-injection, 16-cell (E3.0) stage | | | | | |
|---|---|---|---|---|---|
| # | Total cell number per embryo | Total number of dividing cells | Number of cells exhibiting an atypical/ unusual phenotype morphology | | |
| | | | Abnormal nuclear morphology (e.g. non-spheroidal or mid-body associated) | Defective cytokinesis (i.e. >x2 equally sized nuclei per cell) | Presence of micronuclei |
| **1** | 8 | | 0 | 0 | 0 |
| **2** | 9 | | 1 | 0 | 0 |
| **3** | 8 | | 1 | 0 | 0 |
| **4** | 8 | | 0 | 1 | 0 |
| **5** | 8 | | 0 | 0 | 3 |
| **6** | 8 | | 0 | 0 | 0 |
| **7** | 8 | | 0 | 0 | 1 |
| **8** | 10 | | 0 | 0 | 0 |
| **9** | 9 | | 2 | 0 | 1 |
| **10** | 8 | | 0 | 0 | 0 |
| **11** | 9 | | 1 | 1 | 0 |
| **12** | 8 | | 1 | 0 | 0 |
| **13** | 8 | | 1 | 0 | 1 |
| **14** | 10 | | 0 | 0 | 0 |
| **15** | 14 | | 0 | 0 | 0 |
| **16** | 8 | | 0 | 0 | 0 |
| **17** | 8 | | 0 | 1 | 1 |
| **18** | 8 | | 2 | 0 | 1 |
| **19** | 8 | | 2 | 0 | 0 |
| **20** | 12 | | 2 | 1 | 2 |
| **21** | 6 | | 1 | 0 | 0 |
| **22** | 10 | | 1 | 1 | 1 |
| **23** | 5 | | 1 | 0 | 0 |
| **24** | 5 | | 0 | 0 | 2 |
| **25** | 10 | | 1 | 0 | 2 |
| **26** | 7 | | 0 | 0 | 1 |
| **27** | 14 | | 1 | 1 | 3 |
| **28** | 10 | | 2 | 1 | 2 |
| **29** | 8 | | 1 | 0 | 0 |
| **30** | 8 | | 1 | 0 | 0 |
| **31** | 8 | | 0 | 0 | 0 |
| **TOTAL** | **268** | **0** | **22** | **7** | **21** |
| **AVERAGE** | **8.6** | **0.0** | **0.7** | **0.2** | **0.7** |
| SEM | **0.4** | **0.0** | **0.1** | **0.1** | **0.2** |
| Stat. sig. (exp. vs. con embryo) *p<0.05, **p<0.005 | ** | | ** | * | ** |
| p-value (2-tailed students t-test) | 3.00E-17 | 0.00E+00 | 2.11E-06 | 5.10E-03 | 2.28E-04 |

Supplementary Tables ST6

| Control siRNA (10μM), 2in2 micro-injection, 32-cell (E3.5) stage | | | | | |
|---|---|---|---|---|---|
| # | Total cell number per embryo | Total number of dividing cells | Number of cells exhibiting an atypical/ unusual phenotype morphology | | |
| | | | Abnormal nuclear morphology (e.g. non-spheroidal or mid-body associated) | Defective cytokinesis (i.e. >x2 equally sized nuclei per cell) | Presence of micronuclei |
| **1** | 33 | | 0 | 0 | 0 |
| **2** | 24 | | 0 | 0 | 0 |
| **3** | 28 | | 0 | 0 | 0 |
| **4** | 31 | | 0 | 0 | 0 |
| **5** | 31 | | 0 | 0 | 0 |
| **6** | 38 | | 0 | 0 | 0 |
| **7** | 32 | | 0 | 0 | 0 |
| **8** | 31 | | 0 | 0 | 0 |
| **9** | 35 | | 0 | 0 | 0 |
| **10** | 31 | | 0 | 0 | 0 |
| **11** | 31 | | 0 | 0 | 0 |
| **12** | 34 | | 0 | 0 | 0 |
| **13** | 32 | | 0 | 0 | 0 |
| **14** | 22 | | 0 | 0 | 0 |
| **15** | 24 | | 0 | 0 | 0 |
| **16** | 35 | | 0 | 0 | 0 |
| **17** | 32 | | 0 | 0 | 0 |
| **18** | 38 | | 0 | 0 | 0 |
| **19** | 36 | | 0 | 0 | 0 |
| **20** | 37 | | 0 | 0 | 0 |
| **21** | 32 | | 0 | 0 | 0 |
| **22** | 42 | | 0 | 0 | 0 |
| **23** | 32 | | 0 | 0 | 0 |
| **24** | 29 | | 0 | 0 | 0 |
| **25** | 32 | | 0 | 0 | 0 |
| **26** | 32 | | 0 | 0 | 0 |
| **27** | 32 | | 0 | 0 | 0 |
| **28** | 31 | | 0 | 0 | 1 |
| **29** | 26 | | 0 | 0 | 0 |
| **30** | 28 | | 0 | 0 | 0 |
| **TOTAL** | **951** | **0** | **0** | **0** | **1** |
| **AVERAGE** | **31.7** | **0.0** | **0.0** | **0.0** | **0.0** |
| SEM | **0.8** | **0.0** | **0.0** | **0.0** | **0.03** |

| Wwc2-specific siRNA (10μM), 2in2 micro-injection, 32-cell (E3.5) stage | | | | | |
|---|---|---|---|---|---|
| # | Total cell number per embryo | Total number of dividing cells | Number of cells exhibiting an atypical/ unusual phenotype morphology | | |
| | | | Abnormal nuclear morphology (e.g. non-spheroidal or mid-body associated) | Defective cytokinesis (i.e. >x2 equally sized nuclei per cell) | Presence of micronuclei |
| **1** | 7 | | 0 | 0 | 3 |
| **2** | 15 | | 0 | 0 | 0 |
| **3** | 7 | | 0 | 0 | 3 |
| **4** | 14 | | 0 | 0 | 0 |
| **5** | 6 | | 1 | 0 | 1 |
| **6** | 7 | | 1 | 0 | 1 |
| **7** | 8 | | 1 | 1 | 0 |
| **8** | 7 | | 0 | 0 | 0 |
| **9** | 7 | | 0 | 0 | 1 |
| **10** | 8 | | 0 | 0 | 1 |
| **11** | 8 | | 2 | 0 | 3 |
| **12** | 12 | | 0 | 0 | 0 |
| **13** | 12 | | 0 | 0 | 1 |
| **14** | 7 | | 1 | 0 | 0 |
| **15** | 9 | | 0 | 0 | 4 |
| **16** | 10 | | 1 | 0 | 2 |
| **17** | 7 | | 0 | 0 | 2 |
| **18** | 7 | | 0 | 0 | 0 |
| **19** | 16 | | 0 | 0 | 0 |
| **20** | 9 | | 0 | 1 | 2 |
| **21** | 12 | | 2 | 1 | 3 |
| **22** | 16 | | 1 | 2 | 6 |
| **23** | 13 | | 4 | 1 | 4 |
| **24** | 6 | | 2 | 1 | 0 |
| **25** | 17 | | 1 | 0 | 0 |
| **26** | 9 | | 1 | 0 | 2 |
| **27** | 8 | | 0 | 0 | 0 |
| **28** | 8 | | 0 | 1 | 4 |
| **29** | 8 | | 2 | 0 | 0 |
| **30** | 8 | | 1 | 0 | 0 |
| **31** | 11 | | 0 | 1 | 0 |
| **32** | 6 | | 0 | 0 | 2 |
| **33** | 12 | | 1 | 0 | 1 |
| **34** | 7 | | 0 | 0 | 1 |
| **35** | 15 | | 0 | 0 | 2 |
| **36** | 8 | | 0 | 0 | 0 |
| **TOTAL** | **347** | **0** | **22** | **9** | **49** |
| **AVERAGE** | **9.6** | **0.0** | **0.6** | **0.3** | **1.4** |
| SEM | **0.5** | **0** | **0.2** | **0.1** | **0.3** |
| Stat. sig. (exp. vs. con embryo) *p<0.05, **p<0.005 | ** | | ** | * | ** |
| p-value (2-tailed students t-test) | 2.88E-33 | 0.00E+00 | 4.49E-04 | 8.06E-03 | 1.69E-05 |

**Supplementary Tables ST7**

| Control siRNA (10µM), 2in2 micro-injection, ~64-cell (E4.0) stage | | | | | |
|---|---|---|---|---|---|
| # | Total cell number per embryo | Total number of dividing cells | Number of cells exhibiting an atypical/ unusual phenotype morphology | | |
| | | | Abnormal nuclear morphology (e.g. non-spheroidal or mid-body associated) | Defective cytokinesis (i.e. >x2 equally sized nuclei per cell) | Presence of micronuclei |
| **1** | 60 | 3 | 0 | 0 | 0 |
| **2** | 68 | 4 | 0 | 0 | 0 |
| **3** | 68 | 8 | 0 | 0 | 0 |
| **4** | 72 | 3 | 0 | 0 | 0 |
| **5** | 64 | 1 | 0 | 0 | 0 |
| **6** | 70 | 0 | 0 | 0 | 0 |
| **7** | 62 | 5 | 0 | 0 | 0 |
| **8** | 68 | 4 | 0 | 0 | 0 |
| **9** | 60 | 3 | 0 | 0 | 0 |
| **10** | 66 | 5 | 0 | 0 | 0 |
| **11** | 72 | 2 | 0 | 0 | 0 |
| **12** | 64 | 1 | 0 | 0 | 0 |
| **13** | 68 | 3 | 0 | 0 | 0 |
| **14** | 60 | 2 | 0 | 0 | 0 |
| **15** | 70 | 1 | 0 | 0 | 0 |
| **16** | 66 | 3 | 0 | 0 | 0 |
| **TOTAL** | **1058** | **48** | **0** | **0** | **0** |
| **AVERAGE** | **66.1** | **3.0** | **0.0** | **0.0** | **0.0** |
| SEM | **1.0** | **0.5** | **0.0** | **0.0** | **0.0** |

| Wwc2-specific siRNA (10µM), 2in2 micro-injection, ~64-cell (E4.0) stage | | | | | |
|---|---|---|---|---|---|
| # | Total cell number per embryo | Total number of dividing cells | Number of cells exhibiting an atypical/ unusual phenotype morphology | | |
| | | | Abnormal nuclear morphology (e.g. non-spheroidal or mid-body associated) | Defective cytokinesis (i.e. >x2 equally sized nuclei per cell) | Presence of micronuclei |
| **1** | 34 | 0 | 2 | 0 | 7 |
| **2** | 24 | 1 | 3 | 0 | 9 |
| **3** | 22 | 2 | 1 | 0 | 4 |
| **4** | 24 | 2 | 2 | 0 | 3 |
| **5** | 18 | 0 | 3 | 0 | 2 |
| **6** | 21 | 0 | 4 | 0 | 3 |
| **7** | 20 | 0 | 3 | 0 | 3 |
| **8** | 29 | 1 | 2 | 0 | 8 |
| **9** | 16 | 0 | 4 | 0 | 4 |
| **10** | 18 | 0 | 2 | 0 | 4 |
| **11** | 23 | 0 | 1 | 0 | 5 |
| **12** | 16 | 0 | 4 | 0 | 8 |
| **13** | 18 | 3 | 2 | 0 | 4 |
| **14** | 22 | 2 | 3 | 0 | 6 |
| **15** | 16 | 0 | 4 | 2 | 3 |
| **16** | 23 | 1 | 2 | 0 | 3 |
| **17** | 20 | 0 | 4 | 0 | 7 |
| **18** | 24 | 0 | 3 | 0 | 5 |
| **TOTAL** | **388** | **12** | **49** | **2** | **88** |
| **AVERAGE** | **21.6** | **0.7** | **2.7** | **0.1** | **4.9** |
| SEM | **1.1** | **0.2** | **0.2** | **0.1** | **0.5** |
| Stat. sig. (exp. vs. con embryo) *p<0.05, **p<0.005 | ** | ** | ** | | ** |
| p-value (2-tailed students t-test) | 9.26E-25 | 9.32E-05 | 4.43E-12 | 3.54E-01 | 1.49E-10 |

Supplementary Tables ST8

| Control siRNA (10µM), 2in2 micro-injection, >64-cell (E4.5) stage | | | | | |
|---|---|---|---|---|---|
| # | Total cell number per embryo | Total number of dividing cells | Number of cells exhibiting an atypical/ unusual phenotype morphology | | |
| | | | Abnormal nuclear morphology (e.g. non-spheroidal or mid-body associated) | Defective cytokinesis (i.e. >x2 equally sized nuclei per cell) | Presence of micronuclei |
| **1** | 81 | 4 | 0 | 0 | 0 |
| **2** | 76 | 2 | 0 | 0 | 0 |
| **3** | 67 | 8 | 0 | 0 | 0 |
| **4** | 77 | 4 | 0 | 0 | 0 |
| **5** | 110 | 5 | 0 | 0 | 0 |
| **6** | 107 | 6 | 0 | 0 | 0 |
| **7** | 76 | 5 | 0 | 0 | 0 |
| **8** | 94 | 4 | 0 | 0 | 0 |
| **9** | 102 | 1 | 0 | 0 | 0 |
| **10** | 108 | 7 | 0 | 0 | 0 |
| **11** | 85 | 5 | 0 | 0 | 0 |
| **12** | 73 | 4 | 0 | 0 | 0 |
| **13** | 82 | 7 | 0 | 0 | 0 |
| **14** | 80 | 5 | 0 | 0 | 0 |
| **TOTAL** | **1218** | **67** | **0** | **0** | **0** |
| **AVERAGE** | **87.0** | **4.8** | **0.0** | **0.0** | **0.0** |
| SEM | **3.8** | **0.5** | **0.0** | **0.0** | **0.0** |

| Wwc2-specific siRNA (10µM), 2in2 micro-injection, >64-cell (E4.5) stage | | | | | |
|---|---|---|---|---|---|
| # | Total cell number per embryo | Total number of dividing cells | Number of cells exhibiting an atypical/ unusual phenotype morphology | | |
| | | | Abnormal nuclear morphology (e.g. non-spheroidal or mid-body associated) | Defective cytokinesis (i.e. >x2 equally sized nuclei per cell) | Presence of micronuclei |
| **1** | 13 | 0 | 6 | 1 | 6 |
| **2** | 57 | 7 | 7 | 1 | 5 |
| **3** | 31 | 3 | 5 | 0 | 7 |
| **4** | 13 | 1 | 2 | 0 | 2 |
| **5** | 21 | 6 | 4 | 0 | 5 |
| **6** | 28 | 2 | 4 | 0 | 4 |
| **7** | 30 | 3 | 5 | 0 | 3 |
| **8** | 28 | 2 | 4 | 0 | 3 |
| **9** | 36 | 1 | 3 | 1 | 6 |
| **10** | 26 | 5 | 2 | 0 | 2 |
| **11** | 51 | 5 | 4 | 0 | 4 |
| **12** | 35 | 3 | 2 | 0 | 7 |
| **13** | 45 | 3 | 5 | 0 | 3 |
| **14** | 32 | 3 | 3 | 0 | 4 |
| **15** | 44 | 4 | 4 | 0 | 4 |
| **16** | 19 | 3 | 3 | 0 | 2 |
| **17** | 23 | 5 | 2 | 1 | 3 |
| **TOTAL** | **532** | **56** | **65** | **4** | **70** |
| **AVERAGE** | **31.3** | **3.3** | **3.8** | **0.2** | **4.1** |
| SEM | **3.0** | **0.5** | **0.4** | **0.1** | **0.4** |
| Stat. sig. (exp. vs. con embryo) *p<0.05, **p<0.005 | ** | * | ** | | ** |
| p-value (2-tailed students t-test) | 2.27E-12 | 3.55E-02 | 1.26E-10 | 5.41E-02 | 3.44E-10 |

**Supplementary Tables ST9**

| Control siRNA (10μM), 1in2 micro-injection, 4-cell (E2.0) stage | | | | | | |
|---|---|---|---|---|---|---|
| # | Total cell number per embryo | Cells in injected clone | Cells in non-injected clone | Dividing cells in injected clone | Dividing cells in non-injected clone | Total number of dividing cells |
| **1** | 4 | 2 | 2 | 0 | 0 | 0 |
| **2** | 4 | 2 | 2 | 0 | 0 | 0 |
| **3** | 6 | 3 | 3 | 0 | 1 | 1 |
| **4** | 5 | 3 | 2 | 0 | 1 | 1 |
| **5** | 4 | 2 | 2 | 0 | 0 | 0 |
| **6** | 4 | 2 | 2 | 1 | 1 | 2 |
| **7** | 5 | 2 | 3 | 1 | 0 | 1 |
| **8** | 4 | 2 | 2 | 1 | 1 | 2 |
| **9** | 6 | 3 | 3 | 1 | 1 | 2 |
| **10** | 4 | 2 | 2 | 0 | 0 | 0 |
| **11** | 5 | 2 | 3 | 1 | 0 | 1 |
| **12** | 4 | 2 | 2 | 0 | 0 | 0 |
| **TOTAL** | **55** | **27** | **28** | **5** | **5** | **10** |
| **AVERAGE** | **4.6** | **2.3** | **2.3** | **0.4** | **0.4** | **0.8** |
| SEM | **0.2** | **0.1** | **0.1** | **0.1** | **0.1** | **0.2** |
| Stat. sig. (inter-clone) ‡p<0.05, ‡‡p<0.005 | | | | | | |
| p-value (2-tailed students t-test) | | 6.70E-01 | | 1.00E+00 | | |

| Wwc2-specific siRNA (10μM), 1in2 micro-injection, 4-cell (E2.0) stage | | | | | | |
|---|---|---|---|---|---|---|
| # | Total cell number per embryo | Cells in injected clone | Cells in non-injected clone | Dividing cells in injected clone | Dividing cells in non-injected clone | Total number of dividing cells |
| **1** | 4 | 2 | 2 | 0 | 1 | 1 |
| **2** | 4 | 2 | 2 | 0 | 0 | 0 |
| **3** | 5 | 3 | 2 | 0 | 1 | 1 |
| **4** | 4 | 2 | 2 | 0 | 0 | 0 |
| **5** | 4 | 2 | 2 | 0 | 1 | 1 |
| **6** | 4 | 2 | 2 | 0 | 0 | 0 |
| **7** | 5 | 2 | 3 | 0 | 1 | 1 |
| **8** | 4 | 2 | 2 | 0 | 1 | 1 |
| **9** | 4 | 2 | 2 | 0 | 0 | 0 |
| **10** | 4 | 2 | 2 | 0 | 0 | 0 |
| **11** | 5 | 2 | 3 | 0 | 0 | 0 |
| **12** | 4 | 2 | 2 | 0 | 0 | 0 |
| **13** | 4 | 2 | 2 | 0 | 1 | 1 |
| **14** | 5 | 2 | 3 | 0 | 0 | 0 |
| **15** | 4 | 2 | 2 | 0 | 1 | 1 |
| **TOTAL** | **64** | **31** | **33** | **0** | **7** | **7** |
| **AVERAGE** | **4.3** | **2.1** | **2.2** | **0.0** | **0.5** | **0.5** |
| SEM | **0.1** | **0.1** | **0.1** | **0.0** | **0.1** | **0.1** |
| Stat. sig. (inter-clone) ‡p<0.05, ‡‡p<0.005 | | | | **‡‡** | | |
| p-value (2-tailed students t-test) | | 2.99E-01 | | 1.58E-03 | | |
| Stat. sig. (exp. vs. con embryo) *p<0.05, **p<0.005 | | | | ** | | |
| p-value (2-tailed students t-test) | 2.05E-01 | 1.97E-01 | 4.52E-01 | 4.20E-03 | 8.04E-01 | 1.73E-01 |

**Supplementary Tables ST10**

| Control siRNA (10μM), 1in2 micro-injection, 8-cell (E2.5) stage | | | | | | |
|---|---|---|---|---|---|---|
| # | Total cell number per embryo | Cells in injected clone | Cells in non-injected clone | Dividing cells in injected clone | Dividing cells in non-injected clone | Total number of dividing cells |
| **1** | 8 | 4 | 4 | 0 | 0 | 0 |
| **2** | 8 | 4 | 4 | 0 | 1 | 1 |
| **3** | 8 | 4 | 4 | 0 | 1 | 1 |
| **4** | 9 | 4 | 5 | 0 | 0 | 0 |
| **5** | 8 | 4 | 4 | 0 | 0 | 0 |
| **6** | 8 | 4 | 4 | 1 | 0 | 1 |
| **7** | 8 | 4 | 4 | 0 | 0 | 0 |
| **8** | 7 | 4 | 4 | 0 | 0 | 0 |
| **9** | 8 | 4 | 4 | 0 | 1 | 1 |
| **10** | 9 | 4 | 5 | 1 | 0 | 1 |
| **11** | 8 | 4 | 4 | 0 | 1 | 1 |
| **12** | 9 | 4 | 5 | 0 | 0 | 0 |
| **13** | 8 | 4 | 4 | 1 | 0 | 1 |
| **TOTAL** | **106** | **52** | **55** | **3** | **4** | **7** |
| **AVERAGE** | **8.2** | **4.0** | **4.2** | **0.2** | **0.3** | **0.5** |
| SEM | **0.2** | **0.0** | **0.1** | **0.1** | **0.1** | **0.1** |
| Stat. sig. (inter-clone) ‡p<0.05, ‡‡p<0.005 | | | | | | |
| p-value (2-tailed students t-test) | | 6.99E-02 | | 6.74E-01 | | |

| Wwc2-specific siRNA (10μM), 1in2 micro-injection, 8-cell (E2.5) stage | | | | | | |
|---|---|---|---|---|---|---|
| # | Total cell number per embryo | Cells in injected clone | Cells in non-injected clone | Dividing cells in injected clone | Dividing cells in non-injected clone | Total number of dividing cells |
| **1** | 6 | 2 | 4 | 0 | 2 | 2 |
| **2** | 5 | 2 | 3 | 0 | 1 | 1 |
| **3** | 6 | 2 | 4 | 0 | 0 | 0 |
| **4** | 6 | 2 | 4 | 0 | 0 | 0 |
| **5** | 7 | 3 | 4 | 0 | 0 | 0 |
| **6** | 7 | 3 | 4 | 0 | 1 | 1 |
| **7** | 6 | 2 | 4 | 1 | 1 | 2 |
| **8** | 6 | 2 | 4 | 0 | 1 | 1 |
| **9** | 6 | 2 | 4 | 0 | 0 | 0 |
| **10** | 8 | 4 | 4 | 0 | 1 | 1 |
| **11** | 6 | 2 | 4 | 0 | 0 | 0 |
| **12** | 7 | 2 | 5 | 0 | 0 | 0 |
| **13** | 5 | 2 | 3 | 0 | 1 | 1 |
| **14** | 6 | 2 | 4 | 0 | 0 | 0 |
| **15** | 6 | 2 | 4 | 0 | 0 | 0 |
| **16** | 6 | 2 | 4 | 1 | 2 | 3 |
| **17** | 6 | 2 | 4 | 0 | 0 | 0 |
| **18** | 6 | 2 | 4 | 0 | 0 | 0 |
| **19** | 6 | 2 | 4 | 0 | 1 | 1 |
| **TOTAL** | **117** | **42** | **75** | **2** | **11** | **13** |
| **AVERAGE** | **6.2** | **2.2** | **3.9** | **0.1** | **0.6** | **0.7** |
| SEM | **0.2** | **0.1** | **0.1** | **0.1** | **0.2** | **0.6** |
| Stat. sig. (inter-clone) ‡p<0.05, ‡‡p<0.005 | | **‡‡** | | **‡** | | |
| p-value (2-tailed t-test) | | 2.25E-13 | | 1.01E-02 | | |
| Stat. sig. (exp. vs. con embryo) *p<0.05, **p<0.005 | ** | ** | | | | |
| p-value (2-tailed students t-test) | 1.08E-09 | 5.70E-13 | 6.97E-02 | 3.53E-01 | 2.31E-01 | 5.98E-01 |

**Supplementary Tables ST11**

| Control siRNA (10μM), 1in2 micro-injection, 16-cell (E3.0) stage | | | | | | |
|---|---|---|---|---|---|---|
| # | Total cell number per embryo | Cells in injected clone | Cells in non-injected clone | Dividing cells in injected clone | Dividing cells in non-injected clone | Total number of dividing cells |
| **1** | 16 | 8 | 8 | 0 | 0 | 0 |
| **2** | 16 | 8 | 8 | 0 | 0 | 0 |
| **3** | 16 | 8 | 8 | 0 | 0 | 0 |
| **4** | 15 | 7 | 8 | 0 | 2 | 2 |
| **5** | 17 | 8 | 9 | 1 | 0 | 1 |
| **6** | 16 | 8 | 8 | 0 | 3 | 3 |
| **7** | 16 | 8 | 8 | 0 | 0 | 0 |
| **8** | 15 | 8 | 7 | 0 | 0 | 0 |
| **9** | 15 | 7 | 8 | 0 | 0 | 0 |
| **10** | 16 | 8 | 8 | 0 | 0 | 0 |
| **11** | 15 | 7 | 8 | 0 | 0 | 0 |
| **12** | 16 | 8 | 8 | 1 | 0 | 1 |
| **13** | 15 | 8 | 7 | 1 | 1 | 2 |
| **TOTAL** | **204** | **101** | **103** | **3** | **6** | **9** |
| **AVERAGE** | **15.7** | **7.8** | **7.9** | **0.2** | **0.5** | **0.7** |
| SEM | **0.2** | **0.1** | **0.1** | **0.088** | **0.300** | **0.3** |
| Stat. sig. (inter-clone) ‡p<0.05, ‡‡p<0.005 | | | | | | |
| p-value (2-tailed students t-test) | | 4.09E-01 | | 4.41E-01 | | |

| Wwc2-specific siRNA (10μM), 1in2 micro-injection, 16-cell (E3.0) stage | | | | | | |
|---|---|---|---|---|---|---|
| # | Total cell number per embryo | Cells in injected clone | Cells in non-injected clone | Dividing cells in injected clone | Dividing cells in non-injected clone | Total number of dividing cells |
| **1** | 11 | 4 | 7 | 0 | 1 | 1 |
| **2** | 13 | 5 | 8 | 1 | 0 | 1 |
| **3** | 12 | 4 | 8 | 0 | 0 | 0 |
| **4** | 12 | 4 | 8 | 0 | 0 | 0 |
| **5** | 13 | 5 | 8 | 0 | 1 | 1 |
| **6** | 12 | 3 | 9 | 0 | 1 | 1 |
| **7** | 12 | 4 | 8 | 0 | 0 | 0 |
| **8** | 12 | 4 | 8 | 0 | 0 | 0 |
| **9** | 12 | 4 | 8 | 1 | 0 | 1 |
| **10** | 11 | 4 | 7 | 0 | 1 | 1 |
| **11** | 12 | 4 | 8 | 0 | 1 | 1 |
| **12** | 12 | 4 | 8 | 0 | 0 | 0 |
| **13** | 11 | 4 | 7 | 0 | 0 | 0 |
| **14** | 14 | 6 | 8 | 1 | 0 | 1 |
| **15** | 12 | 4 | 8 | 0 | 0 | 0 |
| **16** | 14 | 4 | 10 | 0 | 1 | 1 |
| **17** | 11 | 3 | 8 | 0 | 0 | 0 |
| **18** | 12 | 4 | 8 | 1 | 1 | 2 |
| **TOTAL** | **218** | **74** | **144** | **4** | **7** | **11** |
| **AVERAGE** | **12.1** | **4.1** | **8.0** | **0.2** | **0.4** | **0.6** |
| SEM | **0.2** | **0.2** | **0.2** | **0.101** | **0.118** | **0.1** |
| Stat. sig. (inter-clone) ‡p<0.05, ‡‡p<0.005 | | ‡‡ | | | | |
| p-value (2-tailed students t-test) | | 2.73E-18 | | 2.91E-01 | | |
| Stat. sig. (exp. vs. con embryo) *p<0.05, **p<0.005 | ** | ** | | | | |
| p-value (2-tailed students t-test) | 4.93E-13 | 1.19E-16 | 7.33E-01 | 9.57E-01 | 7.87E-01 | 7.85E-01 |

**Supplementary Tables ST12**

| Control siRNA (10μM), 1in2 micro-injection, 32-cell (E3.5) stage | | | | | | |
|---|---|---|---|---|---|---|
| # | Total cell number per embryo | Cells in injected clone | Cells in non-injected clone | Dividing cells in injected clone | Dividing cells in non-injected clone | Total number of dividing cells |
| **1** | 43 | 22 | 21 | 3 | 6 | 9 |
| **2** | 31 | 16 | 15 | 0 | 1 | 1 |
| **3** | 33 | 16 | 17 | 0 | 1 | 1 |
| **4** | 35 | 17 | 18 | 6 | 3 | 9 |
| **5** | 35 | 16 | 19 | 3 | 1 | 4 |
| **6** | 32 | 16 | 16 | 0 | 0 | 0 |
| **7** | 31 | 15 | 16 | 1 | 0 | 1 |
| **8** | 33 | 16 | 17 | 0 | 0 | 0 |
| **9** | 31 | 16 | 15 | 0 | 1 | 1 |
| **10** | 34 | 16 | 18 | 0 | 0 | 0 |
| **TOTAL** | **338** | **166** | **172** | **13** | **13** | **26** |
| **AVERAGE** | **33.8** | **16.6** | **17.2** | **1.3** | **1.3** | **2.6** |
| SEM | **1.1** | **0.6** | **0.6** | **0.7** | **0.6** | **1.1** |
| Stat. sig. (inter-clone) ‡p<0.05, ‡‡p<0.005 | | | | | | |
| p-value (2-tailed students t-test) | | 4.92E-01 | | 1.00E+00 | | |

| Wwc2-specific siRNA (10μM), 1in2 micro-injection, 32-cell (E3.5) stage | | | | | | |
|---|---|---|---|---|---|---|
| # | Total cell number per embryo | Cells in injected clone | Cells in non-injected clone | Dividing cells in injected clone | Dividing cells in non-injected clone | Total number of dividing cells |
| **1** | 12 | 3 | 9 | 0 | 1 | 1 |
| **2** | 25 | 10 | 15 | 0 | 1 | 1 |
| **3** | 18 | 3 | 15 | 0 | 0 | 0 |
| **4** | 21 | 6 | 15 | 0 | 2 | 2 |
| **5** | 20 | 4 | 16 | 1 | 0 | 1 |
| **6** | 21 | 5 | 16 | 1 | 0 | 1 |
| **7** | 21 | 7 | 14 | 1 | 0 | 1 |
| **8** | 14 | 4 | 10 | 0 | 2 | 2 |
| **9** | 23 | 8 | 15 | 0 | 0 | 0 |
| **10** | 26 | 12 | 14 | 1 | 0 | 1 |
| **11** | 37 | 16 | 21 | 0 | 5 | 5 |
| **12** | 20 | 6 | 14 | 2 | 1 | 3 |
| **13** | 32 | 16 | 16 | 0 | 0 | 0 |
| **14** | 19 | 5 | 14 | 0 | 0 | 0 |
| **15** | 23 | 8 | 15 | 0 | 0 | 0 |
| **16** | 24 | 8 | 16 | 0 | 0 | 0 |
| **17** | 19 | 4 | 15 | 0 | 0 | 0 |
| **18** | 20 | 4 | 16 | 0 | 0 | 0 |
| **19** | 26 | 10 | 16 | 0 | 0 | 0 |
| **20** | 17 | 4 | 13 | 0 | 0 | 0 |
| **21** | 20 | 4 | 16 | 0 | 0 | 0 |
| **22** | 17 | 4 | 13 | 0 | 2 | 2 |
| **23** | 20 | 4 | 16 | 0 | 0 | 0 |
| **24** | 24 | 8 | 16 | 0 | 0 | 0 |
| **25** | 25 | 9 | 16 | 0 | 0 | 0 |
| **26** | 24 | 8 | 16 | 0 | 0 | 0 |
| **27** | 18 | 6 | 12 | 0 | 2 | 2 |
| **28** | 22 | 6 | 16 | 2 | 0 | 2 |
| **29** | 22 | 6 | 16 | 1 | 0 | 1 |
| **30** | 23 | 8 | 15 | 0 | 0 | 0 |
| **TOTAL** | **653** | **206** | **447** | **9** | **16** | **25** |
| **AVERAGE** | **21.8** | **6.9** | **14.9** | **0.3** | **0.5** | **0.8** |
| SEM | **0.9** | **0.6** | **0.4** | **0.1** | **0.2** | **0.2** |
| Stat. sig. (inter-clone) ‡p<0.05, ‡‡p<0.005 | | ‡‡ | | | | |
| p-value (2-tailed students t-test) | | 7.46E-16 | | 3.13E-01 | | |
| Stat. sig. (exp. vs. con embryo) *p<0.05, **p<0.005 | ** | ** | ** | * | | * |
| p-value (2-tailed students t-test) | 1.23E-08 | 1.95E-10 | 4.18E-03 | 2.01E-02 | 1.24E-01 | 2.15E-02 |

**Supplementary Tables ST13**

| Control siRNA (10µM), 1in2 micro-injection, >64-cell (E4.5) stage | | | | | | |
|---|---|---|---|---|---|---|
| # | Total cell number per embryo | Cells in injected clone | Cells in non-injected clone | Dividing cells in injected clone | Dividing cells in non-injected clone | Total number of dividing cells |
| **1** | 75 | 36 | 39 | 1 | 1 | 2 |
| **2** | 69 | 33 | 36 | 2 | 1 | 3 |
| **3** | 62 | 29 | 33 | 1 | 1 | 2 |
| **4** | 68 | 35 | 33 | 0 | 1 | 1 |
| **5** | 65 | 32 | 33 | 1 | 0 | 1 |
| **6** | 86 | 42 | 44 | 2 | 0 | 2 |
| **7** | 67 | 33 | 34 | 2 | 0 | 2 |
| **8** | 76 | 37 | 39 | 1 | 0 | 1 |
| **9** | 68 | 33 | 35 | 1 | 1 | 2 |
| **10** | 77 | 38 | 39 | 3 | 0 | 3 |
| **11** | 80 | 36 | 44 | 1 | 0 | 1 |
| **12** | 68 | 34 | 34 | 2 | 0 | 2 |
| **13** | 80 | 39 | 41 | 0 | 0 | 0 |
| **TOTAL** | **941** | **457** | **484** | **17** | **5** | **22** |
| **AVERAGE** | **72.4** | **35.2** | **37.2** | **1.3** | **0.4** | **1.7** |
| SEM | **2.0** | **0.9** | **1.1** | **0.2** | **0.1** | **0.2** |
| Stat. sig. (inter-clone) ‡p<0.05, ‡‡p<0.005 | | | | ‡‡ | | |
| p-value (2-tailed students t-test) | | 1.69E-01 | | 2.67E-03 | | |

| Wwc2-specific siRNA (10µM), 1in2 micro-injection, >64-cell (E4.5) stage | | | | | | |
|---|---|---|---|---|---|---|
| # | Total cell number per embryo | Cells in injected clone | Cells in non-injected clone | Dividing cells in injected clone | Dividing cells in non-injected clone | Total number of dividing cells |
| **1** | 45 | 15 | 30 | 0 | 5 | 5 |
| **2** | 33 | 9 | 24 | 0 | 3 | 3 |
| **3** | 44 | 14 | 30 | 0 | 1 | 1 |
| **4** | 37 | 8 | 29 | 0 | 2 | 2 |
| **5** | 62 | 15 | 47 | 2 | 0 | 2 |
| **6** | 45 | 14 | 31 | 0 | 3 | 3 |
| **7** | 44 | 18 | 26 | 0 | 1 | 1 |
| **8** | 61 | 19 | 42 | 0 | 3 | 3 |
| **9** | 69 | 19 | 50 | 1 | 4 | 5 |
| **10** | 53 | 11 | 42 | 1 | 2 | 3 |
| **11** | 59 | 21 | 38 | 1 | 2 | 3 |
| **12** | 52 | 16 | 36 | 0 | 1 | 1 |
| **13** | 43 | 16 | 27 | 1 | 5 | 6 |
| **14** | 57 | 17 | 40 | 1 | 1 | 2 |
| **15** | 54 | 14 | 40 | 0 | 2 | 2 |
| **16** | 54 | 17 | 37 | 0 | 4 | 4 |
| **TOTAL** | **812** | **243** | **569** | **7** | **39** | **46** |
| **AVERAGE** | **50.8** | **15.2** | **35.6** | **0.4** | **2.4** | **2.9** |
| SEM | **2.4** | **0.9** | **1.9** | **0.2** | **0.4** | **0.4** |
| Stat. sig. (inter-clone) ‡p<0.05, ‡‡p<0.005 | | ‡‡ | | ‡‡ | | |
| p-value (2-tailed students t-test) | | 1.21E-10 | | 3.03E-05 | | |
| Stat. sig. (exp. vs. con embryo) *p<0.05, **p<0.005 | ** | ** | | ** | ** | * |
| p-value (2-tailed students t-test) | 3.49E-07 | 7.49E-15 | 4.88E-01 | 3.89E-03 | 6.90E-05 | 1.78E-02 |

**Supplementary Tables ST14**

| IVM culturing control, non-micro-injected GV stage (+18h IBMX & 16h post IBMX wash-out) | | | | | | |
|---|---|---|---|---|---|---|
| # | Meiotic spindle status | | | | 1st meiotic polar body (PB1) | |
| | Normal MII arrested (+PB1) | Normal MI stage (no PB1) | Defective (e.g. dispersed chromosomes, no PB1) | Absent with ultra condensed chromatin | Present | Absent |
| 1 | 0 | 1 | 0 | 0 | 0 | 1 |
| 2 | 1 | 0 | 0 | 0 | 1 | 0 |
| 3 | 1 | 0 | 0 | 0 | 1 | 0 |
| 4 | 1 | 0 | 0 | 0 | 1 | 0 |
| 5 | 1 | 0 | 0 | 0 | 1 | 0 |
| 6 | 1 | 0 | 0 | 0 | 1 | 0 |
| 7 | 1 | 0 | 0 | 0 | 1 | 0 |
| 8 | 1 | 0 | 0 | 0 | 1 | 0 |
| 9 | 1 | 0 | 0 | 0 | 1 | 0 |
| 10 | 1 | 0 | 0 | 0 | 1 | 0 |
| 11 | 1 | 0 | 0 | 0 | 1 | 0 |
| 12 | 1 | 0 | 0 | 0 | 1 | 0 |
| 13 | 1 | 0 | 0 | 0 | 1 | 0 |
| 14 | 1 | 0 | 0 | 0 | 1 | 0 |
| 15 | 1 | 0 | 0 | 0 | 1 | 0 |
| 16 | 1 | 0 | 0 | 0 | 1 | 0 |
| 17 | 1 | 0 | 0 | 0 | 1 | 0 |
| 18 | 1 | 0 | 0 | 0 | 1 | 0 |
| TOTAL | 17 | 1 | 0 | 0 | 17 | 1 |
| PERCENTAGE | 94.4 | 5.6 | 0.0 | 0.0 | 94.4 | 5.6 |

| Control siRNA microinjected GV stage (+18h IBMX & 16h post IBMX wash-out) | | | | | | |
|---|---|---|---|---|---|---|
| # | Meiotic spindle status | | | | 1st meiotic polar body (PB1) | |
| | Normal MII arrested (+PB1) | Normal MI stage (no PB1) | Defective (e.g. dispersed chromosomes, no PB1) | Absent with ultra condensed chromatin | Present | Absent |
| 1 | 1 | 0 | 0 | 0 | 1 | 0 |
| 2 | 1 | 0 | 0 | 0 | 1 | 0 |
| 3 | 1 | 0 | 0 | 0 | 1 | 0 |
| 4 | 1 | 0 | 0 | 0 | 1 | 0 |
| 5 | 0 | 1 | 0 | 0 | 0 | 1 |
| 6 | 1 | 0 | 0 | 0 | 1 | 0 |
| 7 | 1 | 0 | 0 | 0 | 1 | 0 |
| 8 | 1 | 0 | 0 | 0 | 1 | 0 |
| 9 | 1 | 0 | 0 | 0 | 1 | 0 |
| 10 | 1 | 0 | 0 | 0 | 1 | 0 |
| 11 | 1 | 0 | 0 | 0 | 1 | 0 |
| 12 | 1 | 0 | 0 | 0 | 1 | 0 |
| 13 | 1 | 0 | 0 | 0 | 1 | 0 |
| 14 | 1 | 0 | 0 | 0 | 1 | 0 |
| 15 | 1 | 0 | 0 | 0 | 1 | 0 |
| 16 | 1 | 0 | 0 | 0 | 1 | 0 |
| 17 | 1 | 0 | 0 | 0 | 1 | 0 |
| 18 | 1 | 0 | 0 | 0 | 1 | 0 |
| 19 | 1 | 0 | 0 | 0 | 1 | 0 |
| 20 | 1 | 0 | 0 | 0 | 1 | 0 |
| 21 | 1 | 0 | 0 | 0 | 1 | 0 |
| 22 | 1 | 0 | 0 | 0 | 1 | 0 |
| 23 | 0 | 1 | 0 | 0 | 0 | 1 |
| 24 | 1 | 0 | 0 | 0 | 1 | 0 |
| 25 | 1 | 0 | 0 | 0 | 1 | 0 |
| 26 | 1 | 0 | 0 | 0 | 1 | 0 |
| 27 | 1 | 0 | 0 | 0 | 1 | 0 |
| 28 | 1 | 0 | 0 | 0 | 1 | 0 |
| TOTAL | 26 | 2 | 0 | 0 | 26 | 2 |
| PERCENTAGE | 92.9 | 7.1 | 0.0 | 0.0 | 92.9 | 7.1 |
| Stat. sig. con. (siRNA) vs. con. (IVM) embryo ‡p<0.05, ‡‡p<0.005 | | | | | | |
| p-value (2-tailed students t-test) | 8.36E-01 | 8.36E-01 | - | - | 8.36E-01 | 8.36E-01 |

| Control siRNA microinjected GV stage (+18h IBMX & 16h post IBMX wash-out) | | | | | | |
|---|---|---|---|---|---|---|
| # | Meiotic spindle status | | | | 1st meiotic polar body (PB1) | |
| | Normal MII arrested (+PB1) | Normal MI stage (no PB1) | Defective (e.g. dispersed chromosomes, no PB1) | Absent with ultra condensed chromatin | Present | Absent |
| 1 | 0 | 0 | 1 | 0 | 0 | 1 |
| 2 | 0 | 0 | 1 | 0 | 0 | 1 |
| 3 | 0 | 0 | 1 | 0 | 0 | 1 |
| 4 | 1 | 0 | 0 | 0 | 1 | 0 |
| 5 | 0 | 0 | 1 | 0 | 0 | 1 |
| 6 | 0 | 1 | 0 | 0 | 0 | 1 |
| 7 | 0 | 1 | 0 | 0 | 0 | 1 |
| 8 | 0 | 0 | 0 | 1 | 0 | 1 |
| 9 | 0 | 1 | 0 | 0 | 0 | 1 |
| 10 | 0 | 0 | 0 | 1 | 0 | 1 |
| 11 | 0 | 0 | 0 | 1 | 0 | 1 |
| 12 | 0 | 0 | 0 | 1 | 0 | 1 |
| 13 | 0 | 1 | 0 | 0 | 0 | 1 |
| 14 | 0 | 1 | 0 | 0 | 0 | 1 |
| 15 | 0 | 1 | 0 | 0 | 0 | 1 |
| 16 | 1 | 0 | 0 | 0 | 1 | 0 |
| 17 | 0 | 0 | 0 | 1 | 0 | 1 |
| 18 | 0 | 0 | 1 | 0 | 0 | 1 |
| 19 | 0 | 0 | 0 | 1 | 0 | 1 |
| 20 | 0 | 0 | 0 | 1 | 0 | 1 |
| 21 | 0 | 1 | 0 | 0 | 0 | 1 |
| 22 | 0 | 0 | 1 | 0 | 0 | 1 |
| 23 | 0 | 1 | 0 | 0 | 0 | 1 |
| 24 | 0 | 0 | 1 | 0 | 0 | 1 |
| 25 | 0 | 1 | 0 | 0 | 0 | 1 |
| 26 | 0 | 1 | 0 | 0 | 0 | 1 |
| 27 | 0 | 1 | 0 | 0 | 0 | 1 |
| 28 | 0 | 1 | 0 | 0 | 0 | 1 |
| 29 | 0 | 0 | 1 | 0 | 0 | 1 |
| 30 | 0 | 1 | 0 | 0 | 0 | 1 |
| 31 | 0 | 1 | 0 | 0 | 0 | 1 |
| 32 | 1 | 0 | 0 | 0 | 1 | 0 |
| 33 | 0 | 0 | 0 | 1 | 0 | 1 |
| 34 | 0 | 0 | 0 | 1 | 0 | 1 |
| 35 | 0 | 1 | 0 | 0 | 0 | 1 |
| TOTAL | 3 | 15 | 8 | 9 | 3 | 32 |
| PERCENTAGE | 8.6 | 42.9 | 22.9 | 25.7 | 8.6 | 91.4 |
| Stat. sig. exp. vs. con. (IVM) embryo ‡p<0.05, ‡‡p<0.005 | ‡‡ | ‡‡ | ‡ | ‡ | ‡‡ | ‡‡ |
| p-value (2-tailed students t-test) | 4.37E-15 | 4.44E-03 | 2.78E-02 | 1.78E-02 | 4.37E-15 | 4.37E-15 |
| Stat. sig. exp. vs. con. (siRNA) embryo *p<0.05, **p<0.005 | ** | ** | * | ** | ** | ** |
| p-value (2-tailed students t-test) | 7.26E-18 | 1.17E-03 | 6.22E-03 | 3.25E-03 | 7.26E-18 | 7.26E-18 |

Supplementary Tables ST15

#### Control siRNA microinjected GV stage (+18h IBMX & extended 24h post IBMX wash-out)

| # | Meiotic spindle status | | | | | 1st meiotic polar body (PB1) | |
|---|---|---|---|---|---|---|---|
| | Normal MII arrested (+PB1) | Normal MI stage (no PB1) | Defective (e.g. dispersed chromosomes, no PB1) | Absent with ultra condensed chromatin | Absent after failed MI | Present | Absent |
| **1** | 0 | 1 | 0 | 0 | 0 | 0 | 1 |
| **2** | 1 | 0 | 0 | 0 | 0 | 1 | 0 |
| **3** | 1 | 0 | 0 | 0 | 0 | 1 | 0 |
| **4** | 1 | 0 | 0 | 0 | 0 | 1 | 0 |
| **5** | 1 | 0 | 0 | 0 | 0 | 1 | 0 |
| **6** | 1 | 0 | 0 | 0 | 0 | 1 | 0 |
| **7** | 1 | 0 | 0 | 0 | 0 | 1 | 0 |
| **8** | 0 | 1 | 0 | 0 | 0 | 0 | 1 |
| **9** | 1 | 0 | 0 | 0 | 0 | 1 | 0 |
| **10** | 0 | 1 | 0 | 0 | 0 | 0 | 1 |
| **11** | 0 | 0 | 1 | 0 | 0 | 0 | 1 |
| **12** | 1 | 0 | 0 | 0 | 0 | 1 | 0 |
| **13** | 1 | 0 | 0 | 0 | 0 | 1 | 0 |
| **14** | 1 | 0 | 0 | 0 | 0 | 1 | 0 |
| **15** | 1 | 0 | 0 | 0 | 0 | 1 | 0 |
| **16** | 1 | 0 | 0 | 0 | 0 | 1 | 0 |
| **17** | 1 | 0 | 0 | 0 | 0 | 1 | 0 |
| **18** | 1 | 0 | 0 | 0 | 0 | 1 | 0 |
| **19** | 1 | 0 | 0 | 0 | 0 | 1 | 0 |
| **20** | 1 | 0 | 0 | 0 | 0 | 1 | 0 |
| **21** | 1 | 0 | 0 | 0 | 0 | 1 | 0 |
| **22** | 1 | 0 | 0 | 0 | 0 | 1 | 0 |
| **23** | 0 | 1 | 0 | 0 | 0 | 0 | 1 |
| **24** | 1 | 0 | 0 | 0 | 0 | 1 | 0 |
| **25** | 1 | 0 | 0 | 0 | 0 | 1 | 0 |
| **26** | 1 | 0 | 0 | 0 | 0 | 1 | 0 |
| **27** | 0 | 1 | 0 | 0 | 0 | 0 | 1 |
| **TOTAL** | **21** | **5** | **1** | **0** | **0** | **21** | **6** |
| **PERCENTAGE** | **77.8** | **18.5** | **3.7** | **0.0** | **0.0** | **77.8** | **22.2** |

#### *Wwc2* siRNA microinjected GV stage (+18h IBMX & extended 24h post IBMX wash-out)

| # | Meiotic spindle status | | | | | 1st meiotic polar body (PB1) | |
|---|---|---|---|---|---|---|---|
| | Normal MII arrested (+PB1) | Normal MI stage (no PB1) | Defective (e.g. dispersed chromosomes, no PB1) | Absent with ultra condensed chromatin | Absent after failed MI | Present | Absent |
| **1** | 0 | 0 | 1 | 0 | 0 | 0 | 1 |
| **2** | 0 | 1 | 0 | 0 | 0 | 0 | 1 |
| **3** | 0 | 0 | 1 | 0 | 0 | 0 | 1 |
| **4** | 0 | 0 | 1 | 0 | 0 | 0 | 1 |
| **5** | 0 | 0 | 0 | 0 | 1 | 0 | 1 |
| **6** | 0 | 0 | 0 | 0 | 1 | 0 | 1 |
| **7** | 0 | 0 | 0 | 0 | 1 | 0 | 1 |
| **8** | 0 | 0 | 0 | 0 | 1 | 0 | 1 |
| **9** | 0 | 1 | 0 | 0 | 0 | 0 | 1 |
| **10** | 0 | 0 | 1 | 0 | 0 | 0 | 1 |
| **11** | 0 | 0 | 0 | 0 | 1 | 0 | 1 |
| **12** | 0 | 0 | 0 | 1 | 0 | 0 | 1 |
| **13** | 1 | 0 | 0 | 0 | 0 | 1 | 0 |
| **14** | 1 | 0 | 0 | 0 | 0 | 1 | 0 |
| **15** | 0 | 0 | 0 | 0 | 1 | 0 | 1 |
| **16** | 0 | 0 | 0 | 0 | 1 | 0 | 1 |
| **17** | 0 | 0 | 0 | 0 | 1 | 0 | 1 |
| **18** | 0 | 0 | 1 | 0 | 0 | 0 | 1 |
| **19** | 0 | 0 | 1 | 0 | 0 | 0 | 1 |
| **20** | 0 | 0 | 1 | 0 | 0 | 0 | 1 |
| **21** | 0 | 0 | 0 | 0 | 1 | 0 | 1 |
| **22** | 0 | 0 | 0 | 1 | 0 | 0 | 1 |
| **23** | 0 | 1 | 0 | 0 | 0 | 0 | 1 |
| **24** | 0 | 1 | 0 | 0 | 0 | 0 | 1 |
| **25** | 0 | 0 | 0 | 0 | 1 | 0 | 1 |
| **26** | 0 | 0 | 0 | 1 | 0 | 0 | 1 |
| **27** | 0 | 0 | 0 | 0 | 1 | 0 | 1 |
| **28** | 0 | 0 | 1 | 0 | 0 | 0 | 1 |
| **29** | 0 | 0 | 0 | 0 | 1 | 0 | 1 |
| **30** | 0 | 0 | 0 | 0 | 1 | 0 | 1 |
| **31** | 0 | 1 | | 0 | 0 | 0 | 1 |
| **32** | 0 | 0 | 0 | 1 | 0 | 0 | 1 |
| **TOTAL** | **2** | **5** | **8** | **4** | **13** | **2** | **30** |
| **PERCENTAGE** | **6.3** | **15.6** | **25.0** | **12.5** | **40.6** | **6.3** | **93.8** |
| Stat. sig. *Wwc2* KD vs. con. siRNA embryo ‡p<0.05, ‡‡p<0.005 | **‡‡** | **‡‡** | **‡** | | **‡‡** | **‡‡** | **‡‡** |
| p-value (2-tailed students t-test) | 5.07E-11 | 7.73E-01 | 2.01E-02 | 5.85E-02 | 8.72E-05 | 5.07E-11 | 5.07E-11 |

**Supplementary Tables ST16**

#### Control siRNA microinjected GV stage (+18h IBMX & 1h post IBMX wash-out)

| # | Meiotic spindle status | | | | | | 1st meiotic polar body (PB1) | |
|---|---|---|---|---|---|---|---|---|
| | Normal MII arrested (+PB1) | Normal MI stage (no PB1) | Defective (e.g. dispersed chromosomes, no PB1) | Absent with ultra condensed chromatin | GV intact | GVBD (no spindle) | Present | Absent |
| **1** | 0 | 0 | 0 | 0 | 1 | 0 | 0 | 1 |
| **2** | 0 | 0 | 0 | 0 | 0 | 1 | 0 | 1 |
| **3** | 0 | 0 | 0 | 0 | 0 | 1 | 0 | 1 |
| **4** | 0 | 0 | 0 | 0 | 0 | 1 | 0 | 1 |
| **5** | 0 | 0 | 0 | 0 | 0 | 1 | 0 | 1 |
| **6** | 0 | 0 | 0 | 0 | 0 | 1 | 0 | 1 |
| **7** | 0 | 0 | 0 | 0 | 0 | 1 | 0 | 1 |
| **8** | 0 | 0 | 0 | 0 | 0 | 1 | 0 | 1 |
| **9** | 0 | 0 | 0 | 0 | 0 | 1 | 0 | 1 |
| **10** | 0 | 0 | 0 | 0 | 0 | 1 | 0 | 1 |
| **11** | 0 | 0 | 0 | 0 | 0 | 1 | 0 | 1 |
| **12** | 0 | 0 | 0 | 0 | 0 | 1 | 0 | 1 |
| **13** | 0 | 0 | 0 | 0 | 0 | 1 | 0 | 1 |
| **14** | 0 | 0 | 0 | 0 | 0 | 1 | 0 | 1 |
| **TOTAL** | **0** | **0** | **0** | **0** | **1** | **13** | **0** | **14** |
| **PERCENTAGE** | **0.0** | **0.0** | **0.0** | **0.0** | **7.1** | **92.9** | **0.0** | **100.0** |

#### *Wwc2* siRNA microinjected GV stage (+18h IBMX & 1h post IBMX wash-out)

| # | Meiotic spindle status | | | | | | 1st meiotic polar body (PB1) | |
|---|---|---|---|---|---|---|---|---|
| | Normal MII arrested (+PB1) | Normal MI stage (no PB1) | Defective (e.g. dispersed chromosomes, no PB1) | Absent with ultra condensed chromatin | GV intact | GVBD (no spindle) | Present | Absent |
| **1** | 0 | 0 | 0 | 0 | 1 | 0 | 0 | 1 |
| **2** | 0 | 0 | 0 | 0 | 0 | 1 | 0 | 1 |
| **3** | 0 | 0 | 0 | 0 | 1 | 0 | 0 | 1 |
| **4** | 0 | 0 | 0 | 0 | 1 | 0 | 0 | 1 |
| **5** | 0 | 0 | 0 | 0 | 1 | 0 | 0 | 1 |
| **6** | 0 | 0 | 0 | 0 | 0 | 1 | 0 | 1 |
| **7** | 0 | 0 | 0 | 0 | 0 | 1 | 0 | 1 |
| **8** | 0 | 0 | 0 | 0 | 0 | 1 | 0 | 1 |
| **9** | 0 | 0 | 0 | 1 | 0 | 0 | 0 | 1 |
| **10** | 0 | 0 | 0 | 0 | 1 | 0 | 0 | 1 |
| **11** | 0 | 0 | 0 | 0 | 0 | 1 | 0 | 1 |
| **12** | 0 | 0 | 0 | 0 | 1 | 0 | 0 | 1 |
| **13** | 0 | 0 | 0 | 0 | 0 | 1 | 0 | 1 |
| **14** | 0 | 0 | 0 | 0 | 0 | 1 | 0 | 1 |
| **15** | 0 | 0 | 0 | 0 | 0 | 1 | 0 | 1 |
| **TOTAL** | **0** | **0** | **0** | **1** | **6** | **8** | **0** | **15** |
| **PERCENTAGE** | **0.0** | **0.0** | **0.0** | **6.7** | **40.0** | **53.3** | **0.0** | **100.0** |
| Stat. sig. *Wwc2* siRNA vs. con. siRNA embryo ‡p<0.05, ‡‡p<0.005 | | | | | ‡ | ‡ | | |
| p-value (2-tailed students t-test) | n/a | n/a | n/a | 3.43E-01 | 3.99E-02 | 1.64E-02 | n/a | n/a |

#### *Wwc2* siRNA + *Wwc2*-HA mRNA microinjected GV stage (+18h IBMX & 1h post IBMX wash-out)

| # | Meiotic spindle status | | | | | | 1st meiotic polar body (PB1) | |
|---|---|---|---|---|---|---|---|---|
| | Normal MII arrested (+PB1) | Normal MI stage (no PB1) | Defective (e.g. dispersed chromosomes, no PB1) | Absent with ultra condensed chromatin | GV intact | GVBD (no spindle) | Present | Absent |
| **1** | 0 | 0 | 0 | 0 | 1 | 0 | 0 | 1 |
| **2** | 0 | 0 | 0 | 0 | 0 | 1 | 0 | 1 |
| **3** | 0 | 0 | 0 | 0 | 0 | 1 | 0 | 1 |
| **4** | 0 | 0 | 0 | 0 | 0 | 1 | 0 | 1 |
| **5** | 0 | 0 | 0 | 0 | 0 | 1 | 0 | 1 |
| **6** | 0 | 0 | 0 | 0 | 0 | 1 | 0 | 1 |
| **7** | 0 | 0 | 0 | 0 | 0 | 1 | 0 | 1 |
| **8** | 0 | 0 | 0 | 0 | 0 | 1 | 0 | 1 |
| **9** | 0 | 0 | 0 | 0 | 0 | 1 | 0 | 1 |
| **10** | 0 | 0 | 0 | 0 | 0 | 1 | 0 | 1 |
| **11** | 0 | 0 | 0 | 0 | 0 | 1 | 0 | 1 |
| **12** | 0 | 0 | 0 | 0 | 0 | 1 | 0 | 1 |
| **13** | 0 | 0 | 0 | 0 | 0 | 1 | 0 | 1 |
| **14** | 0 | 0 | 0 | 0 | 0 | 1 | 0 | 1 |
| **15** | 0 | 0 | 0 | 0 | 0 | 1 | 0 | 1 |
| **16** | 0 | 0 | 0 | 0 | 1 | 0 | 0 | 1 |
| **TOTAL** | **0** | **0** | **0** | **0** | **2** | **14** | **0** | **16** |
| **PERCENTAGE** | **0.0** | **0.0** | **0.0** | **0.0** | **12.5** | **87.5** | **0.0** | **100.0** |
| Stat. sig. *Wwc2* KD (rescue) vs. con. siRNA embryo ‡p<0.05, ‡‡p<0.005 | | | | | | | | |
| p-value (2-tailed students t-test) | n/a | n/a | n/a | n/a | 6.40E-01 | 6.40E-01 | n/a | n/a |
| Stat. sig. *Wwc2* KD (rescue) vs. *Wwc2* siRNA embryo *p<0.05, **p<0.005 | | | | | | * | | |
| p-value (2-tailed students t-test) | n/a | n/a | n/a | 3.10E-01 | 8.53E-02 | 3.70E-02 | n/a | n/a |

Supplementary Tables ST17

#### Control siRNA microinjected GV stage (+18h IBMX & 6h post IBMX wash-out)

| # | Meiotic spindle status | | | | | | 1st meiotic polar body (PB1) | |
|---|---|---|---|---|---|---|---|---|
| | Normal MII arrested (+PB1) | Normal MI stage (no PB1) | Defective (e.g. dispersed chromosomes, no PB1) | Absent with ultra condensed chromatin | GV intact | GVBD (no spindle) | Present | Absent |
| **1** | 0 | 1 | 0 | 0 | 0 | 0 | 0 | 1 |
| **2** | 0 | 0 | 1 | 0 | 0 | 0 | 0 | 1 |
| **3** | 0 | 1 | 0 | 0 | 0 | 0 | 0 | 1 |
| **4** | 0 | 1 | 0 | 0 | 0 | 0 | 0 | 1 |
| **5** | 0 | 1 | 0 | 0 | 0 | 0 | 0 | 1 |
| **6** | 0 | 1 | 0 | 0 | 0 | 0 | 0 | 1 |
| **7** | 0 | 1 | 0 | 0 | 0 | 0 | 0 | 1 |
| **8** | 0 | 1 | 0 | 0 | 0 | 0 | 0 | 1 |
| **9** | 0 | 0 | 1 | 0 | 0 | 0 | 0 | 1 |
| **10** | 0 | 1 | 0 | 0 | 0 | 0 | 0 | 1 |
| **11** | 0 | 1 | 0 | 0 | 0 | 0 | 0 | 1 |
| **12** | 0 | 1 | 0 | 0 | 0 | 0 | 0 | 1 |
| **13** | 0 | 1 | 0 | 0 | 0 | 0 | 0 | 1 |
| **14** | 0 | 1 | 0 | 0 | 0 | 0 | 0 | 1 |
| **15** | 0 | 1 | 0 | 0 | 0 | 0 | 0 | 1 |
| **TOTAL** | **0** | **13** | **2** | **0** | **0** | **0** | **0** | **15** |
| **PERCENTAGE** | **0.0** | **86.7** | **13.3** | **0.0** | **0.0** | **0.0** | **0.0** | **100.0** |

#### *Wwc2* siRNA microinjected GV stage (+18h IBMX & 6h post IBMX wash-out)

| # | Meiotic spindle status | | | | | | 1st meiotic polar body (PB1) | |
|---|---|---|---|---|---|---|---|---|
| | Normal MII arrested (+PB1) | Normal MI stage (no PB1) | Defective (e.g. dispersed chromosomes, no PB1) | Absent with ultra condensed chromatin | GV intact | GVBD (no spindle) | Present | Absent |
| **1** | 0 | 0 | 1 | 0 | 0 | 0 | 0 | 1 |
| **2** | 0 | 1 | 0 | 0 | 0 | 0 | 0 | 1 |
| **3** | 0 | 0 | 1 | 0 | 0 | 0 | 0 | 1 |
| **4** | 0 | 0 | 1 | 0 | 0 | 0 | 0 | 1 |
| **5** | 0 | 1 | 0 | 0 | 0 | 0 | 0 | 1 |
| **6** | 0 | 0 | 1 | 0 | 0 | 0 | 0 | 1 |
| **7** | 0 | 0 | 1 | 0 | 0 | 0 | 0 | 1 |
| **8** | 0 | 0 | 0 | 1 | 0 | 0 | 0 | 1 |
| **9** | 0 | 1 | 0 | 0 | 0 | 0 | 0 | 1 |
| **10** | 0 | 0 | 0 | 1 | 0 | 0 | 0 | 1 |
| **11** | 0 | 1 | 0 | 0 | 0 | 0 | 0 | 1 |
| **12** | 0 | 1 | 0 | 0 | 0 | 0 | 0 | 1 |
| **13** | 0 | 0 | 0 | 0 | 1 | 0 | 0 | 1 |
| **14** | 0 | 1 | 0 | 0 | 0 | 0 | 0 | 1 |
| **15** | 0 | 0 | 0 | 0 | 1 | 0 | 0 | 1 |
| **16** | 0 | 0 | 0 | 0 | 1 | 0 | 0 | 1 |
| **17** | 0 | 1 | 0 | 0 | 0 | 0 | 0 | 1 |
| **18** | 0 | 1 | 0 | 0 | 0 | 0 | 0 | 1 |
| **TOTAL** | **0** | **8** | **5** | **2** | **3** | **0** | **0** | **18** |
| **PERCENTAGE** | **0.0** | **44.4** | **27.8** | **11.1** | **16.7** | **0.0** | **0.0** | **100.0** |
| Stat. sig. *Wwc2* siRNA vs. con. siRNA embryo ‡p<0.05, ‡‡p<0.005 | | ‡ | | | | | | |
| p-value (2-tailed students t-test) | n/a | 1.10E-02 | 3.27E-01 | 1.94E-01 | 1.03E-01 | n/a | n/a | n/a |

#### *Wwc2* siRNA + *Wwc2*-HA mRNA microinjected GV stage (+18h IBMX & 6h post IBMX wash-out)

| # | Meiotic spindle status | | | | | | 1st meiotic polar body (PB1) | |
|---|---|---|---|---|---|---|---|---|
| | Normal MII arrested (+PB1) | Normal MI stage (no PB1) | Defective (e.g. dispersed chromosomes, no PB1) | Absent with ultra condensed chromatin | GV intact | GVBD (no spindle) | Present | Absent |
| **1** | 0 | 1 | 0 | 0 | 0 | 0 | 0 | 1 |
| **2** | 0 | 1 | 0 | 0 | 0 | 0 | 0 | 1 |
| **3** | 0 | 1 | 0 | 0 | 0 | 0 | 0 | 1 |
| **4** | 0 | 1 | 0 | 0 | 0 | 0 | 0 | 1 |
| **5** | 0 | 1 | 0 | 0 | 0 | 0 | 0 | 1 |
| **6** | 0 | 1 | 0 | 0 | 0 | 0 | 0 | 1 |
| **7** | 0 | 1 | 0 | 0 | 0 | 0 | 0 | 1 |
| **8** | 0 | 1 | 0 | 0 | 0 | 0 | 0 | 1 |
| **9** | 0 | 0 | 1 | 0 | 0 | 0 | 0 | 1 |
| **10** | 0 | 0 | 0 | 0 | 1 | 0 | 0 | 1 |
| **11** | 0 | 1 | 0 | 0 | 0 | 0 | 0 | 1 |
| **12** | 0 | 1 | 0 | 0 | 0 | 0 | 0 | 1 |
| **13** | 0 | 1 | 0 | 0 | 0 | 0 | 0 | 1 |
| **14** | 0 | 1 | 0 | 0 | 0 | 0 | 0 | 1 |
| **15** | 0 | 1 | 0 | 0 | 0 | 0 | 0 | 1 |
| **16** | 0 | 0 | 1 | 0 | 0 | 0 | 0 | 1 |
| **17** | 0 | 1 | 0 | 0 | 0 | 0 | 0 | 1 |
| **18** | 0 | 1 | 0 | 0 | 0 | 0 | 0 | 1 |
| **TOTAL** | **0** | **15** | **2** | **0** | **1** | **0** | **0** | **18** |
| **PERCENTAGE** | **0.0** | **83.3** | **11.1** | **0.0** | **5.6** | **0.0** | **0.0** | **100.0** |
| Stat. sig. *Wwc2* KD (rescue) vs. con. siRNA embryo ‡p<0.05, ‡‡p<0.005 | | ‡‡ | ‡‡ | | ‡‡ | | | |
| p-value (2-tailed students t-test) | n/a | 7.98E-01 | 8.51E-01 | n/a | 3.70E-01 | n/a | n/a | n/a |
| Stat. sig. *Wwc2* KD (rescue) vs. *Wwc2* siRNA embryo *p<0.05, **p<0.005 | | * | | | | | | |
| p-value (2-tailed students t-test) | n/a | 1.43E-02 | 2.18E-01 | 1.54E-01 | 3.02E-01 | n/a | n/a | n/a |

**Supplementary Tables ST18**

#### Control siRNA microinjected GV stage (+18h IBMX & 12h post IBMX wash-out)

| # | Meiotic spindle status | | | | | | $1^{st}$ meiotic polar body (PB1) | |
|---|---|---|---|---|---|---|---|---|
| | Normal MII arrested (+PB1) | Normal MI stage (no PB1) | Defective (e.g. dispersed chromosomes, no PB1) | Absent with ultra condensed chromatin | GV intact | GVBD (no spindle) | Present | Absent |
| 1 | 1 | 0 | 0 | 0 | 0 | 0 | 1 | 0 |
| 2 | 1 | 0 | 0 | 0 | 0 | 0 | 1 | 0 |
| 3 | 0 | 1 | 0 | 0 | 0 | 0 | 0 | 1 |
| 4 | 0 | 1 | 0 | 0 | 0 | 0 | 0 | 1 |
| 5 | 1 | 0 | 0 | 0 | 0 | 0 | 1 | 0 |
| 6 | 1 | 0 | 0 | 0 | 0 | 0 | 1 | 0 |
| 7 | 1 | 0 | 0 | 0 | 0 | 0 | 1 | 0 |
| 8 | 1 | 0 | 0 | 0 | 0 | 0 | 1 | 0 |
| 9 | 1 | 0 | 0 | 0 | 0 | 0 | 1 | 0 |
| 10 | 1 | 0 | 0 | 0 | 0 | 0 | 1 | 0 |
| 11 | 1 | 0 | 0 | 0 | 0 | 0 | 1 | 0 |
| 12 | 0 | 1 | 0 | 0 | 0 | 0 | 0 | 1 |
| 13 | 1 | 0 | 0 | 0 | 0 | 0 | 1 | 0 |
| 14 | 1 | 0 | 0 | 0 | 0 | 0 | 1 | 0 |
| 15 | 0 | 1 | 0 | 0 | 0 | 0 | 0 | 1 |
| **TOTAL** | **11** | **4** | **0** | **0** | **0** | **0** | **11** | **4** |
| **PERCENTAGE** | **73.3** | **26.7** | **0.0** | **0.0** | **0.0** | **0.0** | **73.3** | **26.7** |

#### *Wwc2* siRNA microinjected GV stage (+18h IBMX & 12h post IBMX wash-out)

| # | Meiotic spindle status | | | | | | $1^{st}$ meiotic polar body (PB1) | |
|---|---|---|---|---|---|---|---|---|
| | Normal MII arrested (+PB1) | Normal MI stage (no PB1) | Defective (e.g. dispersed chromosomes, no PB1) | Absent with ultra condensed chromatin | GV intact | GVBD (no spindle) | Present | Absent |
| 1 | 0 | 0 | 0 | 1 | 0 | 0 | 0 | 1 |
| 2 | 1 | 0 | 0 | 0 | 0 | 0 | 1 | 0 |
| 3 | 0 | 0 | 1 | 0 | 0 | 0 | 0 | 1 |
| 4 | 0 | 0 | 1 | 0 | 0 | 0 | 0 | 1 |
| 5 | 0 | 1 | 0 | 0 | 0 | 0 | 0 | 1 |
| 6 | 0 | 0 | 1 | 0 | 0 | 0 | 0 | 1 |
| 7 | 0 | 1 | 0 | 0 | 0 | 0 | 0 | 1 |
| 8 | 0 | 1 | 0 | 0 | 0 | 0 | 0 | 1 |
| 9 | 0 | 0 | 1 | 0 | 0 | 0 | 0 | 1 |
| 10 | 0 | 1 | 0 | 0 | 0 | 0 | 0 | 1 |
| 11 | 0 | 0 | 0 | 1 | 0 | 0 | 0 | 1 |
| 12 | 1 | 0 | 0 | 0 | 0 | 0 | 1 | 0 |
| 13 | 0 | 1 | 0 | 0 | 0 | 0 | 0 | 1 |
| 14 | 0 | 1 | 0 | 0 | 0 | 0 | 0 | 1 |
| 15 | 0 | 1 | 0 | 0 | 0 | 0 | 0 | 1 |
| 16 | 0 | 1 | 0 | 0 | 0 | 0 | 0 | 1 |
| 17 | 0 | 0 | 1 | 0 | 0 | 0 | 0 | 1 |
| 18 | 0 | 0 | 0 | 1 | 0 | 0 | 0 | 1 |
| **TOTAL** | **2** | **8** | **5** | **3** | **0** | **0** | **2** | **16** |
| **PERCENTAGE** | **11.1** | **44.4** | **27.8** | **16.7** | **0.0** | **0.0** | **11.1** | **88.9** |
| Stat. sig. *Wwc2* siRNA vs. con. siRNA embryo ‡p<0.05, ‡‡p<0.005 | ‡‡ | | ‡ | | | | ‡‡ | ‡‡ |
| p-value (2-tailed students t-test) | 7.43E-05 | 3.05E-01 | 2.66E-02 | 1.03E-01 | n/a | n/a | 7.43E-05 | 7.43E-05 |

#### *Wwc2* siRNA + *Wwc2*-HA mRNA microinjected GV stage (+18h IBMX & 12h post IBMX wash-out)

| # | Meiotic spindle status | | | | | | $1^{st}$ meiotic polar body (PB1) | |
|---|---|---|---|---|---|---|---|---|
| | Normal MII arrested (+PB1) | Normal MI stage (no PB1) | Defective (e.g. dispersed chromosomes, no PB1) | Absent with ultra condensed chromatin | GV intact | GVBD (no spindle) | Present | Absent |
| 1 | 0 | 1 | 0 | 0 | 0 | 0 | 0 | 1 |
| 2 | 1 | 0 | 0 | 0 | 0 | 0 | 1 | 0 |
| 3 | 0 | 1 | 0 | 0 | 0 | 0 | 0 | 1 |
| 4 | 1 | 0 | 0 | 0 | 0 | 0 | 1 | 0 |
| 5 | 1 | 0 | 0 | 0 | 0 | 0 | 1 | 0 |
| 6 | 0 | 1 | 0 | 0 | 0 | 0 | 0 | 1 |
| 7 | 1 | 0 | 0 | 0 | 0 | 0 | 1 | 0 |
| 8 | 1 | 0 | 0 | 0 | 0 | 0 | 1 | 0 |
| 9 | 1 | 0 | 0 | 0 | 0 | 0 | 1 | 0 |
| 10 | 1 | 0 | 0 | 0 | 0 | 0 | 1 | 0 |
| 11 | 1 | 0 | 0 | 0 | 0 | 0 | 1 | 0 |
| 12 | 1 | 0 | 0 | 0 | 0 | 0 | 1 | 0 |
| 13 | 1 | 0 | 0 | 0 | 0 | 0 | 1 | 0 |
| 14 | 1 | 0 | 0 | 0 | 0 | 0 | 1 | 0 |
| 15 | 0 | 0 | 1 | 0 | 0 | 0 | 0 | 1 |
| 16 | 0 | 0 | 1 | 0 | 0 | 0 | 0 | 1 |
| 17 | 1 | 0 | 0 | 0 | 0 | 0 | 1 | 0 |
| **TOTAL** | **12** | **3** | **2** | **0** | **0** | **0** | **12** | **5** |
| **PERCENTAGE** | **70.6** | **17.6** | **11.8** | **0.0** | **0.0** | **0.0** | **70.6** | **29.4** |
| Stat. sig. *Wwc2* KD (rescue) vs. con. siRNA embryo ‡p<0.05, ‡‡p<0.005 | | | | | | | | |
| p-value (2-tailed students t-test) | 8.69E-01 | 5.53E-01 | 1.81E-01 | n/a | n/a | n/a | 8.69E-01 | 8.69E-01 |
| Stat. sig. *Wwc2* KD (rescue) vs. *Wwc2* siRNA embryo *p<0.05, **p<0.005 | ** | | | | | | ** | ** |
| p-value (2-tailed students t-test) | 1.11E-04 | 9.28E-02 | 2.49E-01 | 8.26E-02 | n/a | n/a | 1.11E-04 | 1.11E-04 |

**Supplementary Tables ST19**

#### Control siRNA microinjected GV stage (+18h IBMX & 16h post IBMX wash-out)

| # | Meiotic spindle status | | | | | | 1st meiotic polar body (PB1) | |
|---|---|---|---|---|---|---|---|---|
| | Normal MII arrested (+PB1) | Normal MI stage (no PB1) | Defective (e.g. dispersed chromosomes, no PB1) | Absent with ultra condensed chromatin | GV intact | GVBD (no spindle) | Present | Absent |
| **1** | 0 | 1 | 0 | 0 | 0 | 0 | 0 | 1 |
| **2** | 1 | 0 | 0 | 0 | 0 | 0 | 1 | 0 |
| **3** | 1 | 0 | 0 | 0 | 0 | 0 | 1 | 0 |
| **4** | 1 | 0 | 0 | 0 | 0 | 0 | 1 | 0 |
| **5** | 1 | 0 | 0 | 0 | 0 | 0 | 1 | 0 |
| **6** | 1 | 0 | 0 | 0 | 0 | 0 | 1 | 0 |
| **7** | 1 | 0 | 0 | 0 | 0 | 0 | 1 | 0 |
| **8** | 1 | 0 | 0 | 0 | 0 | 0 | 1 | 0 |
| **9** | 1 | 0 | 0 | 0 | 0 | 0 | 1 | 0 |
| **10** | 1 | 0 | 0 | 0 | 0 | 0 | 1 | 0 |
| **11** | 1 | 0 | 0 | 0 | 0 | 0 | 1 | 0 |
| **12** | 1 | 0 | 0 | 0 | 0 | 0 | 1 | 0 |
| **13** | 1 | 0 | 0 | 0 | 0 | 0 | 1 | 0 |
| **14** | 0 | 1 | 0 | 0 | 0 | 0 | 0 | 1 |
| **15** | 0 | 1 | 0 | 0 | 0 | 0 | 0 | 1 |
| **16** | 1 | 0 | 0 | 0 | 0 | 0 | 1 | 0 |
| **17** | 0 | 0 | 1 | 0 | 0 | 0 | 0 | 1 |
| **18** | 1 | 0 | 0 | 0 | 0 | 0 | 1 | 0 |
| **19** | 1 | 0 | 0 | 0 | 0 | 0 | 1 | 0 |
| **20** | 1 | 0 | 0 | 0 | 0 | 0 | 1 | 0 |
| **21** | 1 | 0 | 0 | 0 | 0 | 0 | 1 | 0 |
| **22** | 1 | 0 | 0 | 0 | 0 | 0 | 1 | 0 |
| **TOTAL** | **18** | **3** | **1** | **0** | **0** | **0** | **18** | **4** |
| **PERCENTAGE** | **81.8** | **13.6** | **4.5** | **0.0** | **0.0** | **0.0** | **81.8** | **18.2** |

#### *Wwc2* siRNA microinjected GV stage (+18h IBMX & 16h post IBMX wash-out)

| # | Meiotic spindle status | | | | | | 1st meiotic polar body (PB1) | |
|---|---|---|---|---|---|---|---|---|
| | Normal MII arrested (+PB1) | Normal MI stage (no PB1) | Defective (e.g. dispersed chromosomes, no PB1) | Absent with ultra condensed chromatin | GV intact | GVBD (no spindle) | Present | Absent |
| **1** | 0 | 0 | 1 | 0 | 0 | 0 | 0 | 1 |
| **2** | 0 | 0 | 0 | 1 | 0 | 0 | 0 | 1 |
| **3** | 0 | 0 | 0 | 1 | 0 | 0 | 0 | 1 |
| **4** | 0 | 1 | 0 | 0 | 0 | 0 | 0 | 1 |
| **5** | 0 | 0 | 1 | 0 | 0 | 0 | 0 | 1 |
| **6** | 0 | 1 | 0 | 0 | 0 | 0 | 0 | 1 |
| **7** | 0 | 0 | 0 | 1 | 0 | 0 | 0 | 1 |
| **8** | 0 | 0 | 1 | 0 | 0 | 0 | 0 | 1 |
| **9** | 0 | 0 | 1 | 0 | 0 | 0 | 0 | 1 |
| **10** | 0 | 1 | 0 | 0 | 0 | 0 | 0 | 1 |
| **11** | 0 | 1 | 0 | 0 | 0 | 0 | 0 | 1 |
| **12** | 0 | 1 | 0 | 0 | 0 | 0 | 0 | 1 |
| **13** | 0 | 1 | 0 | 0 | 0 | 0 | 0 | 1 |
| **14** | 1 | 0 | 0 | 0 | 0 | 0 | 1 | 0 |
| **15** | 1 | 0 | 0 | 0 | 0 | 0 | 1 | 0 |
| **16** | 0 | 1 | 0 | 0 | 0 | 0 | 0 | 1 |
| **17** | 0 | 0 | 1 | 0 | 0 | 0 | 0 | 1 |
| **18** | 0 | 1 | 0 | 0 | 0 | 0 | 0 | 1 |
| **19** | 0 | 1 | 0 | 0 | 0 | 0 | 0 | 1 |
| **20** | 0 | 1 | 0 | 0 | 0 | 0 | 0 | 1 |
| **21** | 0 | 0 | 1 | 0 | 0 | 0 | 0 | 1 |
| **22** | 0 | 0 | 0 | 1 | 0 | 0 | 0 | 1 |
| **23** | 0 | 0 | 0 | 1 | 0 | 0 | 0 | 1 |
| **TOTAL** | **2** | **10** | **6** | **5** | **0** | **0** | **2** | **21** |
| **PERCENTAGE** | **8.7** | **43.5** | **26.1** | **21.7** | **0.0** | **0.0** | **8.7** | **91.3** |
| Stat. sig. *Wwc2* siRNA vs. con. siRNA embryo ‡p<0.05, ‡‡p<0.005 | ‡‡ | ‡ | ‡ | ‡ | | | ‡‡ | ‡‡ |
| p-value (2-tailed students t-test) | 8.61E-09 | 2.73E-02 | 4.75E-02 | 2.00E-02 | n/a | n/a | 8.61E-09 | 8.61E-09 |

#### *Wwc2* siRNA + *Wwc2*-HA mRNA microinjected GV stage (+18h IBMX & 16h post IBMX wash-out)

| # | Meiotic spindle status | | | | | | 1st meiotic polar body (PB1) | |
|---|---|---|---|---|---|---|---|---|
| | Normal MII arrested (+PB1) | Normal MI stage (no PB1) | Defective (e.g. dispersed chromosomes, no PB1) | Absent with ultra condensed chromatin | GV intact | GVBD (no spindle) | Present | Absent |
| **1** | 0 | 0 | 1 | 0 | 0 | 0 | 0 | 1 |
| **2** | 1 | 0 | 0 | 0 | 0 | 0 | 1 | 0 |
| **3** | 0 | 1 | 0 | 0 | 0 | 0 | 0 | 1 |
| **4** | 0 | 0 | 1 | 0 | 0 | 0 | 0 | 1 |
| **5** | 1 | 0 | 0 | 0 | 0 | 0 | 1 | 0 |
| **6** | 1 | 0 | 0 | 0 | 0 | 0 | 1 | 0 |
| **7** | 1 | 0 | 0 | 0 | 0 | 0 | 1 | 0 |
| **8** | 0 | 1 | 0 | 0 | 0 | 0 | 0 | 1 |
| **9** | 1 | 0 | 0 | 0 | 0 | 0 | 1 | 0 |
| **10** | 1 | 0 | 0 | 0 | 0 | 0 | 1 | 0 |
| **11** | 0 | 1 | 0 | 0 | 0 | 0 | 0 | 1 |
| **12** | 1 | 0 | 0 | 0 | 0 | 0 | 1 | 0 |
| **13** | 1 | 0 | 0 | 0 | 0 | 0 | 1 | 0 |
| **14** | 1 | 0 | 0 | 0 | 0 | 0 | 1 | 0 |
| **15** | 1 | 0 | 0 | 0 | 0 | 0 | 1 | 0 |
| **16** | 0 | 1 | 0 | 0 | 0 | 0 | 0 | 1 |
| **17** | 1 | 0 | 0 | 0 | 0 | 0 | 1 | 0 |
| **18** | 1 | 0 | 0 | 0 | 0 | 0 | 1 | 0 |
| **19** | 1 | 0 | 0 | 0 | 0 | 0 | 1 | 0 |
| **20** | 1 | 0 | 0 | 0 | 0 | 0 | 1 | 0 |
| **21** | 1 | 0 | 0 | 0 | 0 | 0 | 1 | 0 |
| **22** | 1 | 0 | 0 | 0 | 0 | 0 | 1 | 0 |
| **23** | 1 | 0 | 0 | 0 | 0 | 0 | 1 | 0 |
| **24** | 1 | 0 | 0 | 0 | 0 | 0 | 1 | 0 |
| **25** | 1 | 0 | 0 | 0 | 0 | 0 | 1 | 0 |
| **TOTAL** | **19** | **4** | **2** | **0** | **0** | **0** | **19** | **6** |
| **PERCENTAGE** | **76.0** | **16.0** | **8.0** | **0.0** | **0.0** | **0.0** | **76.0** | **24.0** |
| Stat. sig. *Wwc2* KD (rescue) vs. con. siRNA embryo ‡p<0.05, ‡‡p<0.005 | | | | | | | | |
| p-value (2-tailed students t-test) | 6.36E-01 | 8.25E-01 | 6.38E-01 | #DIV/0! | n/a | n/a | 6.36E-01 | 6.36E-01 |
| Stat. sig. *Wwc2* KD (rescue) vs. *Wwc2* siRNA embryo *p<0.05, **p<0.005 | ** | * | | * | | | ** | ** |
| p-value (2-tailed students t-test) | 1.21E-07 | 3.70E-02 | 9.68E-02 | 1.31E-02 | n/a | n/a | 1.21E-07 | 1.21E-07 |

**Supplementary Tables ST20**

#### Control siRNA (10μM), 1in2 micro-injection, 32-cell (E3.5) stage, anti-CDX2

| # | Total cell number per embryo | Cells in injected clone | Cells in non-injected clone | INJECTED CLONE | | | | | | | NON INJECTED CLONE | | | | | | |
|---|---|---|---|---|---|---|---|---|---|---|---|---|---|---|---|---|---|
| | | | | Nuclear abnormality | OUTER CELLS | | | INNER CELLS | | | Nuclear abnormality | OUTER CELLS | | | INNER CELLS | | |
| | | | | | | CDX2+VE | CDX2-VE | | CDX2+VE | CDX2-VE | | | CDX2+VE | CDX2-VE | | CDX2+VE | CDX2-VE |
| **1** | 33 | 16 | 17 | 0 | 9 | 9 | 0 | 7 | 0 | 7 | 0 | 9 | 9 | 0 | 8 | 0 | 8 |
| **2** | 31 | 16 | 15 | 0 | 11 | 11 | 0 | 5 | 0 | 5 | 0 | 9 | 9 | 0 | 6 | 0 | 6 |
| **3** | 31 | 15 | 16 | 0 | 8 | 8 | 0 | 7 | 0 | 7 | 0 | 8 | 8 | 0 | 8 | 0 | 8 |
| **4** | 39 | 20 | 19 | 0 | 11 | 11 | 0 | 9 | 0 | 9 | 0 | 12 | 12 | 0 | 7 | 0 | 7 |
| **5** | 32 | 15 | 17 | 0 | 9 | 9 | 0 | 6 | 0 | 6 | 0 | 8 | 8 | 0 | 9 | 0 | 9 |
| **6** | 39 | 22 | 17 | 0 | 14 | 14 | 0 | 8 | 0 | 8 | 0 | 12 | 12 | 0 | 5 | 0 | 5 |
| **7** | 33 | 17 | 16 | 0 | 12 | 12 | 0 | 5 | 0 | 5 | 0 | 9 | 9 | 0 | 7 | 0 | 7 |
| **8** | 41 | 21 | 20 | 0 | 12 | 12 | 0 | 9 | 0 | 9 | 0 | 11 | 11 | 0 | 9 | 0 | 9 |
| **9** | 32 | 16 | 16 | 0 | 10 | 10 | 0 | 6 | 0 | 6 | 0 | 9 | 9 | 0 | 7 | 0 | 7 |
| **10** | 32 | 16 | 16 | 0 | 8 | 8 | 0 | 8 | 0 | 8 | 0 | 8 | 8 | 0 | 8 | 0 | 8 |
| **11** | 36 | 18 | 18 | 0 | 10 | 10 | 0 | 8 | 0 | 8 | 0 | 11 | 11 | 0 | 7 | 0 | 7 |
| **12** | 42 | 22 | 20 | 0 | 13 | 13 | 0 | 9 | 0 | 9 | 0 | 12 | 12 | 0 | 8 | 0 | 8 |
| **13** | 44 | 24 | 20 | 0 | 16 | 16 | 0 | 8 | 0 | 8 | 0 | 14 | 14 | 0 | 6 | 0 | 6 |
| **14** | 35 | 18 | 17 | 0 | 11 | 11 | 0 | 7 | 0 | 7 | 0 | 9 | 9 | 0 | 8 | 0 | 8 |
| **TOTAL** | **500** | **256** | **244** | **0** | **154** | **154** | **0** | **102** | **0** | **102** | **0** | **141** | **141** | **0** | **103** | **0** | **103** |
| **AVERAGE** | **35.7** | **18.3** | **17.4** | **0.0** | **11.0** | **11.0** | **0.0** | **7.3** | **0.0** | **7.3** | **0.0** | **10.1** | **10.1** | **0.0** | **7.4** | **0.0** | **7.4** |
| SEM | **1.0** | **0.7** | **0.4** | **0.0** | **0.5** | **0.5** | **0.0** | **0.4** | **0.0** | **0.4** | **0.0** | **0.4** | **0.4** | **0.0** | **0.3** | **0.0** | **0.3** |
| Stat. sig. (inter-clone) ‡p<0.05, ‡‡p<0.005 | | | | | | | | | | | | | | | | | |
| p-value (2-tailed students t-test) | | 3.57E-01 | | 1.00E+00 | 2.53E-01 | 2.53E-01 | 1.00E+00 | 8.83E-01 | 1.00E+00 | 8.83E-01 | | | | | | | |

#### Wwc2-specific siRNA (10μM), 1in2 micro-injection, 32-cell (E3.5) stage, anti-CDX2

| # | Total cell number per embryo | Cells in injected clone | Cells in non-injected clone | INJECTED CLONE | | | | | | | NON INJECTED CLONE | | | | | | |
|---|---|---|---|---|---|---|---|---|---|---|---|---|---|---|---|---|---|
| | | | | Nuclear abnormality | OUTER CELLS | | | INNER CELLS | | | Nuclear abnormality | OUTER CELLS | | | INNER CELLS | | |
| | | | | | | CDX2+VE | CDX2-VE | | CDX2+VE | CDX2-VE | | | CDX2+VE | CDX2-VE | | CDX2+VE | CDX2-VE |
| **1** | 33 | 11 | 22 | 1 | 9 | 6 | 3 | 2 | 0 | 2 | 0 | 14 | 14 | 0 | 8 | 0 | 8 |
| **2** | 32 | 13 | 19 | 2 | 10 | 8 | 2 | 3 | 0 | 3 | 0 | 12 | 12 | 0 | 7 | 0 | 7 |
| **3** | 29 | 10 | 19 | 0 | 9 | 7 | 2 | 1 | 0 | 1 | 0 | 13 | 13 | 0 | 6 | 0 | 6 |
| **4** | 29 | 11 | 18 | 0 | 9 | 6 | 3 | 2 | 0 | 2 | 0 | 11 | 11 | 0 | 7 | 0 | 7 |
| **5** | 34 | 12 | 22 | 1 | 10 | 10 | 0 | 2 | 0 | 2 | 0 | 13 | 13 | 0 | 9 | 0 | 9 |
| **6** | 28 | 10 | 18 | 0 | 10 | 8 | 2 | 0 | 0 | 0 | 0 | 12 | 12 | 0 | 6 | 0 | 6 |
| **7** | 35 | 11 | 24 | 0 | 9 | 6 | 3 | 2 | 0 | 2 | 0 | 16 | 16 | 0 | 8 | 0 | 8 |
| **8** | 28 | 10 | 18 | 2 | 7 | 7 | 0 | 3 | 0 | 3 | 0 | 12 | 12 | 0 | 6 | 0 | 6 |
| **9** | 25 | 9 | 16 | 1 | 8 | 6 | 2 | 1 | 0 | 1 | 0 | 11 | 11 | 0 | 5 | 0 | 5 |
| **10** | 25 | 10 | 15 | 0 | 10 | 9 | 1 | 0 | 0 | 0 | 0 | 9 | 9 | 0 | 6 | 0 | 6 |
| **11** | 29 | 14 | 15 | 2 | 12 | 9 | 3 | 2 | 0 | 2 | 0 | 9 | 9 | 0 | 6 | 0 | 6 |
| **12** | 29 | 13 | 16 | 1 | 10 | 8 | 2 | 3 | 0 | 3 | 0 | 8 | 8 | 0 | 8 | 0 | 8 |
| **13** | 34 | 14 | 20 | 2 | 12 | 9 | 3 | 2 | 0 | 2 | 0 | 13 | 13 | 0 | 7 | 0 | 7 |
| **14** | 30 | 11 | 19 | 0 | 11 | 10 | 1 | 0 | 0 | 0 | 0 | 11 | 11 | 0 | 8 | 0 | 8 |
| **15** | 31 | 13 | 18 | 0 | 12 | 8 | 4 | 1 | 0 | 1 | 0 | 12 | 12 | 0 | 6 | 0 | 6 |
| **16** | 35 | 11 | 24 | 2 | 9 | 6 | 3 | 2 | 0 | 2 | 0 | 16 | 16 | 0 | 8 | 0 | 8 |
| **17** | 26 | 9 | 17 | 0 | 8 | 8 | 0 | 1 | 0 | 1 | 0 | 10 | 10 | 0 | 7 | 0 | 7 |
| **18** | 28 | 11 | 17 | 1 | 9 | 7 | 2 | 2 | 0 | 2 | 0 | 12 | 12 | 0 | 5 | 0 | 5 |
| **TOTAL** | **540** | **203** | **337** | **15** | **174** | **138** | **36** | **29** | **0** | **29** | **0** | **214** | **214** | **0** | **123** | **0** | **123** |
| **AVERAGE** | **30.0** | **11.3** | **18.7** | **0.8** | **9.7** | **7.7** | **2.0** | **1.6** | **0.0** | **1.6** | **0.0** | **11.9** | **11.9** | **0.0** | **6.8** | **0.0** | **6.8** |
| SEM | **0.8** | **0.4** | **0.7** | **0.2** | **0.3** | **0.3** | **0.3** | **0.2** | **0.0** | **0.2** | **0.0** | **0.5** | **0.5** | **0.0** | **0.3** | **0.0** | **0.3** |
| Stat. sig. (inter-clone) ‡p<0.05, ‡‡p<0.005 | | ‡‡ | | ‡‡ | ‡‡ | ‡‡ | ‡‡ | ‡‡ | | ‡‡ | | | | | | | |
| p-value (2-tailed students t-test) | | 1.32E-11 | | 2.27E-04 | 8.86E-04 | 4.66E-08 | 2.96E-08 | 2.86E-16 | 1.00E+00 | 2.86E-16 | | | | | | | |
| Stat. sig. (exp. vs. con embryo) *p<0.05, **p<0.005 | ** | ** | | ** | * | ** | ** | ** | | ** | | * | * | | | | |
| p-value (2-tailed students t-test) | 2.18E-04 | 1.32E-09 | 1.34E-01 | 1.06E-03 | 5.14E-02 | 1.64E-05 | 6.48E-07 | 2.30E-14 | 1.00E+00 | 2.30E-14 | 1.00E+00 | 1.89E-02 | 1.89E-02 | 1.00E+00 | 2.11E-01 | 1.00E+00 | 2.11E-01 |

**Supplementary Tables ST21**

#### Control siRNA (10μM), 1in2 micro-injection, >64-cell (E4.5) stage, anti-CDX2 & anti-NANOG

| # | Total cell number per embryo | Cells in injected clone | Cells in non-injected clone | INJECTED CLONE | | | | | | | | | | | | | NON INJECTED CLONE | | | | | | | | | | | | |
|---|---|---|---|---|---|---|---|---|---|---|---|---|---|---|---|---|---|---|---|---|---|---|---|---|---|---|---|---|---|
| | | | | Nuclear abnormality | OUTER CELLS | | | | | | INNER CELLS | | | | | | Nuclear abnormality | OUTER CELLS | | | | | | INNER CELLS | | | | | |
| | | | | | | CDX2+VE | CDX2-VE | CDX2 & NANOG+VE | NANOG-VE | Apoptotic ♣ | | CDX2+VE | CDX2-VE | NANOG+VE | NANOG-VE | Apoptotic ♣ | | | CDX2+VE | CDX2-VE | CDX2 & NANOG+VE | NANOG-VE | Apoptotic ♣ | | CDX2+VE | CDX2-VE | NANOG+VE | NANOG-VE | Apoptotic ♣ |
| **1** | 77 | 37 | 40 | 0 | 26 | 25 | 0 | 1 | 24 | 1 | 11 | 0 | 11 | 8 | 3 | 1 | 0 | 32 | 30 | 0 | 2 | 28 | 2 | 8 | 0 | 8 | 6 | 2 | 1 |
| **2** | 83 | 40 | 43 | 0 | 29 | 29 | 0 | 0 | 29 | 2 | 11 | 0 | 11 | 9 | 2 | 0 | 0 | 33 | 32 | 0 | 1 | 31 | 1 | 10 | 0 | 10 | 5 | 5 | 1 |
| **3** | 71 | 34 | 37 | 0 | 23 | 21 | 0 | 2 | 19 | 0 | 11 | 0 | 11 | 7 | 4 | 1 | 0 | 29 | 29 | 0 | 0 | 29 | 1 | 8 | 0 | 8 | 5 | 3 | 0 |
| **4** | 84 | 42 | 42 | 0 | 32 | 32 | 0 | 0 | 32 | 0 | 10 | 0 | 10 | 7 | 3 | 1 | 0 | 26 | 25 | 0 | 1 | 24 | 2 | 16 | 0 | 16 | 10 | 6 | 1 |
| **5** | 77 | 38 | 39 | 0 | 28 | 26 | 0 | 2 | 24 | 1 | 10 | 0 | 10 | 8 | 2 | 0 | 0 | 30 | 30 | 0 | 0 | 30 | 2 | 9 | 0 | 9 | 7 | 2 | 0 |
| **6** | 81 | 40 | 41 | 0 | 27 | 26 | 0 | 1 | 25 | 0 | 13 | 0 | 13 | 9 | 4 | 0 | 0 | 35 | 33 | 0 | 2 | 31 | 0 | 6 | 0 | 6 | 3 | 3 | 2 |
| **7** | 84 | 43 | 41 | 0 | 33 | 32 | 0 | 1 | 31 | 1 | 10 | 0 | 10 | 6 | 4 | 2 | 0 | 29 | 29 | 0 | 0 | 29 | 1 | 12 | 0 | 12 | 8 | 4 | 0 |
| **8** | 86 | 43 | 43 | 0 | 34 | 34 | 0 | 0 | 34 | 1 | 9 | 0 | 9 | 6 | 3 | 1 | 0 | 30 | 27 | 0 | 3 | 24 | 2 | 13 | 0 | 13 | 10 | 3 | 0 |
| **9** | 84 | 41 | 43 | 0 | 29 | 27 | 0 | 2 | 25 | 2 | 12 | 0 | 12 | 9 | 3 | 0 | 0 | 34 | 33 | 0 | 1 | 32 | 1 | 9 | 0 | 9 | 7 | 2 | 0 |
| **10** | 77 | 40 | 37 | 0 | 30 | 29 | 0 | 1 | 28 | 1 | 10 | 0 | 10 | 6 | 4 | 2 | 0 | 27 | 27 | 0 | 0 | 27 | 1 | 10 | 0 | 10 | 8 | 2 | 1 |
| **11** | 79 | 41 | 38 | 0 | 32 | 30 | 0 | 2 | 28 | 0 | 9 | 0 | 9 | 6 | 3 | 1 | 0 | 28 | 26 | 0 | 2 | 24 | 2 | 10 | 0 | 10 | 9 | 1 | 0 |
| **TOTAL** | **883** | **439** | **444** | **0** | **323** | **311** | **0** | **12** | **299** | **9** | **116** | **0** | **116** | **81** | **35** | **9** | **0** | **333** | **321** | **0** | **12** | **309** | **15** | **111** | **0** | **111** | **78** | **33** | **6** |
| **AVERAGE** | **80.3** | **39.9** | **40.4** | **0.0** | **29.4** | **28.3** | **0.0** | **1.1** | **27.2** | **0.8** | **10.5** | **0.0** | **10.5** | **7.4** | **3.2** | **0.8** | **0.0** | **30.3** | **29.2** | **0.0** | **1.1** | **28.1** | **1.4** | **10.1** | **0.0** | **10.1** | **7.1** | **3.0** | **0.5** |
| SEM | **1.4** | **0.8** | **0.7** | **0.0** | **1.0** | **1.1** | **0.0** | **0.3** | **1.3** | **0.2** | **0.4** | **0.0** | **0.4** | **0.4** | **0.2** | **0.2** | **0.0** | **0.9** | **0.8** | **0.0** | **0.3** | **0.9** | **0.2** | **0.8** | **0.0** | **0.8** | **0.7** | **0.4** | **0.2** |
| Stat. sig. (inter-clone) ‡p<0.05, ‡‡p<0.005 | | | | | | | | | | | | | | | | | | | | | | | | | | | | | |
| p-value (2-tailed students t-test) | | 6.77E-01 | | 1.00E+00 | 5.00E-01 | 5.24E-01 | 1.00E+00 | 1.00E+00 | 5.71E-01 | 8.81E-02 | 6.20E-01 | 1.00E+00 | 6.20E-01 | 7.27E-01 | 7.21E-01 | 3.85E-01 | | | | | | | | | | | | | |

#### Wwc2-specific siRNA (10μM), 1in2 micro-injection, >64-cell (E4.5) stage, anti-CDX2 & anti-NANOG

| # | Total cell number per embryo | Cells in injected clone | Cells in non-injected clone | INJECTED CLONE | | | | | | | | | | | | | NON INJECTED CLONE | | | | | | | | | | | | |
|---|---|---|---|---|---|---|---|---|---|---|---|---|---|---|---|---|---|---|---|---|---|---|---|---|---|---|---|---|---|
| | | | | Nuclear abnormality | OUTER CELLS | | | | | | INNER CELLS | | | | | | Nuclear abnormality | OUTER CELLS | | | | | | INNER CELLS | | | | | |
| | | | | | | CDX2+VE | CDX2-VE | CDX2 & NANOG+VE | NANOG-VE | Apoptotic ♣ | | CDX2+VE | CDX2-VE | NANOG+VE | NANOG-VE | Apoptotic ♣ | | | CDX2+VE | CDX2-VE | CDX2 & NANOG+VE | NANOG-VE | Apoptotic ♣ | | CDX2+VE | CDX2-VE | NANOG+VE | NANOG-VE | Apoptotic ♣ |
| **1** | 55 | 22 | 33 | 4 | 18 | 16 | 0 | 2 | 16 | 0 | 4 | 0 | 4 | 2 | 2 | 3 | 0 | 23 | 22 | 0 | 1 | 22 | 2 | 10 | 0 | 10 | 8 | 2 | 0 |
| **2** | 55 | 20 | 35 | 3 | 15 | 13 | 2 | 0 | 15 | 1 | 5 | 0 | 5 | 1 | 4 | 4 | 0 | 24 | 23 | 0 | 1 | 23 | 1 | 11 | 0 | 11 | 7 | 4 | 1 |
| **3** | 54 | 17 | 37 | 0 | 12 | 5 | 3 | 4 | 8 | 2 | 5 | 0 | 5 | 3 | 2 | 2 | 0 | 25 | 23 | 0 | 2 | 23 | 1 | 12 | 0 | 12 | 9 | 3 | 0 |
| **4** | 65 | 23 | 42 | 2 | 20 | 16 | 3 | 1 | 19 | 0 | 3 | 0 | 3 | 0 | 3 | 5 | 0 | 28 | 27 | 0 | 1 | 27 | 3 | 14 | 0 | 14 | 10 | 4 | 0 |
| **5** | 63 | 20 | 43 | 2 | 16 | 11 | 4 | 1 | 15 | 0 | 4 | 0 | 4 | 1 | 3 | 2 | 0 | 30 | 27 | 0 | 3 | 27 | 2 | 13 | 0 | 13 | 7 | 6 | 1 |
| **6** | 65 | 26 | 39 | 4 | 23 | 21 | 2 | 0 | 23 | 1 | 3 | 0 | 3 | 1 | 2 | 4 | 0 | 27 | 26 | 0 | 1 | 26 | 2 | 12 | 0 | 12 | 9 | 3 | 0 |
| **7** | 65 | 23 | 42 | 2 | 20 | 18 | 0 | 2 | 18 | 1 | 3 | 0 | 3 | 0 | 3 | 3 | 0 | 26 | 26 | 0 | 0 | 26 | 0 | 16 | 0 | 16 | 14 | 2 | 2 |
| **8** | 66 | 26 | 40 | 3 | 22 | 18 | 3 | 1 | 21 | 2 | 4 | 0 | 4 | 1 | 3 | 2 | 0 | 28 | 27 | 0 | 1 | 27 | 1 | 12 | 0 | 12 | 8 | 4 | 0 |
| **9** | 58 | 18 | 40 | 2 | 16 | 15 | 0 | 1 | 15 | 1 | 2 | 0 | 2 | 2 | 0 | 5 | 0 | 30 | 28 | 0 | 2 | 28 | 2 | 10 | 0 | 10 | 7 | 3 | 2 |
| **10** | 63 | 22 | 41 | 4 | 18 | 16 | 0 | 2 | 16 | 1 | 4 | 0 | 4 | 1 | 3 | 2 | 0 | 32 | 32 | 0 | 0 | 32 | 1 | 9 | 0 | 9 | 7 | 2 | 0 |
| **11** | 65 | 22 | 43 | 3 | 17 | 13 | 4 | 0 | 17 | 0 | 5 | 0 | 5 | 2 | 3 | 1 | 0 | 30 | 29 | 0 | 1 | 29 | 2 | 13 | 0 | 13 | 8 | 5 | 1 |
| **12** | 67 | 25 | 42 | 4 | 21 | 13 | 6 | 2 | 19 | 2 | 4 | 0 | 4 | 2 | 2 | 3 | 0 | 28 | 28 | 0 | 0 | 28 | 2 | 14 | 0 | 14 | 9 | 5 | 0 |
| **13** | 62 | 23 | 39 | 3 | 18 | 15 | 0 | 3 | 15 | 2 | 5 | 0 | 5 | 3 | 2 | 2 | 0 | 26 | 24 | 0 | 2 | 24 | 1 | 13 | 0 | 13 | 8 | 5 | 0 |
| **14** | 53 | 20 | 33 | 4 | 16 | 14 | 1 | 1 | 15 | 1 | 4 | 0 | 4 | 2 | 2 | 4 | 0 | 23 | 23 | 0 | 0 | 23 | 1 | 10 | 0 | 10 | 8 | 2 | 1 |
| **15** | 75 | 26 | 49 | 3 | 23 | 18 | 5 | 0 | 23 | 2 | 3 | 0 | 3 | 0 | 3 | 5 | 0 | 35 | 32 | 0 | 3 | 32 | 1 | 14 | 0 | 14 | 9 | 5 | 0 |
| **16** | 64 | 22 | 42 | 0 | 18 | 17 | 0 | 1 | 17 | 1 | 4 | 0 | 4 | 1 | 3 | 3 | 0 | 29 | 28 | 0 | 1 | 28 | 1 | 13 | 0 | 13 | 9 | 4 | 1 |
| **17** | 63 | 23 | 40 | 3 | 20 | 18 | 2 | 0 | 20 | 2 | 3 | 0 | 3 | 0 | 3 | 3 | 0 | 27 | 25 | 0 | 2 | 25 | 3 | 13 | 0 | 13 | 11 | 2 | 1 |
| **18** | 63 | 20 | 43 | 2 | 16 | 15 | 0 | 1 | 15 | 1 | 4 | 0 | 4 | 1 | 3 | 2 | 0 | 30 | 30 | 0 | 0 | 30 | 1 | 13 | 0 | 13 | 7 | 6 | 0 |
| **19** | 65 | 26 | 39 | 4 | 23 | 17 | 4 | 2 | 21 | 2 | 3 | 0 | 3 | 1 | 2 | 3 | 0 | 27 | 26 | 0 | 1 | 26 | 1 | 12 | 0 | 12 | 9 | 3 | 0 |
| **TOTAL** | **1186** | **424** | **762** | **52** | **352** | **289** | **39** | **24** | **328** | **22** | **72** | **0** | **72** | **24** | **48** | **58** | **0** | **528** | **506** | **0** | **22** | **506** | **28** | **234** | **0** | **234** | **164** | **70** | **10** |
| **AVERAGE** | **62.4** | **22.3** | **40.1** | **2.7** | **18.5** | **15.2** | **2.1** | **1.3** | **17.3** | **1.2** | **3.8** | **0.0** | **3.8** | **1.3** | **2.5** | **3.1** | **0.0** | **27.8** | **26.6** | **0.0** | **1.2** | **26.6** | **1.5** | **12.3** | **0.0** | **12.3** | **8.6** | **3.7** | **0.5** |
| SEM | **1.2** | **0.6** | **0.9** | **0.3** | **0.7** | **0.8** | **0.4** | **0.3** | **0.8** | **0.2** | **0.2** | **0.0** | **0.2** | **0.2** | **0.2** | **0.3** | **0.0** | **0.7** | **0.7** | **0.0** | **0.2** | **0.7** | **0.2** | **0.4** | **0.0** | **0.4** | **0.4** | **0.3** | **0.2** |
| Stat. sig. (inter-clone) ‡p<0.05, ‡‡p<0.005 | | ‡‡ | | ‡‡ | ‡‡ | ‡‡ | ‡‡ | | ‡‡ | | ‡‡ | | ‡‡ | ‡‡ | ‡‡ | ‡‡ | | | | | | | | | | | | | |
| p-value (2-tailed students t-test) | | 1.82E-18 | | 1.73E-11 | 3.66E-11 | 3.77E-13 | 5.52E-05 | 7.55E-01 | 1.32E-10 | 2.13E-01 | 1.61E-20 | 1.00E+00 | 1.61E-20 | 2.24E-18 | 3.47E-03 | 1.45E-09 | | | | | | | | | | | | | |
| Stat. sig. (exp. vs. con embryo) *p<0.05, **p<0.005 | ** | ** | | ** | ** | ** | ** | | ** | | ** | | ** | ** | * | ** | | * | * | | | * | | * | | * | * | | |
| p-value (2-tailed students t-test) | 4.80E-10 | 2.16E-16 | 8.40E-01 | 6.57E-08 | 7.58E-10 | 1.74E-10 | 1.78E-03 | 6.56E-01 | 2.11E-07 | 2.48E-01 | 7.62E-17 | 1.00E+00 | 7.62E-17 | 6.44E-15 | 4.16E-02 | 4.80E-06 | 1.00E+00 | 3.72E-02 | 2.56E-02 | 1.00E+00 | 8.60E-01 | 2.00E-01 | 6.97E-01 | 1.07E-02 | 1.00E+00 | 1.07E-02 | 4.14E-02 | 2.13E-01 | 9.42E-01 |

**Supplementary Tables ST22**

#### Control siRNA (10μM), 1in2 micro-injection, >64-cell (E4.5) stage, anti-GATA4 & anti-NANOG

| # | Total cell number per embryo | Cells in injected clone | Cells in non-injected clone | INJECTED CLONE | | | | | | | | | | | | | | | NON INJECTED CLONE | | | | | | | | | | | | | | |
|---|---|---|---|---|---|---|---|---|---|---|---|---|---|---|---|---|---|---|---|---|---|---|---|---|---|---|---|---|---|---|---|---|---|
| | | | | Nuclear abnormality | OUTER CELLS | GATA4+VE | GATA4-VE | NANOG+VE | NANOG-VE | GATA4 & NANOG -VE | Apoptotic ♣ | INNER CELLS | GATA4+VE | GATA4-VE | NANOG+VE | NANOG-VE | GATA4 & NANOG -VE | Apoptotic ♣ | Nuclear abnormality | OUTER CELLS | GATA4+VE | GATA4-VE | NANOG+VE | NANOG-VE | GATA4 & NANOG -VE | Apoptotic ♣ | INNER CELLS | GATA4+VE | GATA4-VE | NANOG+VE | NANOG-VE | GATA4 & NANOG -VE | Apoptotic ♣ |
| **1** | 80 | 38 | 42 | 0 | 26 | 0 | 26 | 2 | 24 | 24 | 1 | 12 | 3 | 9 | 9 | 3 | 0 | 0 | 0 | 32 | 0 | 32 | 1 | 31 | 31 | 2 | 10 | 2 | 8 | 8 | 2 | 0 | 1 |
| **2** | 88 | 45 | 43 | 0 | 34 | 0 | 34 | 1 | 33 | 33 | 3 | 11 | 4 | 7 | 7 | 4 | 0 | 2 | 0 | 29 | 0 | 29 | 3 | 26 | 26 | 2 | 14 | 5 | 9 | 9 | 5 | 0 | 2 |
| **3** | 84 | 41 | 43 | 0 | 28 | 0 | 28 | 0 | 28 | 28 | 2 | 13 | 6 | 7 | 7 | 6 | 0 | 1 | 0 | 30 | 0 | 30 | 0 | 30 | 30 | 2 | 13 | 3 | 10 | 10 | 3 | 0 | 1 |
| **4** | 91 | 46 | 45 | 0 | 36 | 0 | 36 | 2 | 34 | 34 | 2 | 10 | 4 | 6 | 6 | 4 | 0 | 1 | 0 | 29 | 0 | 29 | 1 | 28 | 28 | 2 | 16 | 5 | 11 | 11 | 5 | 0 | 3 |
| **5** | 87 | 43 | 44 | 0 | 29 | 0 | 29 | 1 | 28 | 28 | 1 | 14 | 3 | 11 | 11 | 3 | 0 | 1 | 0 | 33 | 0 | 33 | 2 | 31 | 31 | 2 | 11 | 5 | 6 | 6 | 5 | 0 | 2 |
| **6** | 80 | 38 | 42 | 0 | 27 | 0 | 27 | 2 | 25 | 25 | 2 | 11 | 3 | 8 | 8 | 3 | 0 | 1 | 0 | 32 | 0 | 32 | 1 | 31 | 31 | 3 | 10 | 4 | 6 | 6 | 4 | 0 | 2 |
| **7** | 84 | 43 | 41 | 0 | 33 | 0 | 33 | 0 | 33 | 33 | 2 | 10 | 4 | 6 | 6 | 4 | 0 | 2 | 0 | 28 | 0 | 28 | 0 | 28 | 28 | 1 | 13 | 3 | 10 | 10 | 3 | 0 | 0 |
| **8** | 81 | 41 | 40 | 0 | 26 | 0 | 26 | 2 | 24 | 24 | 1 | 15 | 3 | 12 | 12 | 3 | 0 | 0 | 0 | 32 | 0 | 32 | 3 | 29 | 29 | 2 | 8 | 2 | 6 | 6 | 2 | 0 | 1 |
| **9** | 83 | 43 | 40 | 0 | 30 | 0 | 30 | 2 | 28 | 28 | 3 | 13 | 4 | 9 | 9 | 4 | 0 | 1 | 0 | 29 | 0 | 29 | 1 | 28 | 28 | 1 | 11 | 3 | 8 | 8 | 3 | 0 | 1 |
| **10** | 83 | 42 | 41 | 0 | 31 | 0 | 31 | 0 | 31 | 31 | 1 | 11 | 4 | 7 | 7 | 4 | 0 | 0 | 0 | 27 | 0 | 27 | 0 | 27 | 27 | 2 | 14 | 3 | 11 | 11 | 3 | 0 | 2 |
| **11** | 84 | 40 | 44 | 0 | 30 | 0 | 30 | 3 | 27 | 27 | 2 | 10 | 2 | 8 | 8 | 2 | 0 | 1 | 0 | 31 | 0 | 31 | 2 | 29 | 29 | 1 | 13 | 3 | 10 | 10 | 3 | 0 | 2 |
| **12** | 78 | 38 | 40 | 0 | 27 | 0 | 27 | 3 | 24 | 24 | 3 | 11 | 2 | 9 | 9 | 2 | 0 | 1 | 0 | 29 | 0 | 29 | 0 | 29 | 29 | 1 | 11 | 4 | 7 | 7 | 4 | 0 | 1 |
| **13** | 83 | 41 | 42 | 0 | 28 | 0 | 28 | 1 | 27 | 27 | 1 | 13 | 5 | 8 | 8 | 5 | 0 | 1 | 0 | 32 | 0 | 32 | 1 | 31 | 31 | 2 | 10 | 2 | 8 | 8 | 2 | 0 | 0 |
| **14** | 90 | 46 | 44 | 0 | 30 | 0 | 30 | 2 | 28 | 28 | 0 | 16 | 4 | 12 | 12 | 4 | 0 | 1 | 0 | 32 | 0 | 32 | 0 | 32 | 32 | 3 | 12 | 4 | 8 | 8 | 4 | 0 | 1 |
| **15** | 78 | 39 | 39 | 0 | 27 | 0 | 27 | 2 | 25 | 25 | 2 | 12 | 3 | 9 | 9 | 3 | 0 | 2 | 0 | 28 | 0 | 28 | 1 | 27 | 27 | 1 | 11 | 2 | 9 | 9 | 2 | 0 | 1 |
| **16** | 85 | 43 | 42 | 0 | 29 | 0 | 29 | 1 | 28 | 28 | 1 | 14 | 3 | 11 | 11 | 3 | 0 | 0 | 0 | 31 | 0 | 31 | 2 | 29 | 29 | 2 | 11 | 3 | 8 | 8 | 3 | 0 | 0 |
| **17** | 86 | 43 | 43 | 0 | 32 | 0 | 32 | 0 | 32 | 32 | 2 | 11 | 3 | 8 | 8 | 3 | 0 | 1 | 0 | 30 | 0 | 30 | 2 | 28 | 28 | 1 | 13 | 4 | 9 | 9 | 4 | 0 | 0 |
| **TOTAL** | **1425** | **710** | **715** | **0** | **503** | **0** | **503** | **24** | **479** | **479** | **29** | **207** | **60** | **147** | **147** | **60** | **0** | **16** | **0** | **514** | **0** | **514** | **20** | **494** | **494** | **30** | **201** | **57** | **144** | **144** | **57** | **0** | **20** |
| **AVERAGE** | **83.8** | **41.8** | **42.1** | **0.0** | **29.6** | **0.0** | **29.6** | **1.4** | **28.2** | **28.2** | **1.7** | **12.2** | **3.5** | **8.6** | **8.6** | **3.5** | **0.0** | **0.9** | **0.0** | **30.2** | **0.0** | **30.2** | **1.2** | **29.1** | **29.1** | **1.8** | **11.8** | **3.4** | **8.5** | **8.5** | **3.4** | **0.0** | **1.2** |
| SEM | **0.9** | **0.6** | **0.4** | **0.0** | **0.7** | **0.0** | **0.7** | **0.2** | **0.8** | **0.8** | **0.2** | **0.4** | **0.2** | **0.5** | **0.5** | **0.2** | **0.0** | **0.2** | **0.0** | **0.4** | **0.0** | **0.4** | **0.2** | **0.4** | **0.4** | **0.2** | **0.5** | **0.3** | **0.4** | **0.4** | **0.3** | **0.0** | **0.2** |
| Stat. sig. (inter-clone) ‡p<0.05, ‡‡p<0.005 | | ‡‡ | | | | | | | | | | | | | | | | | | | | | | | | | | | | | | | |
| p-value (2-tailed students t-test) | | 7.00E-01 | | 1.00E+00 | 4.36E-01 | 1.00E+00 | 4.36E-01 | 5.02E-01 | 3.40E-01 | 3.40E-01 | 8.23E-01 | 5.88E-01 | 6.22E-01 | 7.73E-01 | 7.73E-01 | 6.22E-01 | 1.00E+00 | 3.85E-01 | | | | | | | | | | | | | | | |

#### Wwc2-specific siRNA (10μM), 1in2 micro-injection, >64-cell (E4.5) stage, anti-GATA4 & anti-NANOG

| # | Total cell number per embryo | Cells in injected clone | Cells in non-injected clone | INJECTED CLONE | | | | | | | | | | | | | | | NON INJECTED CLONE | | | | | | | | | | | | | | |
|---|---|---|---|---|---|---|---|---|---|---|---|---|---|---|---|---|---|---|---|---|---|---|---|---|---|---|---|---|---|---|---|---|---|
| | | | | Nuclear abnormality | OUTER CELLS | GATA4+VE | GATA4-VE | NANOG+VE | NANOG-VE | GATA4 & NANOG -VE | Apoptotic ♣ | INNER CELLS | GATA4+VE | GATA4-VE | NANOG+VE | NANOG-VE | GATA4 & NANOG -VE | Apoptotic ♣ | Nuclear abnormality | OUTER CELLS | GATA4+VE | GATA4-VE | NANOG+VE | NANOG-VE | GATA4 & NANOG -VE | Apoptotic ♣ | INNER CELLS | GATA4+VE | GATA4-VE | NANOG+VE | NANOG-VE | GATA4 & NANOG -VE | Apoptotic ♣ |
| **1** | 74 | 26 | 48 | 2 | 20 | 0 | 20 | 3 | 17 | 17 | 1 | 6 | 2 | 4 | 2 | 4 | 2 | 3 | 0 | 32 | 0 | 32 | 1 | 31 | 31 | 1 | 16 | 7 | 9 | 9 | 7 | 0 | 0 |
| **2** | 63 | 22 | 41 | 1 | 17 | 0 | 17 | 0 | 17 | 17 | 2 | 5 | 1 | 4 | 2 | 3 | 2 | 4 | 0 | 29 | 0 | 29 | 0 | 29 | 29 | 1 | 12 | 4 | 8 | 8 | 4 | 0 | 1 |
| **3** | 68 | 21 | 47 | 3 | 17 | 0 | 17 | 1 | 16 | 16 | 2 | 4 | 1 | 3 | 2 | 2 | 1 | 3 | 0 | 34 | 0 | 34 | 2 | 32 | 32 | 2 | 13 | 2 | 11 | 11 | 2 | 0 | 1 |
| **4** | 61 | 21 | 40 | 2 | 19 | 0 | 19 | 0 | 19 | 19 | 1 | 2 | 2 | 0 | 0 | 2 | 0 | 2 | 0 | 25 | 0 | 25 | 0 | 25 | 25 | 2 | 15 | 3 | 12 | 12 | 3 | 0 | 1 |
| **5** | 58 | 20 | 38 | 1 | 16 | 0 | 16 | 0 | 16 | 16 | 0 | 4 | 2 | 2 | 0 | 4 | 2 | 4 | 0 | 29 | 0 | 29 | 1 | 28 | 28 | 1 | 9 | 2 | 7 | 7 | 2 | 0 | 1 |
| **6** | 63 | 22 | 41 | 2 | 20 | 0 | 20 | 0 | 20 | 20 | 2 | 2 | 0 | 2 | 1 | 1 | 1 | 3 | 0 | 30 | 0 | 30 | 2 | 28 | 28 | 1 | 11 | 5 | 6 | 6 | 5 | 0 | 2 |
| **7** | 61 | 22 | 39 | 1 | 18 | 0 | 18 | 0 | 18 | 18 | 0 | 4 | 1 | 3 | 1 | 3 | 2 | 2 | 0 | 26 | 0 | 26 | 0 | 26 | 26 | 1 | 13 | 4 | 9 | 9 | 4 | 0 | 1 |
| **8** | 77 | 28 | 49 | 0 | 26 | 0 | 26 | 1 | 25 | 25 | 1 | 2 | 0 | 2 | 0 | 2 | 2 | 4 | 0 | 33 | 0 | 33 | 1 | 32 | 32 | 2 | 16 | 5 | 11 | 11 | 5 | 0 | 1 |
| **9** | 69 | 28 | 41 | 2 | 22 | 0 | 22 | 2 | 20 | 20 | 2 | 6 | 3 | 3 | 2 | 4 | 1 | 3 | 0 | 28 | 0 | 28 | 2 | 26 | 26 | 1 | 13 | 4 | 9 | 9 | 4 | 0 | 1 |
| **10** | 69 | 25 | 44 | 3 | 19 | 0 | 19 | 3 | 16 | 16 | 1 | 6 | 3 | 3 | 2 | 4 | 1 | 2 | 0 | 31 | 0 | 31 | 1 | 30 | 30 | 1 | 13 | 3 | 10 | 10 | 3 | 0 | 0 |
| **11** | 68 | 23 | 45 | 0 | 18 | 0 | 18 | 2 | 16 | 16 | 2 | 5 | 2 | 3 | 2 | 3 | 1 | 1 | 0 | 29 | 0 | 29 | 2 | 27 | 27 | 2 | 16 | 5 | 11 | 11 | 5 | 0 | 3 |
| **12** | 65 | 23 | 42 | 2 | 20 | 0 | 20 | 1 | 19 | 19 | 0 | 3 | 3 | 0 | 0 | 3 | 0 | 5 | 0 | 28 | 0 | 28 | 1 | 27 | 27 | 3 | 14 | 4 | 10 | 10 | 4 | 0 | 0 |
| **13** | 65 | 22 | 43 | 2 | 16 | 0 | 16 | 0 | 16 | 16 | 0 | 6 | 3 | 3 | 2 | 4 | 1 | 2 | 0 | 30 | 0 | 30 | 0 | 30 | 30 | 2 | 13 | 6 | 7 | 7 | 6 | 0 | 1 |
| **14** | 67 | 28 | 39 | 4 | 23 | 0 | 23 | 2 | 21 | 21 | 1 | 5 | 2 | 3 | 2 | 3 | 1 | 3 | 0 | 27 | 0 | 27 | 1 | 26 | 26 | 2 | 12 | 3 | 9 | 9 | 3 | 0 | 0 |
| **15** | 65 | 23 | 42 | 2 | 20 | 0 | 20 | 1 | 19 | 19 | 1 | 3 | 2 | 1 | 0 | 3 | 1 | 4 | 0 | 26 | 0 | 26 | 0 | 26 | 26 | 0 | 16 | 2 | 14 | 14 | 2 | 0 | 2 |
| **16** | 66 | 26 | 40 | 3 | 22 | 0 | 22 | 1 | 21 | 21 | 2 | 4 | 3 | 1 | 1 | 3 | 0 | 2 | 0 | 28 | 0 | 28 | 1 | 27 | 27 | 1 | 12 | 4 | 8 | 8 | 4 | 0 | 0 |
| **17** | 73 | 27 | 46 | 3 | 23 | 0 | 23 | 2 | 21 | 21 | 2 | 4 | 2 | 2 | 2 | 2 | 0 | 5 | 0 | 32 | 0 | 32 | 2 | 30 | 30 | 1 | 14 | 5 | 9 | 9 | 5 | 0 | 0 |
| **18** | 63 | 21 | 42 | 1 | 18 | 0 | 18 | 1 | 17 | 17 | 1 | 3 | 1 | 2 | 1 | 2 | 1 | 3 | 0 | 29 | 0 | 29 | 0 | 29 | 29 | 1 | 13 | 4 | 9 | 9 | 4 | 0 | 1 |
| **19** | 64 | 24 | 40 | 1 | 20 | 0 | 20 | 2 | 18 | 18 | 2 | 4 | 2 | 2 | 2 | 2 | 0 | 4 | 0 | 27 | 0 | 27 | 1 | 26 | 26 | 3 | 13 | 2 | 11 | 11 | 2 | 0 | 1 |
| **20** | 62 | 19 | 43 | 3 | 18 | 0 | 18 | 2 | 16 | 16 | 1 | 1 | 0 | 1 | 1 | 0 | 0 | 2 | 0 | 30 | 0 | 30 | 0 | 30 | 30 | 1 | 13 | 6 | 7 | 7 | 6 | 0 | 0 |
| **21** | 67 | 28 | 39 | 2 | 23 | 0 | 23 | 2 | 21 | 21 | 2 | 5 | 4 | 1 | 1 | 4 | 0 | 3 | 0 | 27 | 0 | 27 | 2 | 25 | 25 | 1 | 12 | 3 | 9 | 9 | 3 | 0 | 0 |
| **22** | 68 | 27 | 41 | 1 | 22 | 0 | 22 | 0 | 22 | 22 | 2 | 5 | 3 | 2 | 1 | 4 | 1 | 4 | 0 | 29 | 0 | 29 | 0 | 29 | 29 | 1 | 12 | 4 | 8 | 8 | 4 | 0 | 1 |
| **23** | 71 | 24 | 47 | 3 | 20 | 0 | 20 | 1 | 19 | 19 | 2 | 4 | 2 | 2 | 2 | 2 | 0 | 2 | 0 | 34 | 0 | 34 | 1 | 33 | 33 | 2 | 13 | 2 | 11 | 11 | 2 | 0 | 2 |
| **24** | 60 | 20 | 40 | 2 | 18 | 0 | 18 | 2 | 16 | 16 | 1 | 2 | 1 | 1 | 0 | 2 | 1 | 3 | 0 | 25 | 0 | 25 | 0 | 25 | 25 | 2 | 15 | 3 | 12 | 12 | 3 | 0 | 1 |
| **TOTAL** | **1587** | **570** | **1017** | **46** | **475** | **0** | **475** | **29** | **446** | **446** | **31** | **95** | **45** | **50** | **29** | **66** | **21** | **73** | **0** | **698** | **0** | **698** | **21** | **677** | **677** | **35** | **319** | **92** | **227** | **227** | **92** | **0** | **21** |
| **AVERAGE** | **66.1** | **23.8** | **42.4** | **1.9** | **19.8** | **0.0** | **19.8** | **1.2** | **18.6** | **18.6** | **1.3** | **4.0** | **1.9** | **2.1** | **1.2** | **2.8** | **0.9** | **3.0** | **0.0** | **29.1** | **0.0** | **29.1** | **0.9** | **28.2** | **28.2** | **1.5** | **13.3** | **3.8** | **9.5** | **9.5** | **3.8** | **0.0** | **0.9** |
| SEM | **0.9** | **0.6** | **0.6** | **0.2** | **0.5** | **0.0** | **0.5** | **0.2** | **0.5** | **0.5** | **0.2** | **0.3** | **0.2** | **0.2** | **0.2** | **0.2** | **0.2** | **0.2** | **0.0** | **0.5** | **0.0** | **0.5** | **0.2** | **0.5** | **0.5** | **0.1** | **0.4** | **0.3** | **0.4** | **0.4** | **0.3** | **0.0** | **0.2** |
| Stat. sig. (inter-clone) ‡p<0.05, ‡‡p<0.005 | | ‡‡ | | ‡‡ | ‡‡ | | ‡‡ | | ‡‡ | ‡‡ | | ‡‡ | ‡‡ | ‡‡ | ‡‡ | ‡‡ | ‡‡ | ‡‡ | | | | | | | | | | | | | | | |
| p-value (2-tailed students t-test) | | 1.50E-25 | | 4.98E-12 | 1.77E-16 | 1.00E+00 | 1.77E-16 | 2.02E-01 | 6.29E-18 | 6.29E-18 | 4.37E-01 | 1.68E-24 | 2.09E-06 | 7.89E-21 | 8.97E-24 | 4.31E-03 | 6.06E-07 | 2.17E-10 | | | | | | | | | | | | | | | |
| Stat. sig. (exp. vs. con embryo) *p<0.05, **p<0.005 | ** | ** | | ** | ** | | ** | | ** | ** | | ** | ** | ** | ** | * | ** | ** | | | | | | | | | * | | | | | | |
| p-value (2-tailed students t-test) | 9.70E-16 | 1.93E-22 | 7.08E-01 | 2.15E-09 | 3.49E-14 | 1.00E+00 | 3.49E-14 | 5.20E-01 | 4.06E-13 | 4.06E-13 | 1.07E-01 | 8.46E-19 | 1.34E-05 | 9.22E-17 | 1.12E-19 | 2.40E-02 | 2.00E-05 | 7.66E-09 | 1.00E+00 | 1.23E-01 | 1.00E+00 | 1.23E-01 | 2.93E-01 | 2.20E-01 | 2.20E-01 | 1.74E-01 | 1.51E-02 | 2.41E-01 | 9.11E-02 | 9.11E-02 | 2.41E-01 | 1.00E+00 | 2.61E-01 |

Supplementary Tables ST23

#### Control siRNA (10μM), 1in2 micro-injection, >64-cell (E4.5) stage, anti-GATA4 & anti-SOX2

| # | Total cell number per embryo | Cells in injected clone | Cells in non-injected clone | INJECTED CLONE | | | | | | | | | | | | | | | NON INJECTED CLONE | | | | | | | | | | | | | | |
|---|---|---|---|---|---|---|---|---|---|---|---|---|---|---|---|---|---|---|---|---|---|---|---|---|---|---|---|---|---|---|---|---|---|
| | | | | Nuclear abnormality | OUTER CELLS | | | | | | | INNER CELLS | | | | | | | Nuclear abnormality | OUTER CELLS | | | | | | | INNER CELLS | | | | | | |
| | | | | | | GATA4+VE | GATA4-VE | SOX2+VE | SOX2-VE | GATA4 & SOX2 -VE | Apoptotic ♣ | | GATA4+VE | GATA4-VE | SOX2+VE | SOX2-VE | GATA4 & SOX2 -VE | Apoptotic ♣ | | | GATA4+VE | GATA4-VE | SOX2+VE | SOX2-VE | GATA4 & SOX2 -VE | Apoptotic ♣ | | GATA4+VE | GATA4-VE | SOX2+VE | SOX2-VE | GATA4 & SOX2 -VE | Apoptotic ♣ |
| 1 | 84 | 40 | 44 | 0 | 26 | 0 | 26 | 0 | 26 | 26 | 1 | 14 | 3 | 11 | 11 | 3 | 0 | 0 | 0 | 32 | 0 | 32 | 0 | 32 | 32 | 2 | 12 | 4 | 8 | 8 | 4 | 0 | 1 |
| 2 | 80 | 40 | 40 | 0 | 30 | 0 | 30 | 0 | 30 | 30 | 2 | 10 | 2 | 8 | 8 | 2 | 0 | 0 | 0 | 28 | 0 | 28 | 0 | 28 | 28 | 1 | 12 | 2 | 10 | 10 | 2 | 0 | 1 |
| 3 | 79 | 39 | 40 | 0 | 29 | 0 | 29 | 0 | 29 | 29 | 1 | 10 | 1 | 9 | 9 | 1 | 0 | 1 | 0 | 27 | 0 | 27 | 0 | 27 | 27 | 0 | 13 | 4 | 9 | 9 | 4 | 0 | 2 |
| 4 | 89 | 44 | 45 | 0 | 31 | 0 | 31 | 0 | 31 | 31 | 2 | 13 | 3 | 10 | 10 | 3 | 0 | 2 | 0 | 35 | 0 | 35 | 0 | 35 | 35 | 2 | 10 | 2 | 8 | 8 | 2 | 0 | 1 |
| 5 | 80 | 41 | 39 | 0 | 32 | 0 | 32 | 0 | 32 | 32 | 1 | 9 | 1 | 8 | 8 | 1 | 0 | 1 | 0 | 27 | 0 | 27 | 0 | 27 | 27 | 1 | 12 | 3 | 9 | 9 | 3 | 0 | 3 |
| 6 | 82 | 42 | 40 | 0 | 26 | 0 | 26 | 0 | 26 | 26 | 3 | 16 | 4 | 12 | 12 | 4 | 0 | 0 | 0 | 31 | 0 | 31 | 0 | 31 | 31 | 2 | 9 | 4 | 5 | 5 | 4 | 0 | 1 |
| 7 | 85 | 42 | 43 | 0 | 28 | 0 | 28 | 0 | 28 | 28 | 2 | 14 | 2 | 12 | 12 | 2 | 0 | 1 | 0 | 30 | 0 | 30 | 0 | 30 | 30 | 1 | 13 | 5 | 8 | 8 | 5 | 0 | 1 |
| 8 | 83 | 43 | 40 | 0 | 30 | 0 | 30 | 0 | 30 | 30 | 1 | 13 | 4 | 9 | 9 | 4 | 0 | 2 | 0 | 29 | 0 | 29 | 0 | 29 | 29 | 0 | 11 | 3 | 8 | 8 | 3 | 0 | 0 |
| 9 | 84 | 41 | 43 | 0 | 30 | 0 | 30 | 0 | 30 | 30 | 1 | 11 | 4 | 7 | 7 | 4 | 0 | 0 | 0 | 28 | 0 | 28 | 0 | 28 | 28 | 3 | 15 | 3 | 12 | 12 | 3 | 0 | 2 |
| 10 | 82 | 41 | 41 | 0 | 30 | 0 | 30 | 0 | 30 | 30 | 2 | 11 | 3 | 8 | 8 | 3 | 0 | 2 | 0 | 30 | 0 | 30 | 0 | 30 | 30 | 1 | 11 | 3 | 8 | 8 | 3 | 0 | 0 |
| 11 | 81 | 40 | 41 | 0 | 27 | 0 | 27 | 0 | 27 | 27 | 2 | 13 | 4 | 9 | 9 | 4 | 0 | 1 | 0 | 32 | 0 | 32 | 0 | 32 | 32 | 2 | 9 | 2 | 7 | 7 | 2 | 0 | 1 |
| 12 | 83 | 41 | 42 | 0 | 28 | 0 | 28 | 0 | 28 | 28 | 0 | 13 | 5 | 8 | 8 | 5 | 0 | 1 | 0 | 32 | 0 | 32 | 0 | 32 | 32 | 1 | 10 | 2 | 8 | 8 | 2 | 0 | 0 |
| 13 | 92 | 46 | 46 | 0 | 30 | 0 | 30 | 0 | 30 | 30 | 1 | 16 | 4 | 12 | 12 | 4 | 0 | 0 | 0 | 32 | 0 | 32 | 0 | 32 | 32 | 3 | 14 | 4 | 10 | 10 | 4 | 0 | 3 |
| 14 | 96 | 47 | 49 | 0 | 33 | 0 | 33 | 0 | 33 | 33 | 0 | 14 | 5 | 9 | 9 | 5 | 0 | 1 | 0 | 36 | 0 | 36 | 0 | 36 | 36 | 0 | 13 | 6 | 7 | 7 | 6 | 0 | 1 |
| 15 | 91 | 46 | 45 | 0 | 35 | 0 | 35 | 0 | 35 | 35 | 1 | 11 | 3 | 8 | 8 | 3 | 0 | 0 | 0 | 33 | 0 | 33 | 0 | 33 | 33 | 1 | 12 | 2 | 10 | 10 | 2 | 0 | 0 |
| 16 | 87 | 42 | 45 | 0 | 29 | 0 | 29 | 0 | 29 | 29 | 2 | 13 | 3 | 10 | 10 | 3 | 0 | 2 | 0 | 35 | 0 | 35 | 0 | 35 | 35 | 1 | 10 | 2 | 8 | 8 | 2 | 0 | 2 |
| TOTAL | 1358 | 675 | 683 | 0 | 474 | 0 | 474 | 0 | 474 | 474 | 22 | 201 | 51 | 150 | 150 | 51 | 0 | 14 | 0 | 497 | 0 | 497 | 0 | 497 | 497 | 21 | 186 | 51 | 135 | 135 | 51 | 0 | 19 |
| AVERAGE | 84.9 | 42.2 | 42.7 | 0.0 | 29.6 | 0.0 | 29.6 | 0.0 | 29.6 | 29.6 | 1.4 | 12.6 | 3.2 | 9.4 | 9.4 | 3.2 | 0.0 | 0.9 | 0.0 | 31.1 | 0.0 | 31.1 | 0.0 | 31.1 | 31.1 | 1.3 | 11.6 | 3.2 | 8.4 | 8.4 | 3.2 | 0.0 | 1.2 |
| SEM | 1.2 | 0.6 | 0.7 | 0.0 | 0.6 | 0.0 | 0.6 | 0.0 | 0.6 | 0.6 | 0.2 | 0.5 | 0.3 | 0.4 | 0.4 | 0.3 | 0.0 | 0.2 | 0.0 | 0.7 | 0.0 | 0.7 | 0.0 | 0.7 | 0.7 | 0.2 | 0.4 | 0.3 | 0.4 | 0.4 | 0.3 | 0.0 | 0.2 |
| Stat. sig. (inter-clone) ‡p<0.05, ‡‡p<0.005 | | ‡‡ | | | | | | | | | | | | | | | | | | | | | | | | | | | | | | | |
| p-value (2-tailed students t-test) | | 5.91E-01 | | 1.00E+00 | 1.34E-01 | 1.00E+00 | 1.34E-01 | 1.00E+00 | 1.34E-01 | 1.34E-01 | 8.42E-01 | 1.76E-01 | 1.00E+00 | 1.10E-01 | 1.10E-01 | 1.00E+00 | 1.00E+00 | 3.33E-01 | | | | | | | | | | | | | | | |

#### *Wwc2*-specific siRNA (10μM), 1in2 micro-injection, >64-cell (E4.5) stage, anti-GATA4 & anti-SOX2

| # | Total cell number per embryo | Cells in injected clone | Cells in non-injected clone | INJECTED CLONE | | | | | | | | | | | | | | | NON INJECTED CLONE | | | | | | | | | | | | | | |
|---|---|---|---|---|---|---|---|---|---|---|---|---|---|---|---|---|---|---|---|---|---|---|---|---|---|---|---|---|---|---|---|---|---|
| | | | | Nuclear abnormality | OUTER CELLS | | | | | | | INNER CELLS | | | | | | | Nuclear abnormality | OUTER CELLS | | | | | | | INNER CELLS | | | | | | |
| | | | | | | GATA4+VE | GATA4-VE | SOX2+VE | SOX2-VE | GATA4 & SOX2 -VE | Apoptotic ♣ | | GATA4+VE | GATA4-VE | SOX2+VE | SOX2-VE | GATA4 & SOX2 -VE | Apoptotic ♣ | | | GATA4+VE | GATA4-VE | SOX2+VE | SOX2-VE | GATA4 & SOX2 -VE | Apoptotic ♣ | | GATA4+VE | GATA4-VE | SOX2+VE | SOX2-VE | GATA4 & SOX2 -VE | Apoptotic ♣ |
| 1 | 64 | 26 | 38 | 2 | 22 | 0 | 22 | 0 | 22 | 22 | 2 | 4 | 1 | 3 | 1 | 3 | 2 | 3 | 0 | 27 | 0 | 27 | 0 | 27 | 27 | 1 | 11 | 2 | 9 | 9 | 2 | 0 | 2 |
| 2 | 66 | 22 | 44 | 0 | 17 | 0 | 17 | 0 | 17 | 17 | 1 | 5 | 3 | 2 | 1 | 4 | 1 | 4 | 0 | 30 | 0 | 30 | 0 | 30 | 30 | 3 | 14 | 3 | 11 | 11 | 3 | 0 | 2 |
| 3 | 66 | 23 | 43 | 1 | 19 | 0 | 19 | 0 | 19 | 19 | 2 | 4 | 2 | 2 | 2 | 2 | 0 | 1 | 0 | 32 | 0 | 32 | 0 | 32 | 32 | 1 | 11 | 4 | 7 | 7 | 4 | 0 | 0 |
| 4 | 76 | 26 | 50 | 2 | 20 | 0 | 20 | 0 | 20 | 20 | 1 | 6 | 2 | 4 | 3 | 3 | 1 | 2 | 0 | 36 | 0 | 36 | 0 | 36 | 36 | 1 | 14 | 4 | 10 | 10 | 4 | 0 | 1 |
| 5 | 69 | 26 | 43 | 1 | 25 | 0 | 25 | 0 | 25 | 25 | 1 | 1 | 1 | 0 | 0 | 1 | 0 | 3 | 0 | 29 | 0 | 29 | 0 | 29 | 29 | 2 | 14 | 3 | 11 | 11 | 3 | 0 | 1 |
| 6 | 64 | 20 | 44 | 1 | 15 | 0 | 15 | 0 | 15 | 15 | 2 | 5 | 1 | 4 | 3 | 2 | 1 | 4 | 0 | 31 | 0 | 31 | 0 | 31 | 31 | 1 | 13 | 4 | 9 | 9 | 4 | 0 | 2 |
| 7 | 63 | 21 | 42 | 3 | 16 | 0 | 16 | 0 | 16 | 16 | 0 | 5 | 3 | 2 | 2 | 3 | 0 | 2 | 0 | 28 | 0 | 28 | 0 | 28 | 28 | 0 | 14 | 2 | 12 | 12 | 2 | 0 | 1 |
| 8 | 72 | 27 | 45 | 4 | 21 | 0 | 21 | 0 | 21 | 21 | 3 | 6 | 4 | 2 | 1 | 5 | 1 | 4 | 0 | 33 | 0 | 33 | 0 | 33 | 33 | 1 | 12 | 3 | 9 | 9 | 3 | 0 | 2 |
| 9 | 70 | 27 | 43 | 2 | 23 | 0 | 23 | 0 | 23 | 23 | 1 | 4 | 2 | 2 | 2 | 2 | 0 | 2 | 0 | 29 | 0 | 29 | 0 | 29 | 29 | 3 | 14 | 4 | 10 | 10 | 4 | 0 | 0 |
| 10 | 65 | 22 | 43 | 1 | 19 | 0 | 19 | 0 | 19 | 19 | 2 | 3 | 0 | 3 | 3 | 0 | 0 | 3 | 0 | 31 | 0 | 31 | 0 | 31 | 31 | 1 | 12 | 3 | 9 | 9 | 3 | 0 | 0 |
| 11 | 64 | 23 | 41 | 0 | 18 | 0 | 18 | 0 | 18 | 18 | 2 | 5 | 2 | 3 | 1 | 4 | 2 | 4 | 0 | 28 | 0 | 28 | 0 | 28 | 28 | 1 | 13 | 2 | 11 | 11 | 2 | 0 | 2 |
| 12 | 64 | 24 | 40 | 3 | 18 | 0 | 18 | 0 | 18 | 18 | 1 | 6 | 3 | 3 | 2 | 4 | 1 | 1 | 0 | 27 | 0 | 27 | 0 | 27 | 27 | 2 | 13 | 1 | 12 | 12 | 1 | 0 | 0 |
| 13 | 68 | 26 | 42 | 2 | 21 | 0 | 21 | 0 | 21 | 21 | 2 | 5 | 4 | 1 | 1 | 4 | 0 | 5 | 0 | 31 | 0 | 31 | 0 | 31 | 31 | 1 | 11 | 2 | 9 | 9 | 2 | 0 | 2 |
| 14 | 70 | 29 | 41 | 1 | 23 | 0 | 23 | 0 | 23 | 23 | 3 | 6 | 3 | 3 | 3 | 3 | 0 | 4 | 0 | 26 | 0 | 26 | 0 | 26 | 26 | 3 | 15 | 4 | 11 | 11 | 4 | 0 | 0 |
| 15 | 70 | 22 | 48 | 4 | 18 | 0 | 18 | 0 | 18 | 18 | 1 | 4 | 2 | 2 | 0 | 4 | 2 | 3 | 0 | 32 | 0 | 32 | 0 | 32 | 32 | 2 | 16 | 6 | 10 | 10 | 6 | 0 | 1 |
| 16 | 71 | 27 | 44 | 2 | 22 | 0 | 22 | 0 | 22 | 22 | 1 | 5 | 2 | 3 | 2 | 3 | 1 | 2 | 0 | 29 | 0 | 29 | 0 | 29 | 29 | 0 | 15 | 4 | 11 | 11 | 4 | 0 | 1 |
| 17 | 74 | 30 | 44 | 2 | 24 | 0 | 24 | 0 | 24 | 24 | 2 | 6 | 3 | 3 | 2 | 4 | 1 | 1 | 0 | 30 | 0 | 30 | 0 | 30 | 30 | 1 | 14 | 2 | 12 | 12 | 2 | 0 | 0 |
| 18 | 73 | 23 | 50 | 1 | 19 | 0 | 19 | 0 | 19 | 19 | 2 | 4 | 4 | 0 | 0 | 4 | 0 | 3 | 0 | 38 | 0 | 38 | 0 | 38 | 38 | 2 | 12 | 4 | 8 | 8 | 4 | 0 | 1 |
| 19 | 71 | 28 | 43 | 0 | 23 | 0 | 23 | 0 | 23 | 23 | 0 | 5 | 2 | 3 | 2 | 3 | 1 | 2 | 0 | 28 | 0 | 28 | 0 | 28 | 28 | 1 | 15 | 3 | 12 | 12 | 3 | 0 | 2 |
| 20 | 73 | 26 | 47 | 2 | 21 | 0 | 21 | 0 | 21 | 21 | 3 | 5 | 4 | 1 | 1 | 4 | 0 | 0 | 0 | 33 | 0 | 33 | 0 | 33 | 33 | 3 | 14 | 5 | 9 | 9 | 5 | 0 | 0 |
| TOTAL | 1373 | 498 | 875 | 34 | 404 | 0 | 404 | 0 | 404 | 404 | 32 | 94 | 48 | 46 | 32 | 62 | 14 | 53 | 0 | 608 | 0 | 608 | 0 | 608 | 608 | 30 | 267 | 65 | 202 | 202 | 65 | 0 | 20 |
| AVERAGE | 68.7 | 24.9 | 43.8 | 1.7 | 20.2 | 0.0 | 20.2 | 0.0 | 20.2 | 20.2 | 1.6 | 4.7 | 2.4 | 2.3 | 1.6 | 3.1 | 0.7 | 2.7 | 0.0 | 30.4 | 0.0 | 30.4 | 0.0 | 30.4 | 30.4 | 1.5 | 13.4 | 3.3 | 10.1 | 10.1 | 3.3 | 0.0 | 1.0 |
| SEM | 0.9 | 0.6 | 0.7 | 0.3 | 0.6 | 0.0 | 0.6 | 0.0 | 0.6 | 0.6 | 0.2 | 0.3 | 0.3 | 0.3 | 0.2 | 0.3 | 0.2 | 0.3 | 0.0 | 0.7 | 0.0 | 0.7 | 0.0 | 0.7 | 0.7 | 0.2 | 0.3 | 0.3 | 0.3 | 0.3 | 0.3 | 0.0 | 0.2 |
| Stat. sig. (inter-clone) ‡p<0.05, ‡‡p<0.005 | | ‡‡ | | ‡‡ | ‡‡ | | ‡‡ | | ‡‡ | ‡‡ | | ‡‡ | ‡ | ‡‡ | ‡‡ | | ‡‡ | ‡‡ | | | | | | | | | | | | | | | |
| p-value (2-tailed students t-test) | | 5.35E-22 | | 1.28E-07 | 1.52E-13 | 1.00E+00 | 1.52E-13 | 1.00E+00 | 1.52E-13 | 1.52E-13 | 7.31E-01 | 5.05E-22 | 2.79E-02 | 5.28E-21 | 5.67E-23 | 6.97E-01 | 1.25E-04 | 3.23E-05 | | | | | | | | | | | | | | | |
| Stat. sig. (exp. vs. con embryo) *p<0.05, **p<0.005 | ** | ** | | ** | ** | | ** | | ** | ** | | ** | | ** | ** | | ** | ** | | | | | | | | | ** | | ** | ** | | | |
| p-value (2-tailed students t-test) | 8.29E-13 | 3.96E-20 | 2.92E-01 | 1.69E-06 | 1.64E-12 | 1.00E+00 | 1.64E-12 | 1.00E+00 | 1.64E-12 | 1.64E-12 | 4.35E-01 | 6.86E-16 | 5.45E-02 | 7.03E-17 | 1.06E-18 | 8.31E-01 | 5.56E-04 | 3.67E-05 | 1.00E+00 | 5.08E-01 | 1.00E+00 | 5.08E-01 | 1.00E+00 | 5.08E-01 | 5.08E-01 | 5.59E-01 | 2.76E-03 | 8.79E-01 | 2.41E-03 | 2.41E-03 | 8.79E-01 | 1.00E+00 | 5.45E-01 |

**Supplementary Tables ST24**

#### Control siRNA (10μM), 1in2 micro-injection, 32-cell (E3.5) stage

| # | Total cell number per embryo | Cells in injected clone | Cells in non-injected clone | INJECTED CLONE: Nuclear abnormality | INJECTED CLONE: OUTER CELLS | INJECTED CLONE: INNER CELLS | NON INJECTED CLONE: Nuclear abnormality | NON INJECTED CLONE: OUTER CELLS | NON INJECTED CLONE: INNER CELLS |
|---|---|---|---|---|---|---|---|---|---|
| 1 | 38 | 19 | 19 | 0 | 12 | 7 | 0 | 12 | 7 |
| 2 | 31 | 14 | 17 | 0 | 8 | 6 | 0 | 8 | 9 |
| 3 | 35 | 17 | 18 | 0 | 11 | 6 | 0 | 12 | 6 |
| 4 | 37 | 19 | 18 | 0 | 12 | 7 | 0 | 12 | 6 |
| 5 | 31 | 14 | 17 | 0 | 8 | 6 | 0 | 8 | 9 |
| 6 | 32 | 15 | 17 | 0 | 10 | 5 | 0 | 10 | 7 |
| 7 | 32 | 18 | 14 | 0 | 11 | 7 | 0 | 9 | 5 |
| 8 | 36 | 19 | 17 | 0 | 11 | 8 | 0 | 11 | 6 |
| 9 | 31 | 15 | 16 | 0 | 9 | 6 | 0 | 9 | 7 |
| 10 | 34 | 16 | 18 | 0 | 10 | 6 | 0 | 10 | 8 |
| 11 | 37 | 19 | 18 | 0 | 11 | 8 | 0 | 11 | 7 |
| 12 | 31 | 13 | 18 | 0 | 10 | 3 | 0 | 12 | 6 |
| 13 | 39 | 20 | 19 | 0 | 13 | 7 | 0 | 12 | 7 |
| 14 | 35 | 17 | 18 | 0 | 11 | 6 | 0 | 11 | 7 |
| 15 | 35 | 17 | 18 | 0 | 11 | 6 | 0 | 12 | 6 |
| **TOTAL** | **514** | **252** | **262** | **0** | **158** | **94** | **0** | **159** | **103** |
| **AVERAGE** | **34.3** | **16.8** | **17.5** | **0.0** | **10.5** | **6.3** | **0.0** | **10.6** | **6.9** |
| SEM | **0.7** | **0.6** | **0.3** | **0.0** | **0.4** | **0.3** | **0.0** | **0.4** | **0.3** |
| Stat. sig. (inter-clone) ‡p<0.05, ‡‡p<0.005 | | | | | | | | | |
| p-value (2-tailed students t-test) | | 3.18E-01 | | 1.00E+00 | 9.01E-01 | 1.73E-01 | | | |

#### Wwc2-specific siRNA (10μM), 1in2 micro-injection, 32-cell (E3.5) stage

| # | Total cell number per embryo | Cells in injected clone | Cells in non-injected clone | INJECTED CLONE: Nuclear abnormality | INJECTED CLONE: OUTER CELLS | INJECTED CLONE: INNER CELLS | NON INJECTED CLONE: Nuclear abnormality | NON INJECTED CLONE: OUTER CELLS | NON INJECTED CLONE: INNER CELLS |
|---|---|---|---|---|---|---|---|---|---|
| 1 | 38 | 16 | 22 | 2 | 12 | 4 | 0 | 15 | 7 |
| 2 | 33 | 13 | 20 | 1 | 11 | 2 | 0 | 12 | 8 |
| 3 | 28 | 11 | 17 | 0 | 9 | 2 | 0 | 11 | 6 |
| 4 | 33 | 14 | 19 | 3 | 11 | 3 | 0 | 12 | 7 |
| 5 | 38 | 16 | 22 | 0 | 12 | 4 | 0 | 13 | 9 |
| 6 | 29 | 11 | 18 | 1 | 11 | 0 | 0 | 11 | 7 |
| 7 | 28 | 13 | 15 | 0 | 10 | 3 | 0 | 10 | 5 |
| 8 | 28 | 11 | 17 | 1 | 8 | 3 | 0 | 11 | 6 |
| 9 | 29 | 11 | 18 | 0 | 10 | 1 | 0 | 10 | 8 |
| 10 | 25 | 9 | 16 | 2 | 9 | 0 | 0 | 11 | 5 |
| 11 | 32 | 14 | 18 | 1 | 12 | 2 | 0 | 11 | 7 |
| 12 | 32 | 13 | 19 | 1 | 10 | 3 | 0 | 13 | 6 |
| 13 | 30 | 11 | 19 | 0 | 10 | 1 | 0 | 12 | 7 |
| 14 | 40 | 16 | 24 | 2 | 12 | 4 | 0 | 16 | 8 |
| 15 | 31 | 13 | 18 | 0 | 12 | 1 | 0 | 12 | 6 |
| 16 | 39 | 17 | 22 | 1 | 14 | 3 | 0 | 14 | 8 |
| 17 | 31 | 12 | 19 | 0 | 12 | 0 | 0 | 12 | 7 |
| 18 | 34 | 13 | 21 | 1 | 9 | 4 | 0 | 14 | 7 |
| 19 | 28 | 10 | 18 | 1 | 9 | 1 | 0 | 12 | 6 |
| **TOTAL** | **606** | **244** | **362** | **17** | **203** | **41** | **0** | **232** | **130** |
| **AVERAGE** | **31.9** | **12.8** | **19.1** | **0.9** | **10.7** | **2.2** | **0.0** | **12.2** | **6.8** |
| SEM | **1.0** | **0.5** | **0.5** | **0.2** | **0.4** | **0.3** | **0.0** | **0.4** | **0.2** |
| Stat. sig. (inter-clone) ‡p<0.05, ‡‡p<0.005 | | ‡‡ | | ‡‡ | ‡ | ‡‡ | | | |
| p-value (2-tailed students t-test) | | 4.78E-10 | | 7.81E-05 | 5.04E-03 | 1.41E-13 | | | |
| Stat. sig. (exp. vs. con embryo) *p<0.05, **p<0.005 | | ** | * | ** | | ** | | * | |
| p-value (2-tailed students t-test) | 7.23E-02 | 1.32E-05 | 2.21E-02 | 4.07E-04 | 7.69E-01 | 3.84E-10 | 1.00E+00 | 5.58E-03 | 9.49E-01 |

#### Wwc2-specific siRNA (10μM) +HA-Wwc2 (siRNA resistant) mRNA (80ng/μL), 1in2 micro-injection, 32-cell (E3.5) stage

| # | Total cell number per embryo | Cells in injected clone | Cells in non-injected clone | INJECTED CLONE: Nuclear abnormality | INJECTED CLONE: OUTER CELLS | INJECTED CLONE: INNER CELLS | NON INJECTED CLONE: Nuclear abnormality | NON INJECTED CLONE: OUTER CELLS | NON INJECTED CLONE: INNER CELLS |
|---|---|---|---|---|---|---|---|---|---|
| 1 | 35 | 18 | 17 | 0 | 12 | 6 | 0 | 10 | 7 |
| 2 | 32 | 16 | 16 | 0 | 10 | 6 | 0 | 11 | 5 |
| 3 | 32 | 17 | 15 | 0 | 10 | 7 | 0 | 9 | 6 |
| 4 | 33 | 16 | 17 | 1 | 12 | 4 | 0 | 9 | 8 |
| 5 | 35 | 18 | 17 | 0 | 11 | 7 | 0 | 10 | 7 |
| 6 | 32 | 15 | 17 | 0 | 11 | 4 | 0 | 8 | 9 |
| 7 | 37 | 18 | 19 | 0 | 10 | 8 | 0 | 15 | 4 |
| 8 | 29 | 14 | 15 | 0 | 9 | 5 | 0 | 11 | 4 |
| 9 | 33 | 16 | 17 | 1 | 11 | 5 | 0 | 10 | 7 |
| 10 | 32 | 17 | 15 | 0 | 10 | 7 | 0 | 9 | 6 |
| 11 | 37 | 20 | 17 | 0 | 12 | 8 | 0 | 11 | 6 |
| 12 | 33 | 16 | 17 | 0 | 10 | 6 | 0 | 11 | 6 |
| 13 | 34 | 17 | 17 | 0 | 11 | 6 | 0 | 9 | 8 |
| 14 | 42 | 21 | 21 | 0 | 13 | 8 | 0 | 14 | 7 |
| 15 | 32 | 16 | 16 | 0 | 10 | 6 | 0 | 11 | 5 |
| 16 | 39 | 19 | 20 | 0 | 13 | 6 | 0 | 12 | 8 |
| 17 | 30 | 14 | 16 | 1 | 9 | 5 | 0 | 9 | 7 |
| 18 | 37 | 19 | 18 | 0 | 12 | 7 | 0 | 11 | 7 |
| 19 | 37 | 18 | 19 | 0 | 11 | 7 | 0 | 13 | 6 |
| 20 | 35 | 17 | 18 | 0 | 10 | 7 | 0 | 12 | 6 |
| 21 | 36 | 17 | 19 | 0 | 11 | 6 | 0 | 12 | 7 |
| 22 | 36 | 19 | 17 | 0 | 13 | 6 | 0 | 11 | 6 |
| **TOTAL** | **758** | **378** | **380** | **3** | **241** | **137** | **0** | **238** | **142** |
| **AVERAGE** | **34.5** | **17.2** | **17.3** | **0.1** | **11.0** | **6.2** | **0.0** | **10.8** | **6.5** |
| SEM | **0.7** | **0.4** | **0.3** | **0.1** | **0.3** | **0.2** | **0.0** | **0.4** | **0.3** |
| Stat. sig. (inter-clone) ‡p<0.05, ‡‡p<0.005 | | | | | | | | | |
| p-value (2-tailed students t-test) | | 8.59E-01 | | 7.57E-02 | 7.64E-01 | 5.36E-01 | | | |
| Stat. sig. (exp. vs. con embryo) *p<0.05, **p<0.005 | | | | | | | | | |
| p-value (2-tailed students t-test) | 8.50E-01 | 5.66E-01 | 6.93E-01 | 1.43E-01 | 3.38E-01 | 9.21E-01 | 1.00E+00 | 6.95E-01 | 3.16E-01 |
| Stat. sig. (exp. vs. exp/KD embryo) §p<0.05, §§p<0.005 | § | §§ | §§ | §§ | | §§ | | § | |
| p-value (2-tailed students t-test) | 3.15E-02 | 3.05E-08 | 5.73E-03 | 5.97E-04 | 5.32E-01 | 1.86E-12 | 1.00E+00 | 1.18E-02 | 2.99E-01 |

**Supplementary Tables ST25**

| Control siRNA (10µM), 1in2 micro-injection, >64-cell (E4.5) stage | | | | | | | | | |
|---|---|---|---|---|---|---|---|---|---|
| # | Total cell number per embryo | Cells in injected clone | Cells in non-injected clone | INJECTED CLONE | | | NON INJECTED CLONE | | |
| | | | | Nuclear abnormality | OUTER CELLS | INNER CELLS | Nuclear abnormality | OUTER CELLS | INNER CELLS |
| **1** | 74 | 36 | 38 | 0 | 26 | 10 | 0 | 29 | 9 |
| **2** | 72 | 34 | 38 | 0 | 25 | 9 | 0 | 28 | 10 |
| **3** | 76 | 39 | 37 | 0 | 29 | 10 | 0 | 28 | 9 |
| **4** | 79 | 39 | 40 | 0 | 28 | 11 | 0 | 30 | 10 |
| **5** | 73 | 36 | 37 | 0 | 27 | 9 | 0 | 29 | 8 |
| **6** | 80 | 39 | 41 | 0 | 29 | 10 | 0 | 32 | 9 |
| **7** | 73 | 36 | 37 | 0 | 28 | 8 | 0 | 28 | 9 |
| **8** | 79 | 40 | 39 | 0 | 28 | 12 | 0 | 29 | 10 |
| **9** | 78 | 40 | 38 | 0 | 31 | 9 | 0 | 26 | 12 |
| **10** | 82 | 42 | 40 | 0 | 31 | 11 | 0 | 30 | 10 |
| **11** | 78 | 38 | 40 | 0 | 28 | 10 | 0 | 28 | 12 |
| **12** | 81 | 41 | 40 | 0 | 30 | 11 | 0 | 31 | 9 |
| **13** | 74 | 37 | 37 | 0 | 29 | 8 | 0 | 30 | 7 |
| **TOTAL** | **999** | **497** | **502** | **0** | **369** | **128** | **0** | **378** | **124** |
| **AVERAGE** | **76.8** | **38.2** | **38.6** | **0.0** | **28.4** | **9.8** | **0.0** | **29.1** | **9.5** |
| SEM | **0.9** | **0.6** | **0.4** | **0.0** | **0.5** | **0.3** | **0.0** | **0.4** | **0.4** |
| Stat. sig. (inter-clone) ‡p<0.05, ‡‡p<0.005 | | | | | | | | | |
| p-value (2-tailed students t-test) | | 6.16E-01 | | 1.00E+00 | 2.98E-01 | 5.54E-01 | | | |

| Wwc2-specific siRNA (10µM), 1in2 micro-injection, >64-cell (E4.5) stage | | | | | | | | | |
|---|---|---|---|---|---|---|---|---|---|
| # | Total cell number per embryo | Cells in injected clone | Cells in non-injected clone | INJECTED CLONE | | | NON INJECTED CLONE | | |
| | | | | Nuclear abnormality | OUTER CELLS | INNER CELLS | Nuclear abnormality | OUTER CELLS | INNER CELLS |
| **1** | 54 | 18 | 36 | 2 | 17 | 1 | 0 | 27 | 9 |
| **2** | 64 | 23 | 41 | 1 | 21 | 2 | 0 | 30 | 11 |
| **3** | 73 | 26 | 47 | 2 | 24 | 2 | 0 | 35 | 12 |
| **4** | 57 | 19 | 38 | 1 | 19 | 0 | 0 | 28 | 10 |
| **5** | 57 | 21 | 36 | 1 | 20 | 1 | 0 | 27 | 9 |
| **6** | 68 | 27 | 41 | 3 | 25 | 2 | 0 | 30 | 11 |
| **7** | 66 | 25 | 41 | 0 | 24 | 1 | 0 | 31 | 10 |
| **8** | 54 | 18 | 36 | 2 | 18 | 0 | 0 | 28 | 8 |
| **9** | 61 | 21 | 40 | 0 | 19 | 2 | 0 | 29 | 11 |
| **10** | 60 | 23 | 37 | 2 | 22 | 1 | 0 | 27 | 10 |
| **11** | 72 | 29 | 43 | 4 | 26 | 3 | 0 | 36 | 7 |
| **12** | 64 | 21 | 43 | 1 | 21 | 0 | 0 | 33 | 10 |
| **13** | 68 | 25 | 43 | 2 | 23 | 2 | 0 | 31 | 12 |
| **TOTAL** | **818** | **296** | **522** | **21** | **279** | **17** | **0** | **392** | **130** |
| **AVERAGE** | **62.9** | **22.8** | **40.2** | **1.6** | **21.5** | **1.3** | **0.0** | **30.2** | **10.0** |
| SEM | **1.8** | **1.0** | **0.9** | **0.3** | **0.8** | **0.3** | **0.0** | **0.8** | **0.4** |
| Stat. sig. (inter-clone) ‡p<0.05, ‡‡p<0.005 | | ‡‡ | | ‡‡ | ‡‡ | ‡‡ | | | |
| p-value (2-tailed students t-test) | | 3.04E-12 | | 2.53E-05 | 7.36E-08 | 2.17E-15 | | | |
| Stat. sig. (exp. vs. con embryo) *p<0.05, **p<0.005 | ** | ** | | ** | ** | ** | | | |
| p-value (2-tailed students t-test) | 3.40E-07 | 1.43E-12 | 1.47E-01 | 2.53E-05 | 9.32E-08 | 1.81E-16 | 1.00E+00 | 2.61E-01 | 4.19E-01 |

| Wwc2-specific siRNA (10µM) +HA-Wwc2 (siRNA resistant) mRNA (80ng/µL), 1in2 micro-injection, >64-cell (E4.5) stage | | | | | | | | | |
|---|---|---|---|---|---|---|---|---|---|
| # | Total cell number per embryo | Cells in injected clone | Cells in non-injected clone | INJECTED CLONE | | | NON INJECTED CLONE | | |
| | | | | Nuclear abnormality | OUTER CELLS | INNER CELLS | Nuclear abnormality | OUTER CELLS | INNER CELLS |
| **1** | 57 | 28 | 29 | 1 | 20 | 8 | 0 | 22 | 7 |
| **2** | 83 | 40 | 43 | 0 | 25 | 15 | 0 | 36 | 7 |
| **3** | 60 | 29 | 31 | 0 | 21 | 8 | 0 | 23 | 8 |
| **4** | 83 | 38 | 45 | 1 | 24 | 14 | 0 | 35 | 10 |
| **5** | 79 | 38 | 41 | 0 | 27 | 11 | 0 | 32 | 9 |
| **6** | 71 | 35 | 36 | 0 | 26 | 9 | 0 | 29 | 7 |
| **7** | 89 | 42 | 47 | 0 | 30 | 12 | 0 | 37 | 10 |
| **8** | 65 | 33 | 32 | 0 | 23 | 10 | 0 | 24 | 8 |
| **9** | 55 | 27 | 28 | 1 | 19 | 8 | 0 | 22 | 6 |
| **10** | 80 | 44 | 36 | 0 | 34 | 10 | 0 | 28 | 8 |
| **11** | 71 | 34 | 37 | 0 | 25 | 9 | 0 | 27 | 10 |
| **12** | 82 | 40 | 42 | 0 | 28 | 12 | 0 | 34 | 8 |
| **13** | 75 | 38 | 37 | 0 | 29 | 9 | 0 | 30 | 7 |
| **14** | 74 | 37 | 37 | 0 | 25 | 12 | 0 | 27 | 10 |
| **15** | 79 | 39 | 40 | 0 | 32 | 7 | 0 | 31 | 9 |
| **16** | 74 | 37 | 37 | 0 | 27 | 10 | 0 | 30 | 7 |
| **17** | 69 | 35 | 34 | 0 | 28 | 7 | 0 | 25 | 9 |
| **18** | 76 | 42 | 34 | 0 | 32 | 10 | 0 | 26 | 8 |
| **TOTAL** | **1322** | **656** | **666** | **3** | **475** | **181** | **0** | **518** | **148** |
| **AVERAGE** | **73.4** | **36.4** | **37.0** | **0.2** | **26.4** | **10.1** | **0.0** | **28.8** | **8.2** |
| SEM | **2.2** | **1.1** | **1.2** | **0.1** | **1.0** | **0.5** | **0.0** | **1.1** | **0.3** |
| Stat. sig. (inter-clone) ‡p<0.05, ‡‡p<0.005 | | | | | | ‡‡ | | | |
| p-value (2-tailed students t-test) | | 7.44E-01 | | 7.39E-02 | 1.17E-01 | 4.99E-03 | | | |
| Stat. sig. (exp. vs. con embryo) *p<0.05, **p<0.005 | | | | | | | | | * |
| p-value (2-tailed students t-test) | 2.24E-01 | 2.28E-01 | 2.94E-01 | 1.30E-01 | 1.16E-01 | 7.64E-01 | 1.00E+00 | 8.29E-01 | 1.03E-02 |
| Stat. sig. (exp. vs. exp/KD embryo) §p<0.05, §§p<0.005 | §§ | §§ | §§ | §§ | §§ | §§ | | | §§ |
| p-value (2-tailed students t-test) | 1.59E-03 | 1.46E-09 | 7.01E-02 | 1.86E-05 | 9.12E-04 | 1.06E-13 | 1.00E+00 | 3.65E-01 | 1.14E-03 |

**Supplementary Tables ST26**

#### Control siRNA (10µM), 2in2 micro-injection, 32-cell (E3.5) stage

| # | Total cell number per embryo | Total midbody number per embryo | Midbodies per total cell number | OUTER CELLS | | | INNER CELLS | | |
|---|---|---|---|---|---|---|---|---|---|
| | | | | Cell number | OUTER midbody | Midbodies per OUTER cell | Cell number | INNER midbody | Midbodies per INNER cell |
| **1** | 33 | 12 | 0.36 | 21 | 9 | 0.43 | 12 | 3 | 0.25 |
| **2** | 38 | 7 | 0.18 | 24 | 7 | 0.29 | 14 | 0 | 0.00 |
| **3** | 39 | 10 | 0.26 | 26 | 8 | 0.31 | 13 | 2 | 0.15 |
| **4** | 43 | 17 | 0.40 | 28 | 11 | 0.39 | 15 | 6 | 0.40 |
| **5** | 33 | 14 | 0.42 | 23 | 10 | 0.43 | 10 | 4 | 0.40 |
| **6** | 31 | 12 | 0.39 | 22 | 9 | 0.41 | 9 | 3 | 0.33 |
| **7** | 41 | 13 | 0.32 | 26 | 8 | 0.31 | 15 | 5 | 0.33 |
| **8** | 35 | 13 | 0.37 | 24 | 9 | 0.38 | 11 | 4 | 0.36 |
| **9** | 37 | 12 | 0.32 | 25 | 7 | 0.28 | 12 | 5 | 0.42 |
| **10** | 33 | 10 | 0.30 | 23 | 8 | 0.35 | 10 | 2 | 0.20 |
| **11** | 40 | 8 | 0.20 | 26 | 7 | 0.27 | 14 | 1 | 0.07 |
| **12** | 31 | 6 | 0.19 | 22 | 6 | 0.27 | 9 | 0 | 0.00 |
| **13** | 34 | 11 | 0.32 | 23 | 8 | 0.35 | 11 | 3 | 0.27 |
| **14** | 33 | 14 | 0.42 | 23 | 10 | 0.43 | 10 | 4 | 0.40 |
| **15** | 37 | 12 | 0.32 | 24 | 9 | 0.38 | 13 | 3 | 0.23 |
| **16** | 36 | 8 | 0.22 | 25 | 6 | 0.24 | 11 | 2 | 0.18 |
| **17** | 35 | 10 | 0.29 | 24 | 10 | 0.42 | 11 | 0 | 0.00 |
| **18** | 33 | 7 | 0.21 | 23 | 6 | 0.26 | 10 | 1 | 0.10 |
| **TOTAL** | **642** | **196** | **5.51** | **432** | **148** | **6.19** | **210** | **48** | **4.11** |
| **AVERAGE** | **35.7** | **10.9** | **0.31** | **24.0** | **8.2** | **0.34** | **11.7** | **2.7** | **0.23** |
| SEM | **0.8** | **0.7** | **0.02** | **0.4** | **0.4** | **0.02** | **0.5** | **0.4** | **0.03** |

#### Wwc2-specific siRNA (10µM), 2in2 micro-injection, 32-cell (E3.5) stage

| # | Total cell number per embryo | Total midbody number per embryo | Midbodies per total cell number | OUTER CELLS | | | INNER CELLS | | |
|---|---|---|---|---|---|---|---|---|---|
| | | | | Cell number | OUTER midbody | Midbodies per OUTER cell | Cell number | INNER midbody | Midbodies per INNER cell |
| **1** | 16 | 2 | 0.13 | 14 | 2 | 0.14 | 2 | 0 | 0.00 |
| **2** | 16 | 5 | 0.31 | 15 | 4 | 0.27 | 1 | 1 | 1.00 |
| **3** | 13 | 4 | 0.31 | 12 | 4 | 0.33 | 1 | 0 | 0.00 |
| **4** | 24 | 5 | 0.21 | 19 | 5 | 0.26 | 5 | 0 | 0.00 |
| **5** | 19 | 6 | 0.32 | 17 | 6 | 0.35 | 2 | 0 | 0.00 |
| **6** | 18 | 7 | 0.39 | 16 | 7 | 0.44 | 2 | 0 | 0.00 |
| **7** | 25 | 5 | 0.20 | 20 | 4 | 0.20 | 5 | 1 | 0.20 |
| **8** | 18 | 5 | 0.28 | 15 | 5 | 0.33 | 3 | 0 | 0.00 |
| **9** | 9 | 3 | 0.33 | 9 | 3 | 0.33 | 0 | 0 | 0.00 |
| **10** | 15 | 5 | 0.33 | 14 | 5 | 0.36 | 1 | 0 | 0.00 |
| **11** | 16 | 5 | 0.31 | 13 | 5 | 0.38 | 3 | 0 | 0.00 |
| **12** | 15 | 4 | 0.27 | 14 | 4 | 0.29 | 1 | 0 | 0.00 |
| **13** | 14 | 4 | 0.29 | 12 | 4 | 0.33 | 2 | 0 | 0.00 |
| **14** | 11 | 2 | 0.18 | 11 | 2 | 0.18 | 0 | 0 | 0.00 |
| **15** | 17 | 3 | 0.18 | 16 | 3 | 0.19 | 1 | 0 | 0.00 |
| **16** | 16 | 4 | 0.25 | 15 | 3 | 0.20 | 1 | 1 | 1.00 |
| **17** | 14 | 2 | 0.14 | 13 | 2 | 0.15 | 1 | 0 | 0.00 |
| **18** | 14 | 2 | 0.14 | 13 | 2 | 0.15 | 1 | 0 | 0.00 |
| **TOTAL** | **290** | **73** | **4.56** | **258** | **70** | **4.90** | **32** | **3** | **2.20** |
| **AVERAGE** | **16.1** | **4.1** | **0.25** | **14.3** | **3.9** | **0.27** | **1.8** | **0.2** | **0.12** |
| SEM | **0.9** | **0.3** | **0.02** | **0.6** | **0.3** | **0.02** | **0.3** | **0.1** | **0.08** |
| Stat. sig. (inter-clone) ‡p<0.05, ‡‡p<0.005 | **‡‡** | **‡‡** | | **‡‡** | **‡‡** | **‡** | **‡‡** | **‡‡** | |
| p-value (2-tailed students t-test) | 2.68E-17 | 2.16E-10 | 5.29E-02 | 1.66E-14 | 3.09E-10 | 1.05E-02 | 1.75E-18 | 2.00E-06 | 2.14E-01 |
